# A green alternative to fragrant agarwood sesquiterpenoid production

**DOI:** 10.1101/2023.10.06.561217

**Authors:** Sergio Gutiérrez, Sebastian Overmans, Gordon B. Wellman, Vasilios G. Samaras, Claudia Oviedo, Martin Gede, Gyorgy Szekely, Kyle J. Lauersen

## Abstract

Certain endangered Thymelaeaceous trees are major sources of the fragrant and highly valued resinous agarwood, comprised of hundreds of oxygenated sesquiterpenoids (STPs). Despite growing pressure on natural agarwood sources, the chemical complexity of STPs severely limits synthetic production. Here, we catalogued the chemical diversity in 58 agarwood samples by two-dimensional gas chromatography–mass spectrometry and partially recreated complex STP mixtures through synthetic biology. We improved STP yields in the unicellular alga *Chlamydomonas reinhardtii* by combinatorial engineering to biosynthesise nine macrocyclic STP backbones found in agarwood. A bioprocess following green-chemistry principles was developed that exploits ‘milking’ of STPs without cell lysis, solvent–solvent STP extraction, solvent–STP nanofiltration, and bulk STP oxy-functionalisation to obtain terpene mixtures like those of agarwood. This process occurs with total solvent recycling and enables continuous production. Our synthetic-biology approach offers a sustainable alternative to harvesting agarwood trees to obtain mixtures of complex, fragrant, oxygenated STPs.

## Introduction

Agarwood is the common name given to the fragrant resinous woods derived from trees in the Thymeleaceae family, including *Aquilaria*, *Enkleia*, *Aetoxylon*, *Gonystylus*, *Wikstroemia*, and *Gyrinops*^1^. Indigenous to Southeast Asian countries, these trees are prized for the complex fragrances of their resins that originate from multiple oxygenated and non-oxygenated sesquiterpenoids^2–4^. These resins form after physical wounding of the tree, and their unique chemical profile develops over periods up to several decades^3,4^. The use of agarwood products for incense, perfumes, medicines and ornamentation has been recorded across cultures and centuries ^3–5^. Common applications are burning wood chips as incense and steam-distillate oils as perfume ingredients^4,6^. In particular, agarwood products were valued at $US 44 billion within the global perfume market^7^. Demand for agarwood has rapidly outpaced the supply of slow-growing natural sources, leading to the inclusion of 13 out of 20 *Aquilaria* species in the Appendix II of the Convention on International Trade in Endangered Species of Wild Fauna and Flora (CITES) as potentially threatened with extinction^8^. Despite global efforts to regulate trade and develop alternative production modes through silviculture or tissue culture, the clandestine sourcing of wild trees remains an issue that further pressures their existence^4,9^.

The chemical composition of agarwood is species-specific and often unique to individual plants, depending on certain abiotic factors^10,11^. The main fragrant compounds in agarwood are sesquiterpenoids (STPs), an ecologically important class of 15-carbon atom terpenoid compounds that play a crucial role in cloud formation^12^. These STPs originate from at least seven distinct primary chemical skeletons: agarofuran, agarospirane/vetispirane, cadinene, eremophilane/valencene, eudesmane/selinene, guaianes, and prezizanes^3,10,13^. Both abiotic and biotic factors govern their creation *in situ* where oxygenation of STPs to derivatives occurs, further shaping their individual scent profiles. From a metabolic-engineering standpoint, the tree itself serves only as the vehicle and matrix for chemical biosynthesis and resin accumulation, such that STP biosynthesis by metabolic engineering could be an alternative means for production of fragrant chemicals^4,14^. However, biochemical pathways leading to agarwood STPs are complex, poorly understood, and can be non-enzymatic in nature. So far, only guaiene STP synthases (STPSs) have been identified from *Aquilaria crassna* and *A. sinensis* and enzymatic routes to other primary STPs skeletons found in agarwood are not yet identified^11,14–16^.

Although the biochemical pathways to agarwood STPs are not elucidated, the diversity of natural product chemistry found in other organisms can provide alternative starting points to recreate agarwood STP chemical complexity. The primary carbon rings of agarofuran, cadinene, valencene, and selinane mirror STPs biosynthesised by valencene^17^, valerianol^18^, cadinol^19^, and aristolochene^20^ synthases, whereas zizaene synthase^21^ produces a prezizane-like STP. This existing repertoire of known STP synthases from plant and fungal species can produce STP mixtures that largely replicate the primary STP skeletons in agarwood.

Here, we aimed to reproduce some of the chemical complexity of agarwood STPs using synthetic biology and green bioprocessing. To inform the engineering approach, we first catalogued the chemical complexity of STPs in 58 agarwood samples using high-resolution, two-dimensional (2D) gas chromatography–time-of-flight mass spectrometry (GCxGC–TOF/MS). We then engineered unicellular algae to produce nine distinct STP chemical skeletons widely found in agarwood. A low-energy ‘microbial-milking’ technique was exploited to extract the STPs from growing algal cells, and energy-efficient solvent nanofiltration was used to concentrate the extracted STPs. Finally, we used green chemical reactions on these STP skeletons to generate a complex mixture of 103 oxygenated STPs typically found in agarwood. This process employs an autotrophic host and a low-energy, gentle STP-harvesting technique that enables solvent recycling, thus representing a practical alternative to harvesting agarwood trees to obtain their desirable, fragrant, terpenoid mixtures

## Results

### Chemical characterisation of agarwood and distillates

The *Aquilaria* sp. wood and distillate samples investigated here originate from various Southeast Asian countries (Fig. 1a, Supplementary Table 1). Wood samples (n = 36) exhibit a range of colours, with darker tones indicating higher resinous content (Fig. 1b), while the distillate oils (n = 22) exhibit varying amber-brown shades (Fig. 1c). Representative 2D GCxGC–TOF/MS analyses of wood (sample 34) and distillate (sample 39) are presented in Fig. 1d and e. Comprehensive data for all samples can be found in Supplementary File 1. Wood samples and distillate oils contain a combination of monoterpenes, sesquiterpenoids (STPs), functionalised sesquiterpenoids (fSTPs), and carboxylic acids. Each sample was unique in its chemical profile, but 298 STPs and fSTPs were common across agarwood samples and 338 STPs and fSTPs were common to all distillate samples (Supplementary Tables 2,3). Various STP skeleton types were found within the samples, including: agarofurans (α-agarofuran), agarospiranes (isoagarospirol), eudesmanes (isolongifolene, eudesmol, copaene, selinene, maaliene), eremophilanes (valencene, aristolochene, nootkatone, aristolene), guaianes (α-guaiene, β-guaiene, δ-guaiene, rotundone, viridiflorol, ledol, gurjunene, arodadendrene, guaiol, cedrol, patchoulene), cadinanes (τ-cadinol) and prezizanes (prezizaene, zizaene). Additional skeleton structures were identified, including patchoulol, caryophyllene, santalol, and bisabolol (Supplementary Tables 2–4, Supplementary Files 1–3).

**Figure 1.**
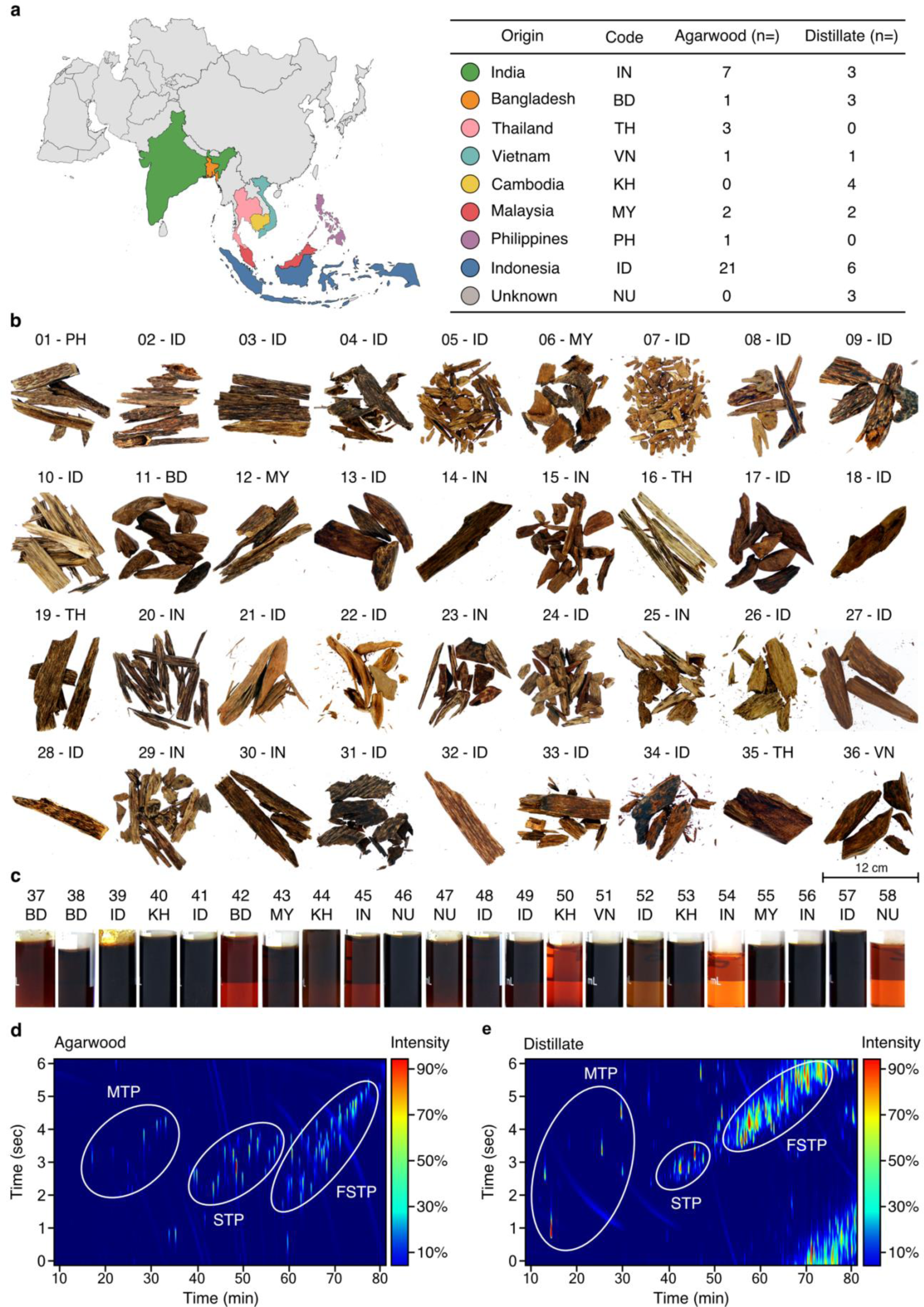
Comprehensive chemical characterisation of agarwood samples and distillates. **a.** Left: Map of Asia, colour-coded to represent different origins of agarwood samples. Right: Countries, their ISO 366 codes, and the total samples analysed from each, further categorised into agarwood wood chips ("bakhour") and steam distillate oils ("oudh"). **b,c.** Photographs of agarwood (b) and distillate samples (c). **d**. 2D chromatogram of wood-extract sample (34) with three primary chemical clusters circled. **e.** 2D chromatogram for distillate sample (39). MTP: monoterpenoids; STP: non-functionalised sesquiterpenoids; fSTP: functionalised sesquiterpenoids. 2D chromatograms for each sample shown are found in Supplementary Table 1. Supplementary Tables 2 and 3 provide summaries of common STPs identified in agarwoods and distillates, respectively. Supplementary Files 1–3 contain the supporting data for GC–TOF/MS, GC–MS/FID, and GCxGC–TOF/MS, respectively.

The identified sesquiterpenoid skeletons, spanning from the aromatic agarofurans to the complex prezizanes, offer a detailed insight into agarwood products and their chemistry. Compounds such as α-guaiene, δ-guaiene, and isolongifolene, expected from earlier research, were confirmed, underscoring the reliability of our analytical approach. Although, many compounds were identified across all agarwood and its distillates, each sample retained a distinct profile. These detailed observations enhance our comprehension of the natural chemical complexity of agarwood samples.

### Optimising algal metabolism to biosynthesise heterologous STPs

*Chlamydomonas reinhardtii* is an emerging host for metabolic engineering and heterologous terpene production^22,23^, as is known to have enhanced heterologous terpenoid yields following metabolic alterations^24–26^. These included the knockdown of native squalene synthase (*Cr*SQS-k.d.) increasing farnesyl pyrophosphate pools in the cytoplasm, leading to major improvements in sesquiterpenoid yields when either heterologous bisabolene or patchoulol sesquiterpenoid synthases were co-expressed^24,26^. Volatile engineered heterologous isoprene biosynthesis increased when carotenoid homeostasis was disrupted by introducing a beta-carotene ketolase to biosynthesise ketocarotenoids in the plastid^25^ (Fig. 2a, right panel). Here, the effects of these interventions alone or in combination were quantified in algae using the fragrant STP patchoulol as a reporter of biosynthetic activity (Fig. 2b).

**Figure 2.**
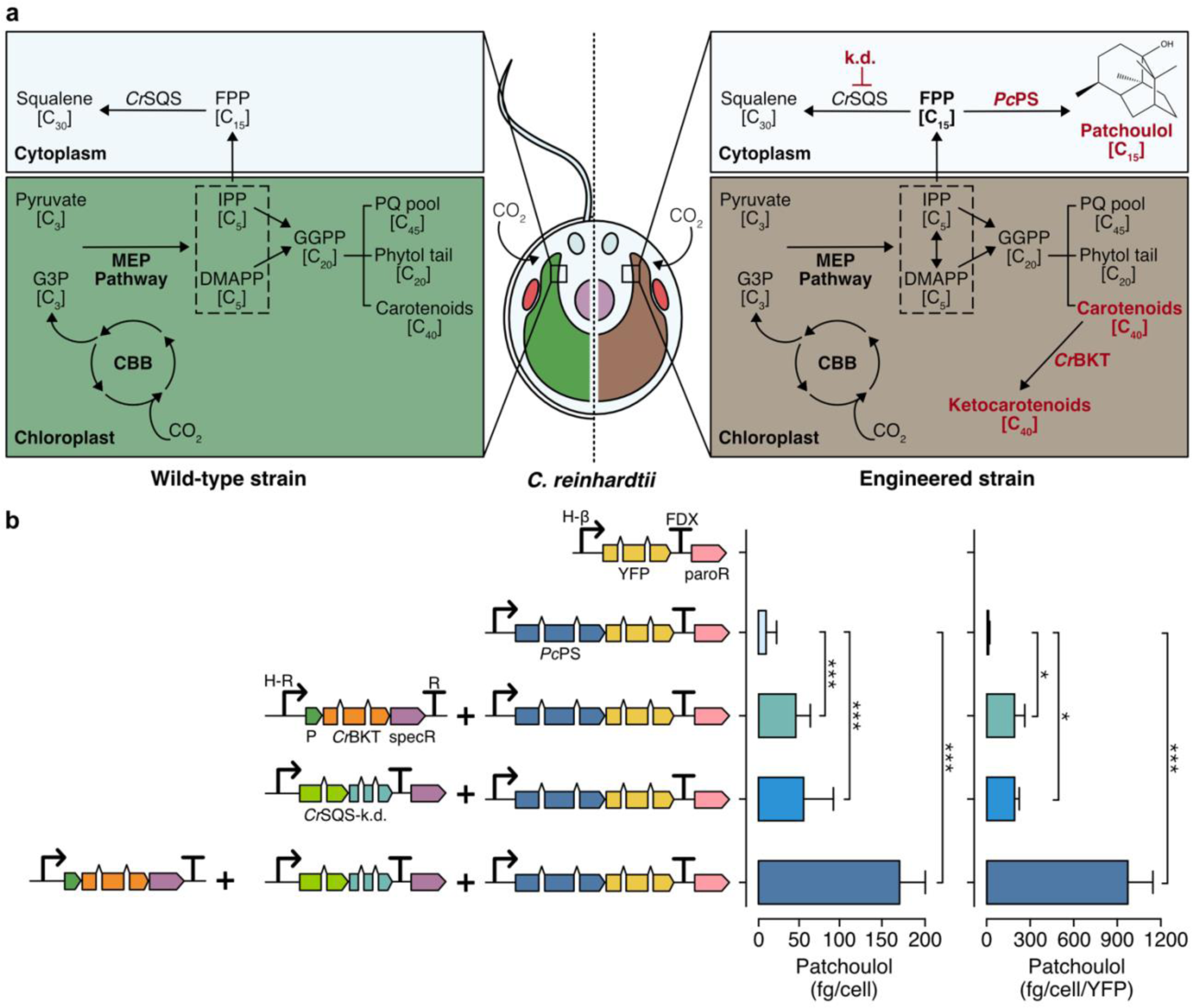
Native and engineered terpenoid pathway in *C. reinhardtii* for enhanced sesquiterpenoid biosynthesis. **a.** Left: Illustration of the endogenous terpenoid biosynthetic pathway in wild type *C. reinhardtii*. Right: Representation of an engineered *C. reinhardtii* strain characterised by absent flagella, cell-wall deficiency, and modifications of its terpenoid biosynthesis pathway known to improve heterologous STP production. Engineered changes are highlighted in red. **b.** Left: Depiction of algal-optimised transgenes including mVenus (yellow fluorescent protein, YFP), *Pc*PS–mVenus fusion, *Cr*BKT, and *Cr*SQS-k.d., transformed into *C. reinhardtii*. Right: Yield of patchoulol from each engineered construct combination. Abbreviations: k.d.: knockdown, *Pc*PS: *P. cablin* patchoulol synthase; *Cr*BKT: *C. reinhardtii* β-carotene ketolase; *Cr*SQS: *C. reinhardtii* squalene synthase amiRNA expression construct, FPP: farnesyl pyrophosphate, G3P: glyceraldehyde 3-phosphate, MEP: methylerythritol phosphate, CBB: Calvin-Benson-Bassham cycle, IPP: isopentenyl pyrophosphate, DMAPP: dimethylallyl pyrophosphate, PQ: plastoquinone, paroR: paromomycin resistance cassette, specR: spectinomycin resistance cassette, H-β: combined HSP70A-β tubulin promoter, FDX: ferredoxin 1 gene terminator, P: photosystem I reaction centre subunit II chloroplast-targeting peptide. Statistically significant differences between each combination were determined through ANOVA employing Turkey’s method for multiple comparisons. Statistical significance: \**p* < 0.05, \*\**p* < 0.01, \*\*\**p* < 0.001. Detailed calculations, cell concentrations and p-values are in Supplementary Table 4.

When expressed and localised in the algal cytoplasm, the patchoulol synthase from *Pogostemon cablin* (*Pc*PS) produced 5.4 ± 2.1 fg cell^−1^ of patchoulol. Biosynthesis of ketocarotenoids and perturbation of carotenoid biosynthesis through *Cr*BKT overexpression led to higher patchoulol levels of 41.2 ± 5.9 fg cell^−1^, representing an 8-fold increase over *Pc*PS expression alone. Suppressing endogenous *Cr*SQS through artificial micro-RNA in a strain expressing *Pc*PS further increased patchoulol yield to 71.7 ± 7.8 fg cell^−1^ (a 12-fold increase over a strain expressing *Pc*PS alone). The combination of *Cr*SQS-k.d. and *Cr*BKT overexpression in a strain expressing *Pc*PS gave patchoulol yields of 132.7 ± 41.1 fg cell^−1^ (2,215 µg L^−1^ culture, Supplementary Fig. 1, Supplementary Table 5). This strategy resulted in a 25-fold improvement in patchoulol yields over *Pc*PS alone and approximately a 3-fold enhancement over each individual strategy (Supplementary Table 5, one-way ANOVA (F(6, 2) = 158, *p* < 0.05). No statistically significant growth changes were detected in any strain grown photoheterotrophically with acetate (F(3, 20) = 0.035, *p* = 0.99) or photoautotrophically with CO_2_ (F(3, 20) = 0.265, *p* = 0.84) (Supplementary Fig. 2). A strain with both *Cr*BKT overexpression and *Cr*SQS-k.d. was used in all subsequent STP production experiments.

### Biosynthesis of diverse STPs in *C. reinhardtii*

Transformation and expression of 21 STPS isoforms in various fusions to the yellow fluorescent protein (YFP) reporter were performed, along with a YFP reporter alone control (23 plasmids in total, Fig. 3a, Supplementary Table 6). Due to non-homologous DNA integration in the algal nuclear genome, random position effects change expression dynamics across a population of transformants and large numbers of colonies must be screened to identify robust over-expressing transformants. Here, high-throughput robotics assisted handling was coupled to plate-level fluorescent protein expression screening for at least 384 transformants per plasmid. This screening allowed identification individual transformants for each target plasmids with robust STPS-FP fusion expression. The STPSs selected for expression were chosen based on their reported ability to make primary STPs skeletons commonly found in our analysis of agarwood samples. These STPSs biosynthesise nine distinct STP products (Fig. 3b). Santalene, zizaene, aristolochene, valencene, δ-guaiene, cadinol, valerianol, patchoulol, and bisabolol, were detectable as new GC–MS/FID peaks in dodecane overlays obtained from transgenic strains compared to overlays from control strains (Fig. 3b,e and Supplementary File 4). Each STPS isoform varied in the level of sesquiterpenoids they biosynthesised (Fig. 3b). Zizaene synthase had higher activity when YFP was placed at the N-terminal (plasmid 8, Fig. 3b), similar to reports of SUMO-tagged variants in *E. coli* ^21^.

**Figure 3.**
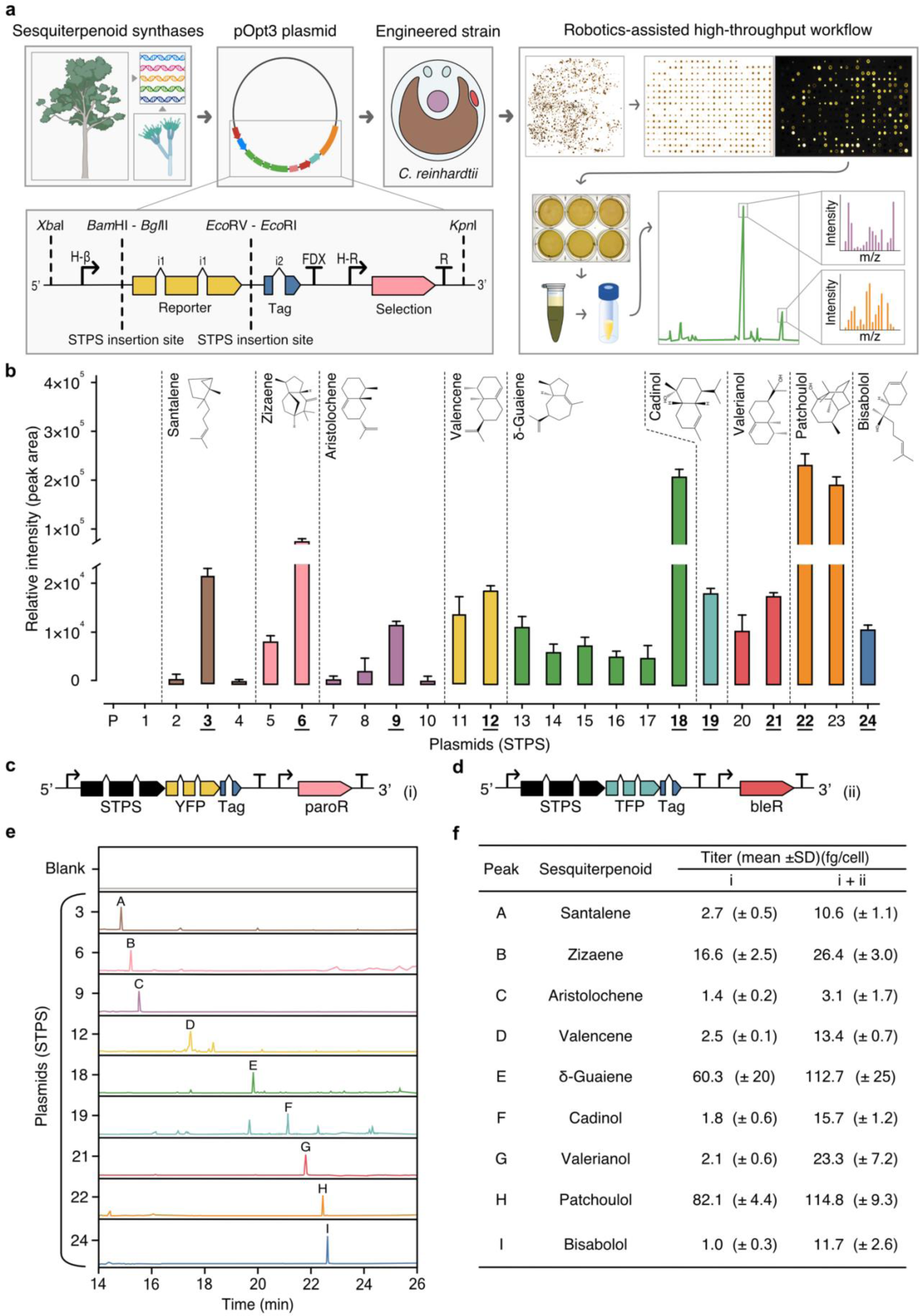
Biosynthesis of diverse STPs in *C. reinhardtii* through sesquiterpenoid synthase isoforms expression. **a.** The research workflow depicts the progression from selecting sesquiterpenoid synthases across various species for expression in *C. reinhardtii*. Isoform integration location into the pOpt3 in fusion to fluorescent reporter and robotics-assisted colony screening. Top-right box: Depicted is a transformant plate one week after transformation, organised transformant colonies and YFP signal indicating synthase–reporter fusion expression. Brightest colonies were screened for expression in replica-plate assays with dodecane solvent overlay, which could be separated from the culture by gravity and directly used for analysis in GC–MS/FID. **b.** Isoform performance was quantified from the most highly expressed transformants per STP (n = 25). Those chosen for further analysis are underlined **c.** An illustrated vector structure of the STPS–YFP with paromomycin resistance and (**d**) –mTFP1 with bleomycin resistance**. e.** GC–MS/FID chromatograms for each target STP. Annotated peaks include: A – santalene, B – zizaene, C – aristolochene, D – valencene, E – δ-guaiene, F – cadinol, G – valerianol, H – patchoulol, I – bisabolol. **f**. STP titres of single and double transformations. Abbreviations: YFP: mVenus yellow fluorescent protein; TFP: mTFP1 cyan fluorescent protein; paroR and bleR: paromomycin and bleomycin resistance, respectively; STPS: terpene synthase; Tag: strepII epitope. Chevrons i1/i2: intron addition to codon-optimised transgenes to enable strong algal expression. For a comprehensive list of plasmids, refer to Supplementary Table 5. Detailed calculations of productivity and cell concentrations are in Supplementary Tables 6–8. Supplementary Files 5 and 6 provide supporting data for GC– MS/FID and genetic constructs, respectively.

The optimal isoform with highest target-compound production was identified for each STPS (Fig. 3b). Subsequently, a second transformation was performed with the same STPS (Fig. 3c) fused to the monomeric teal (cyan) fluorescent protein 1 (mTFP1) reporter with a different selection marker (Fig. 3d). Secondary transformation was used to increase heterologous STPS enzyme titter in the cell to catalyse increased target STP biosynthesis. STP yield improvements were observed in all strains after secondary transformation (Fig. 3f), with an average 5.3-fold improvement in titre for each product (Supplementary Tables 7–9). Additional STP peaks were also identified in these extracts by GCxGC–TOF/MS, representing known side products of the individual synthases (Supplementary Table 10).

### A green bioprocess for functionalised STPs

To achieve sustainable production of complex STP mixtures present in agarwood, we developed a bioprocess design that minimised waste and could be operated at room temperature with only non-hazardous reagents (Fig. 4a). STP-producing algal cells were subjected to culture–solvent two-phase cultivations with a fluorocarbon underlay that allowed simple gravity separation of the fluorocarbon liquids containing STPs. Algal cultivation was followed by liquid–liquid extraction of STPs from the fluorocarbon solutions with ethanol, yielding individual STP–ethanol solutions, which were combined to form a STP mixture in ethanol (Fig. 4a,b). Low-energy, pressure-driven, organic-solvent nanofiltration (OSN) was then applied at room temperature, using only the pressure derived from a gas cylinder to drive ethanol through a nanofiltration membrane (Fig. 4a). OSN enabled concentration of STPs in ethanol by 53 times without evaporative loss (Fig. 4b).

**Figure 4.**
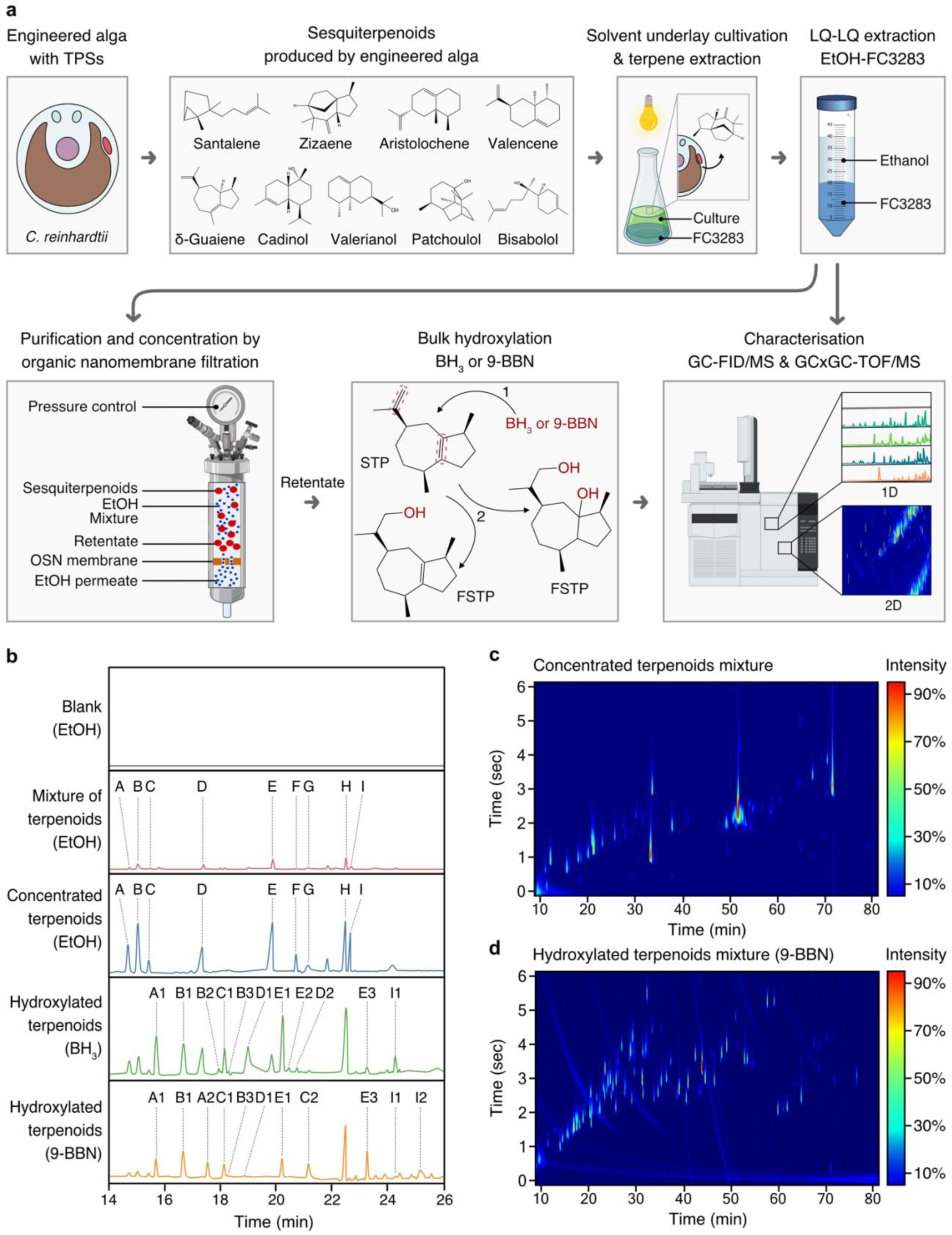
A green bioprocess workflow for the generation of agarwood-related sesquiterpenoids. **a.** STPs biosynthesised by engineered algae can be milked from the growing culture using heavy perfluorinated solvent (FC3283) underlays. STP-containing fluorocarbons can be separated from culture and STPs rapidly transferred to ethanol through liquid–liquid extraction from fluorocarbons. STPs can be concentrated in ethanol at room temperature in gas-pressure-driven nanofiltration cells wherein a nanoporous membrane retains STPs and allows permeation of ethanol. Concentrated STPs in ethanol can be subjected to hydroxylation reactions using BH_3_ or 9-BBN to increase molecular species diversity. Analysis throughout the process can be performed by GC–MS and GCxGC–TOF/MS. **b.** One-dimensional (1D) chromatograms of analysed process fractions, with STP skeletons labelled A to I and their respective derivatives after functionalisation: A: Santalene, A1: Santalol, A2: Bergamotene, B: Zizaene, B1: Cedrenol, B3: 12-nor-Preziza-7(15)-en-2-one, B3: Khusimol, C: Aristolochene, C1: Aristolone, C2: Dihydrokaranone, D: Valencene, D1: Nootkatol, D2: Nootkatone, E: δ-Guaiene, E1: Guaiol, E2: Globulol, E3: Guaia-1(10),11-dien-15,2-olide, F: Cadinol, G: Valerianol, H: Patchoulol, I: Bisabolol, I1: Bisabolol oxide B, I2: Bisabolol oxide A. **c.** Two-dimensional (2D) chromatogram depicting the starting mixed STPs. **d**. 2D chromatogram of STPs post-hydroxylation. Abbreviations: BH_3_: borane; 9-BBN: 9-borabicyclo[3.3.1]nonane; nfSTP: non-functionalised sesquiterpenoid; fSTP: functionalised sesquiterpenoid; H_2_O_2_: hydrogen peroxide; GC–MS/FID: gas chromatography with flame-ionisation detection and mass spectrometry; GCxGC–TOF/MS: 2D gas chromatography with time-of-flight mass spectrometry; EtOH: ethanol; 1: step one; anti-Markovnikov hydroboration of the STP, BH_3_ adds boron to the alkene in the less substituted carbon. 2: step two, a three-part oxidation process: initiation by H_2_O_2_ ion donation to the trialkyl borane, rearrangement leading to a hydroxide ion removal, and oxidation completion by reaction with aqueous NaOH, forming alcohol and sodium borate. Supplementary Tables 9 and 10 provide detailed GCxGC–TOF/MS data for panels (**c**) and (**d**), Supplementary File 4 contains supporting raw data for (**b**). Supplementary file 7 contains supporting raw data for GCxGC–TOF/MS.

The hydroboration-oxidation technique is commonly used in organic synthesis for alkenes conversion to alcohols. Compared to halogenation it results in consistent stereochemistry and minimal waste generation. The non-functionalised cyclic primary STP structures produced by our algae were then subjected to hydroboration reactions to cause bulk hydroxy-functionalisation of double-bond locations within the STP backbone (Fig. 4a). Reactions used either borane (BH_3_) or 9-borabicyclo[3.3.1]nonane (9-BBN) reagents and subsequent treatment with hydrogen peroxide (H_2_O_2_) furnished the hydroxy-functionalised STP backbones, thereby mimicking natural chemical-oxygenation processes (Fig. 4a). The nine STPs observed in chromatograms were successfully hydroxylated, including α-santalol/β-santol, cedrenol, 12-nor-preziza-7(15)-en-2-one, khusimol, aristolone, dihydrokaranone, nootkatol, nootkatone, guaiol, globulol, and bisabolol oxide (Fig. 4b and Supplementary Tables 10,11). Using GCxGC–TOF/MS quantification, STP diversity increased in the 9-BBN reaction: from an initial 42 to 103 chemical species (Fig. 4c,d; Supplementary Tables 10,11). Some hydroxylated derivatives matched compounds typically found in agarwood terpene profiles, including α-santalol/β-santol, cedrenol, guaiol, and bisabolol oxide B (Fig. 5 and Supplementary Tables 2,3,10–12).

**Figure 5.**
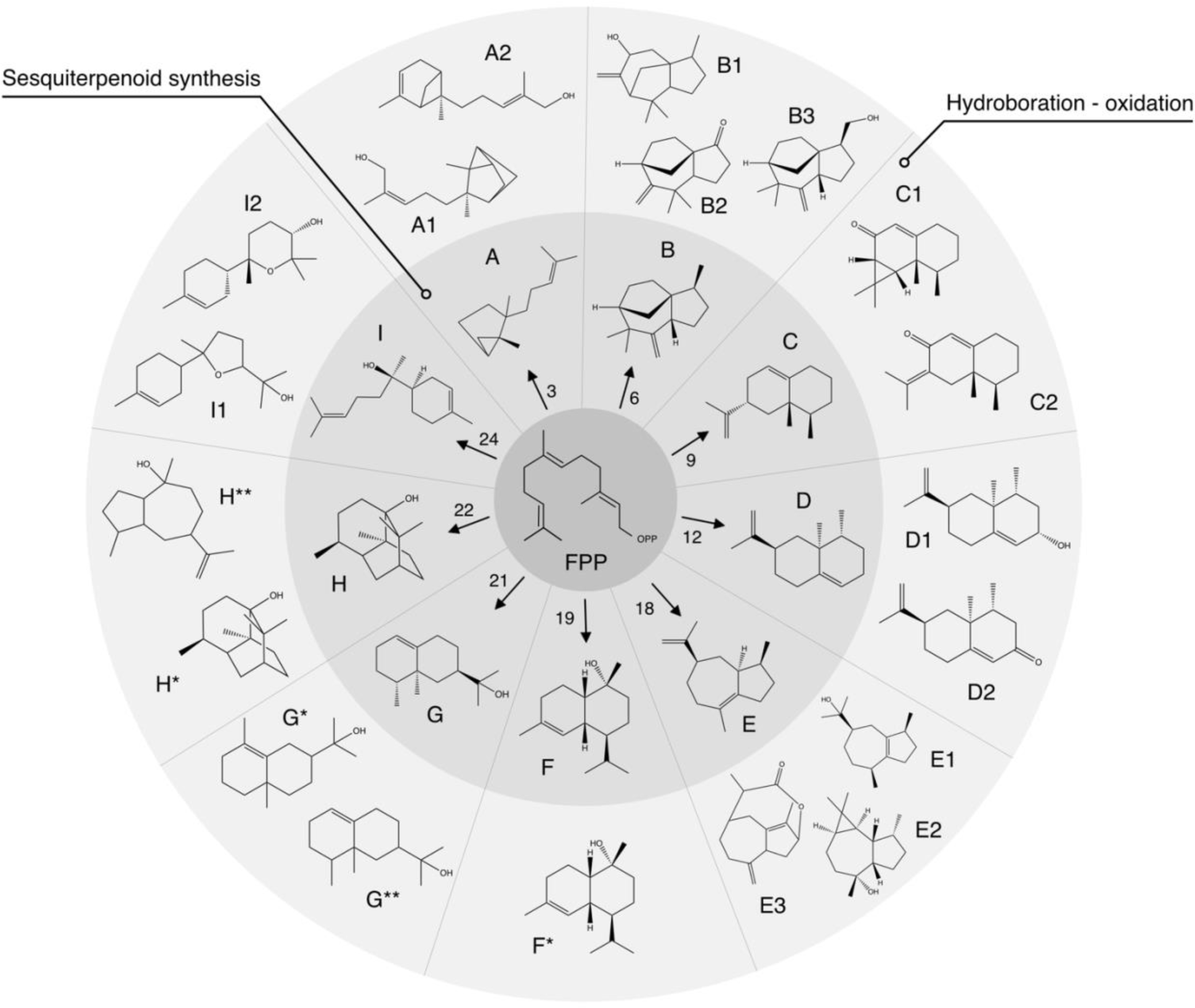
Summary of agarwood-like sesquiterpenoid engineering from combined algal-chemical process. The illustration highlights the array of agarwood-STPs produced and observed with GC–MS. Farnesyl pyrophosphate (FPP) is the precursor of STPs. Numbers indicate STP synthase isoform used to generate each STP from alga. STPs are labelled A–I, and their observed functionalised derivatives achieved through a hydroxy-boron oxidation reaction are labelled with numbers or asterisks. A: Santalene, A1: Santalol, A2: Bergamotene, B: Zizaene, B1: Cedrenol, B3: 12-nor-Preziza-7(15)-en-2-one, B3: Khusimol, C: Aristolochene, C1: Aristolone, C2: Dihydrokaranone, D: Valencene, D1: Nootkatol, D2: Nootkatone, E: δ-Guaiene, E1: Guaiol, E2: Globulol, E3: Guaia-1(10),11-dien-15,2-olide, F: Cadinol, G: Valerianol, G*: Eudesmol, G**: Jinkoheremol, H: Patchoulol, H*: Pogostol, and I: Bisabolol, I1: Bisabolol oxide B, I2: Bisabolol oxide A. Supporting information, including mass spectra and raw data from GC–MS/FID, are in Supplementary Table 11 and Supplementary File 5.

## Discussion

Agarwoods have captivated human interest for centuries due to their complex and diverse fragrances^27^. However, the increasing demand for their resinous wood has placed an unprecedented pressure on their natural sources, particularly older trees native to Southeast Asian forests, and tissue wounding to promote resin formation further increases this pressure^3,8^. In this sample set, we observed that each agarwood and distillate sample contained hundreds of unique STPs and share a subset of these chemicals in varying abundances (Supplementary Files 1–3; Supplementary Tables 2–4). There was no obvious correlation between agarwood prices, origin locations, or sample type, indicating that their market value is subjective, largely based on perceived fragrance or rarity (Supplementary Table 1).

We sought to develop a sustainable approach using metabolically engineered algae to heterologously biosynthesise the STP backbones found in agarwood, followed by oxy-functionalisation using green chemistry as an alternative to tree sources of STPs. Our goal was to generate not only a microbial production alternative to agarwood STPs, but also to develop a sustainable extraction process that generates minimal waste and uses only gentle reagents within the principles of green chemistry and bio-process design^28,29^. We opted for a microalgal host as our bioengineering platform due to its ability to thrive on the nutrients found in wastewaters and CO_2_ as a sole carbon source^30^. This not only enhances the sustainability of the process but also potentially facilitates the conversion of wasted materials into valuable resources, aligning with the principles of circular economy. Our downstream bioprocess uses only low-energy methods for STP extraction and concentration; that is, microbial-milking of STPs during cultivation, liquid–liquid STP transfer to ethanol, and nanofiltration for STP concentration. Using boron and H_2_O_2_, we generated complex, functionalised STP mixtures that resemble those found in agarwood samples in nature.

Endogenous isoprenoids in *C. reinhardtii* are supplied exclusively by the methyl-D-erythritol phosphate (MEP) pathway localised to the plastid (Fig. 2a)^31^. That algae rely on this pathway for essential photosynthetic carotenoids implies an inherent capacity to channel carbon fixed by photosynthesis to isoprenoids, which could be tapped for heterologous terpene biosynthesis. The first reports of micrograms of heterologous terpenes per litre of algal culture^32^ have now been superseded by hundreds of milligrams per litre^25,33^, owing to improved transgene design^34,35^, improved metabolic channelling and high-cell-density cultivation strategies^26,36^. Here, we demonstrate for the first time a combinatorial modification of two metabolic pathways simultaneously in *C. reinhardtii* by perturbing plastid carotenoid and cytoplasmic sterol biosynthesis in the same strain (Fig. 2). This combination improved target STP production by up to 25-fold over wild-type levels, and enabled us to use this strain to produce other STPs of interest. Improvements in transgene optimisation and DNA synthesis capacity together with improved robotic handling and fluorescent screening methods of transformants has facilitated STP isoform benchmarking directly *in alga* (Fig. 3). This is a key step as isoform-performance testing is a role previously conducted only in fermentative microbes. Screening directly *in alga* enabled nuances of expression in this host to be ruled out directly when choosing a target isoform.

Terpene extraction from engineered microbes is a challenge in bioprocess design. Microbes lack the specialised tissues of multicellular organisms to accumulate hydrophobic terpene products, necessitating a physical hydrophobic contactor be used as sink for terpene extraction during cultivation. This can be achieved in numerous ways: using resinous particles^37^, membrane contactors^38^, and through biocompatible solvent–culture two-phase milking^17,39^. We previously found that perfluorinated heat-transfer fluids (namely fluorocarbons) can be used as heavy solvents that sit underneath microbial cultures as an underlay for terpene milking, and that terpenes accumulated in the fluorocarbons rapidly partitioned into ethanol in subsequent liquid–liquid extractions^39^. We leveraged this property to first milk STP skeletons from algal cultures and subsequently pool the ethanol extracts to make a starting mixture of STPs (Fig. 4). This extraction method is advantageous for its simplicity, operating at standard atmospheric conditions, and not requiring complex machinery to yield STPs in ethanol. Although not yet demonstrated at larger scales, the process presented here could readily be incorporated into tubular photobioreactor or even raceway pond algal production facilties.

To create alternatives to agarwood-derived STPs, synthases for the central seven classes of common STP skeletons were required. However, only a handful of δ-guaiene synthase isoforms from *Aquilaria* trees have been reported^11,15,40^. We overcame this limitation by using substitute terpene synthases from different plants and fungi that could reproduce the chemical skeletons we detected in agarwood (Table 1). However, the fragrances of agarwood largely comprise oxy-functionalised derivative STPs. Terpene functionalisation in metabolic engineering is traditionally conducted by the concerted action of highly regio- and stereo-specific enzymes, such as P450s^41^. The chemical complexity of agarwood terpene profiles is both the product of time and likely non-enzymatic chemistry, however, some P450s may be implicated in certain classes of functionalisation observed within^42^. These P450s have yet to be biochemically characterised and would require expression optimisation gene-by-gene to participate in single-STP functionalisation in a microbe. By starting with a mixture of STP skeletons and subjecting them to a chemical process that catalyses oxygenation reactions at sterically accessible double bonds, we could bypass biological functionalisation to obtain complex STP mixtures (Fig. 4). In practice, this procedure mimics parts of the natural maturation process of STPs subjected to abiotic factors in resins over time and successfully reproduced numerous functionalised STPs typical of agarwood samples (Figs. 4, 5).

**Table 1.** Sesquiterpenoid synthase isoforms.

| Isoform number | Origin | Sesquiterpenoid | UniProt | Ref. |
| --- | --- | --- | --- | --- |
| 1 | *Santalum album* | Santalene | E3W202 | 54 |
| 2 | *Clausena lansium* | Santalene | E5LLI1 | 54 |
| 3 | *Cinnamomum camphora* | Santalene | A0A7S6HRZ4 | 55 |
| 4 | *Chrysopogon zizanioides* | Zizaene | AJQ30127 | 37 |
| 5 | *Penicillium expansus* | Aristolochene | C0LW28 | 56 |
| 6 | *Penicillium camemberti* | Aristolochene | A0A0G4NXL3 | 57 |
| 7 | *Penicillium chrysogenum* | Aristolochene | A0A167QKS4 | 58 |
| 8 | *Penicillium roqueforti* | Aristolochene | Q03471 | 59 |
| 9 | *Citrus sinensis* | Valencene | D0UZK2 | 60 |
| 10 | *Callitropsis nootkatensis* | Valencene | S4SC87 | 61 |
| 11 | *Aquilaria crassna* | δ-guaiene | D0VMR6 | 15 |
| 12 | *Aquilaria crassna* | δ-guaiene | D0VMR8 | 15 |
| 13 | *Aquilaria crassna* | δ-guaiene | D0VMR7 | 15 |
| 14 | *Aquilaria sinensis* | δ-guaiene | K9MQ67 | 62 |
| 15 | *Aquilaria sinensis* | δ-guaiene | K9MNV6 | 62 |
| 16 | *Aquilaria sinensis* | δ-guaiene | K9MPP8 | 62 |
| 17 | *Lavandula angustifolia* | τ-Cadinol | U3LW50 | 63 |
| 18 | *Camellia sinensis* | Valerianol | A0A167V661 | 18 |
| 19 | *Camellia hiemalis* | Valerianol | A0A348AUV5 | 18 |
| 20 | *Pogostemon cablin* | Patchoulol | Q49SP3* | 64 |
| 21 | *Cynara cardunculus* | Bisabolol | XP_024994640 | 65 |

\*Mutation C415F and H454A

Traditional methods of oxidation or hydroxylation of STPs have often involved the use of heavy-metal catalysts^43^ (such as chromium and mercury), strong oxidising agents^44^ (i.e., osmium tetroxide), toxic solvents^45^ (i.e., benzene), or hazardous reagents^46^ (i.e., chlorine) — all of which pose serious environmental and health risks. We instead used organoboron reagents to introduce hydroxyl groups to the most accessible double bonds of the STP skeletons, and eliminated use of heavy-metal reagents and hazardous by-products associated with conventional methods (Fig. 4b–d, Fig. 5 and Supplementary Table 11).

Because mono- and di-terpenes are also present in agarwood (Fig. 1, Supplemental File 1), using more complex mixtures of STPs obtained by additional metabolic engineering will undoubtedly increase the complexity achievable in synthetically functionalised mixtures. Here, we did not express specific α-humulene, β-eudesmol^14^ and vetispiradiene synthases, which could broaden the complexity of the starting mixtures. The combination of possibilities for these mixtures is immense, offering wide scope for additional combinations of microbial engineered terpene production and synthetic chemistry to yield fragrant mixtures. Having said that, this work also should not be considered to over-promise the potential of synthetic-biology alternatives to natural agarwood. Previous hype around novel terpene production techniques has created instability in the natural-product market, leading to economic challenges within legitimate supply chains^47^. This process should instead initially be considered complementary to traditional agarwood production, especially silviculture or tissue-culture alternatives, as one pathway to replicate some of the aromatic functionalised STPs.

In summary, we have developed a strategy following green-bioprocess design principles that combines metabolic engineering, terpene milking, and pharmaceutical-grade chemical processing to obtain terpene mixtures that mimic some of the natural chemical diversity found in agarwood. While our set of 103 semi-synthetic engineered sesquiterpenoids does not fully mirror the diversity and yields of agarwood, our approach presents a sustainable complement to legitimate sources of these products, and offers a potential solution to the current demand for agarwood that has surpassed natural supply for these culturally important goods. We demonstrate the feasibility of a cyclic, synthetic-biology-mediated and chemically complemented process to generate functionalised STP mixtures.

## Materials and methods

### Agarwood sample collection and processing

Agarwood samples were procured from the old-town market "Al Balad" in Jeddah, Saudi Arabia (21.481° N, 39.187° E) in Winter 2023. The collection includes 36 different samples of agarwood chips, dried *Aquilaria* spp. wood (also known as "bahkour"), and 22 agarwood steam-distillate oils, known as "oudh". The origins of these samples cannot be accurately traced; however, many were labelled with their country of origin. The samples varied in price at the time of purchase, reflecting their purported rarity, the density of fragrant compounds, and complexity of aromatic notes (Supplementary Table 1). It was determined that the appropriate organic solvent for extraction was acetone based on preliminary analysis with different solvents, and each agarwood sample was then diluted in order to obtain chromatograms with clear peaks for product detection and identification. The wood samples were weighed (1 g) and ground into a fine powder using a combination of freeze–thaw cycles with liquid nitrogen and mechanical grinding. The homogenised samples were immersed in 5 mL of 1:1 hexane:acetone and subjected to ultrasonic agitation at 40°C for 8 h to facilitate terpenoid extraction. The samples were then passed through a 0.2 µm filter to obtain clear solvent extracts that were evaporated under a nitrogen stream for 20 min to concentrate the terpenoids in the samples. Concentrated samples were resuspended in 500 µL acetone and stored at –20°C until gas chromatography–flame-ionisation detector and mass spectrometry (GC–MS/FID) and two-dimensional gas chromatography with time-of-flight mass spectrometry (GCxGC–TOF/MS) analyses. All photographs were captured with a Canon EOS RP camera using a Canon RF 24–105 mm f/4-7.1 IS STM lens (Canon, Tokyo, Japan) and ColorChecker Passport (CCPP2, Calibrite LLC, DE, USA) used for colour calibration.

### Algal strain cultivation

*Chlamydomonas reinhardtii* strain UPN22^26^ was used for all experiments. *C. reinhardtii* UPN22 is a derivative of the UVM4 strain^48^. It has been genetically enhanced to use phosphite and nitrate as a sole source of phosphorous and nitrogen, respectively, to minimise contamination and maximise cell densities in cultivation^26^. Strains were cultured in Tris-acetate phosphite nitrate (TAPhi-NO_3_, ^26^) liquid medium with shaking at 120 rpm or on solid agar and 150 µmol m^−2^ s^−1^ light intensity from a combination of cool- and warm white LED tubes (light spectra as reported in^26^). 500 mL algal cultures were agitated with stirring in Erlenmeyer flasks under the same growth conditions.

### Plasmid construction, algal transformation, and screening for sesquiterpenoid synthase expression

Heterologous expression of sesquiterpenoid synthases (STPSs) in *C. reinhardtii* was achieved through synthetic transgene redesign based on the amino-acid sequences of each STPS, following previously established protocols^34,49^. We selected 21 STPSs from various species encoding isoforms that yield nine different STP skeletons (Table 1). Targeted STPSs for aristolochene, δ-guaiene, santalene, valencene, valerianol, zizaene, τ-cadinol, bisabolol, and patchoulol were designed; all accession numbers are listed in Table 1. Their amino-acid sequences were used to generate algal-adapted nucleotide coding sequences using the Intronserter programme^34^. This programme back-translates amino-acid sequences to frequently used codons, removes unwanted restriction sites, and systematically integrates the first intron of *Cr*RBCS2i1 at a set distance to enable expression from the *C. reinhardtii* nuclear genome, as previously reported^49,50^. Algal-adapted coding sequences were synthesised and sub-cloned into pOptimized_3 expression plasmids^35^ by Genscript (Piscataway, NJ, USA). Ketocarotenoid biosynthesis in alga was achieved by transformation with the pOpt2_*Cr*BKT_aadA plasmid^51^, and knockdown of *C. reinhardtii* squalene synthase (Uniprot: A8IE29) was achieved using the previously reported luciferase–artificial microRNA expression plasmid pOpt2_ *c*CA_*g*Luc-TAA_i3_ami_Spec^24^. Both plasmids confer resistance to spectinomycin as a selectable marker. Combining both cassettes into a single plasmid was unsuccessful. Co-transformation of both plasmids and selection on spectinomycin was combined with robotics assisted colony picking and plate-level screening of 768 transformants to find those with luciferase activity (indicative of SQS knockdown) and a brown-colony phenotype from *Cr*BKT-mediated ketocarotenoid biosynthesis. All plasmid constructs used are listed in Supplementary Table 6 and the complete annotated plasmid sequences are available in Supplementary File 5.

The glass-bead protocol was used to transform the nuclear genome of *C. reinhardtii* with a plasmid DNA^52^. Each plasmid was linearised using restriction enzymes (*Xba*I + *Kpn*I, Thermo Scientific FastDigest), and 10 µg of DNA was used for each transformation. Following an ∼8 h recovery period in liquid TAPhi-NO_3_ medium under low light, algal cells were plated on a selective medium with paromomycin (10 µg mL^−1^), spectinomycin (200 µg mL^−1^) or zeocin (15 µg mL^—1^) antibiotics either individual or combinations of selection agents relative to each target plasmid. Plates were illuminated continuously for ∼7 d before colony picking. A PIXL robot (Singer Instruments, Watchet, UK) transferred up to 384 colonies per transformation event to TAPhi-NO_3_ agar plates. After an additional 3 d, a ROTOR robot (Singer Instruments) was used to replicate colonies onto new medium and plates containing amido black (150 µg mL^−1^) for fluorescence screening as previously described^35^.

All algal-optimised STPSs were expressed as fusions with mVenus (yellow fluorescent protein) or the monomeric teal (cyan) fluorescent protein 1 (mTFP1). Transgene expression of each STPS was determined by fluorescence imaging at the agar-plate level, as previously described^32,35^. Chlorophyll fluorescence was observed with 2 sec of 475/20 nm excitation and 640/160 nm emission to show colony presence/absence on amido black-containing plates. Cyan-green fluorescence was captured with 420/20 nm excitation and 480/20 nm emission filters using 2.5 min exposure. Yellow fluorescent signals were captured with 504/10 nm excitation and 530/10 nm emission filters with 30-sec exposures. Transformants displaying strong fluorescent-protein signals were selected and inoculated into 12-well plates containing 2 mL of liquid TAPhi-NO_3_ medium and grown with shaking at 160 rpm. Predicted molecular masses of the expressed heterologous STPS–FP fusions were verified using sodium dodecyl sulphate-polyacrylamide gel electrophoresis (SDS–PAGE) in-gel fluorescence against the fusion-protein fluorescent reporter.

### Heterologous sesquiterpenoid biosynthesis analyses

For each transformant, biosynthesis of heterologous STPs was assessed by gas chromatography–mass spectrometry (GC–MS/FID). Four individual transformants containing each plasmid with the highest fluorescence signals were selected, and solvent–culture two-phase living extractions were performed using a 10% v/v dodecane–culture overlay in 6-well plates, as previously reported^17,53^. Cultivations were performed in biological triplicate for 6 d in 4.5 mL TAPhi-NO_3_ media with 500 µL dodecane overlay. Phases were separated, and culture samples were taken for cell-density analysis by flow cytometry as previously described^26^, and dodecane was spun at 20,000 x *g* for 3 min, then 150 µL clarified solvent was transferred into amber GC vials in triplicate prior to analysis.

Samples were analysed as previously described^26,38^ using an Agilent 7890A gas chromatograph equipped with a mass spectrometer and a flame-ionisation detector (GC– MS/FID). The gas chromatograph comprises a 5975C inert MSD with a triple-axis detector and a DB-5MS column (30 m × 0.25 mm i.d., 0.25 μm film thickness). The injector, interface, and ion-source temperature profiles were set to 250°C, 250°C, and 220°C, respectively. In splitless mode, 1 μL of the sample was injected using an autosampler (G4513A, Agilent). The column flow was constant at 1 mL min^−1^, with helium as carrier gas. The initial GC oven temperature was set to 80°C for 1 min, increased to 120°C at 10°C min^−1^, raised to 160°C at 3°C min^−1^, and to 240°C at 10°C min^−1^, holding for 3 min. After a 12-min solvent delay, mass spectra were recorded using a scanning range of 50–750 *m/z* at 20 scans per second. Chromatograms were analysed with MassHunter Workstation software version B.08.00 (Agilent), and STPs were identified using the National Institute of Standards and Technology (NIST) library (Gaithersburg, MD, USA). Further identification was conducted with purified standard calibration curves ranging from concentrations of 1–1,200 μM in dodecane of δ-guaiene (CAT#B942760), patchoulol (CAT#P206200), santalene (CAT#S15065), valerianol (CAT#V914000, Toronto Research Chemicals, ON, Canada), bisabolol (CAT#95426), valencene (CAT#06808), cedrene (CAT#22133, Sigma-Aldrich, MO, USA) (Supplementary Fig. 4). For compound identification, retention-time acquisition, internal digital-library calibration, and method development, we used a set of 12 microampules containing a standard terpene mixture, which covered 98 terpenes at 1 mM in methanol (CAT# MSITPN101, MetaSci, ON, Canada, Supplementary Table 4).

### Two-dimensional Gas chromatography time-of-flight mass spectrometry (GCxGC– TOF/MS) analysis

Comprehensive two-dimensional gas chromatography time-of-flight mass spectrometry (GCxGC–TOF/MS) analysis of agarwood and distillate extracts in acetone was performed using an Agilent 7890B gas-chromatography system equipped with a Zoex ZX1 cryogenic thermal modulator and a JEOL TOF MS (AccuTOF GCx-plus, JEOL, Japan). The GCxGC system featured a normal (non-polar x mid-polar) two-dimensional column configuration, comprising a first-dimension column with a 30 m non-polar HP-5MS UI capillary column (5%-phenyl-methylpolysiloxane) and a second-dimension column with a 2 m mid-polar BPX-50 capillary column (50% phenyl polysilphenylene-siloxane). We used helium (99.999%) as the carrier gas at a constant flow rate of 0.8 mL min^−1^. The GCxGC–TOF/MS injector temperature was maintained at 300°C with a 10:1 split ratio. The oven temperature was initially held at 80°C for 1 min, then increased to 325°C at 2°C min^−1^. The modulation period was set at 6 s with a pulse time of 0.35 ms. The mass spectrometer operated in electron ionisation (EI+) mode at 70 eV. Both the transfer-line and ion-source temperatures were maintained at 250°C. The detector voltage of TOF was set to 2,500 Volts, and data were acquired at a rate of 50 Hz. Mass spectra were obtained within a mass-to-charge ratio (*m/z)* from 50 to 600.

### Gas chromatography data analysis

The analysis of terpenoid extracts followed a procedure similar to previously reported methods^26,38,39^, and qualitative analyses primarily relied on the retention index and match factor. The GC–MS/FID data were processed using the MassHunter Workstation software version B.08.00 (Agilent Technologies, USA). The identification of compounds was assisted by the NIST Mass Spectral Library Version 2.3 (National Institute of Standards and Technology, Gaithersburg, MD, USA). The mass-spectral data, derived from the GCxGC– TOF/MS analysis, were evaluated with GC Image^TM^ Version 2.9 software (Lincoln, NE, USA) and referenced against the NIST2020/EPA/NIH EI Mass Spectral Library. The spectral data were cross-referenced with library spectra to identify potential chemical structures, facilitated by calculating the match factor and probability, thereby generating a list of probable compound matches. The match factor directly compares the unknown mass spectrum peaks with those in the library spectra, indicating their similarity. In contrast, the probability determines the relative likelihood of the list of hits being accurate, assuming that the unknown spectrum is present in the library. Using these metrics provides the relative assuredness of matching chemical structures within the library spectra.

### Sesquiterpenoid production and concentration

Transformants of best-performing STPS isoforms that accumulated the most of each heterologous sesquiterpenoid in screening conditions, were subjected to solvent–culture two-phase cultivations at 300 mL scale in TAPhi-NO3 medium using FC-3283 as a solvent underlay phase, as previously described^39^. FC-3283 is a perfluorinated amine that is inert and denser than water, forming an underlay to the culture and accumulating heterologous terpene products from the algae. After 6 days of cultivation, gravity and gentle centrifugation separated the fluorocarbon and cultures. After centrifugation to clarify, fluorocarbons were subjected to liquid–liquid extraction to partition accumulated STPs into an equal volume of 96% ethanol. The mixture was shaken for 16 h at room temperature at 200 rpm. Next, samples were again centrifuged gently to further separate the phases. FC-3283, after ethanol extraction, can be reused on algal cultures and is effectively recycled in this process. 500 μL aliquots from each phase were sampled and stored in separate GC vials at –20°C for subsequent analysis and separation performance quantification.

For each of the nine STPs biosynthesised by the algae, 20 mL 96% ethanol-containing STOs were generated. The ethanol fractions were pooled and subjected to organic solvent nanofiltration (OSN) in a dead-end cell to concentrate the terpenes in ethanol without evaporative losses. A Duramem solvent-resistant membrane (Evonik, Germany) with a nominal molecular weight cut-off value of 300 g mol^−1^ suitable for STP retention and chemical compatibility with ethanol was selected. The 200 mL ethanol–sesquiterpenoid mixture was loaded into the OSN chamber containing a 16 mm membrane disc and subjected to 20 bar pressure delivered by CO_2_ as an inert gas, to drive the ethanol phase through the nanofiltration membrane at 2.84 L m^−2^ h^−1^ flux. While ethanol permeated the membrane, STPs were kept in the retentate due to their molecular weight and consequently concentrated in the ethanol. The permeate ethanol is suitable for recycling and can be reused in subsequent liquid–liquid extraction processes.

### Bulk hydroxylation of sesquiterpenoid backbones

To selectively introduce hydroxyl groups at the double bonds present in the terpenes, hydroboration-oxidation reactions were performed using two distinct organoboron reagents: borane–tetrahydrofuran complex (BH_3_·THF) (CAT#176192, Sigma-Aldrich) and 9-borabicyclo[3.3.1]nonane (151076, Sigma-Aldrich). Three different stoichiometric ratios (0.5, 1.0, and 2.0) were explored for each reagent to investigate different degrees of hydroxylation within the STP mixture. The sterically hindered 9-BBN was anticipated to selectively react with the least sterically demanding sites of the terpenes, while the borane–tetrahydrofuran complex was expected to facilitate hydroxylation even at the more challenging endocyclic positions. To remove residual ethanol and water content from the concentrated terpene mixture, a rotary evaporator was used under reduced pressure at 40°C, ensuring effective removal without compromising the integrity of the terpenes. To ensure the complete removal of water, molecular sieves (CAT#105734, Sigma-Aldrich) were added to the terpene–THF solvent mixture. The reactions were carried out in anhydrous tetrahydrofuran (THF, Sigma-Aldrich) to enable removal by evaporation after completion. Hydrogen peroxide (H_2_O_2_, 30%, VWR Chemicals) and sodium hydroxide aqueous solution (NaOH, Sigma-Aldrich) were used to convert the organoboron intermediates into non-toxic, water-soluble boric acid after the reaction. All subsequent terpene derivatives were extracted from the aqueous mixture with ethyl acetate.

### Data analysis

To evaluate the impact of engineered modifications on STP biosynthesis and growth characteristics in *C. reinhardtii* strains, one-way analysis of variance (ANOVA) was performed to compare mean STP (patchoulol) titres and growth rates among different transformants and the parental control strain. This statistical approach allowed for the simultaneous analysis of differences between multiple groups, to provide a robust assessment of the experimental manipulations. ANOVAs were performed separately for STP titres and growth rates under mixotrophic and phototrophic conditions. Post-hoc pairwise comparisons using Tukey’s HSD were conducted to identify which groups were statistically significantly different. A one-way ANOVA was used to assess the effects of differences in STP biosynthesis among tested strategies (single or double transformation), followed by a post hoc Tukey’s HSD test for specific pairwise data-set comparisons. Mean production values (in fg cell^−1^ and mg L^−1^) for each STP were compared, considering standard deviations to evaluate data variability. Mean values were considered statistically significantly different at a level of *p* < 0.05. For data analyses, JMP v.16 (SAS Institute Inc, NC, USA) and Rstudio 3.6.2 (Posit Software, Boston, USA) were used. Data visualisation was done using JMP v.16 and GraphPad Prism v.10 (GraphPad Software, MA, USA). Image adjustments involved ColourChecker calibration (Calibrite LLC, DE, USA) paired with Adobe Lightroom (Adobe Inc., CA, USA) for colour accuracy. Images were cropped and organised in Affinity Photo v.1.10.6 (Serif Ltd., WB, UK). Diagrams and illustrations were made using Affinity Designer v.1.10.6 (Serif Ltd., WB, UK), chemical structures with ChemDraw v.20.1 (PerkinElmer, MA, USA) and all visual elements were harmonised in Affinity Publisher v.1.10.6 (Serif Ltd., WB, UK).

## Additional information

Data supporting the findings of this work are available within the paper and its Supplementary material. Source GCxGC–TOF/MS, GC–MS/FID for the findings of this study and genetic data can be accessed in DRYAD (https://doi.org/10.5061/dryad.h70rxwdq5).

## Acknowledgements

This work received support from the competitive research grant (4715 to K.J.L.) and baseline research funding from King Abdullah University of Science and Technology (KAUST) to G.S. and K.J.L. K.J.L would like to express his gratitude to Ahmed Alfahad of King Abdulaziz City for Science and Technology (KACST) who found and suggested several of the terpene synthases used in this work and whose discussions inspired the conception of this work.

## Author contributions

S.G., S.O., G.B.W., and K.J.L. conceptualised and planned the project. S.G. and K.J.L. designed the genes and plasmids for this study. S.G. and S.O. undertook in situ extraction experiments and analysed the GC–FID/MS results. S.G. and V.G.S. conducted experiments and interpreted the GCxGC–TOF/MS data. C.O. executed the nanofiltration experiments and M.G. conducted the hydroxy-boron oxidation experiments in the lab of G.S., who also participated in experimental design. S.G., S.O., G.B.W., and K.J.L. analysed the data and wrote the manuscript. K.J.L. supervised, managed and was responsible for funding acquisition. All authors reviewed and discussed the manuscript.

## References

1. Yin, Y., Jiao, L., Dong, M., Jiang, X. & Zhang, S. Wood Resources, Identification, and Utilization of Agarwood in China. in Agarwood: Science Behind the Fragrance (ed. Mohamed, R.) 21–38 (Springer, 2016). doi:10.1007/978-981-10-0833-7_2.

2. Ahmaed, D. T. Investigation of Agarwood Compounds in Aquilaria malaccensis & Aquilaria Rostrata Chipwood by Using Solid Phase Microextraction. Biomed. J. Sci. Tech. Res. 1, (2017).

3. Gao, Z.-H. & Wei, J.-H. Molecular Mechanism Studies of Terpenoid Biosynthesis in Agarwood. in Agarwood: Science Behind the Fragrance (ed. Mohamed, R.) 73–87 (Springer, 2016). doi:10.1007/978-981-10-0833-7_5.

4. Naziz, P. S., Das, R. & Sen, S. The scent of stress: Evidence from the unique fragrance of agarwood. Front. Plant Sci. 10, (2019).

5. López-Sampson, A. & Page, T. History of Use and Trade of Agarwood. Econ. Bot. 72, 107–129 (2018).

6. Hashim, Y. Z. H.-Y., Kerr, P. G., Abbas, P. & Mohd Salleh, H. Aquilaria spp. (agarwood) as source of health beneficial compounds: A review of traditional use, phytochemistry and pharmacology. J. Ethnopharmacol. 189, 331–360 (2016).

7. Global Market Study on Agarwood Essential Oil Market: Synthetic Biological Production of Agarwood Essential Oil Profiting Market Growth. Persistence Market Research https://www.persistencemarketresearch.com/market-research/agarwood-essential-oil-market.asp.

8. UNEP-WCMC, C. Checklist of CITES Species. CITES Secr. Geneva Switz. UNEP-WCMC Camb. U. K. (2014).

9. Barden, A., Awang Anak, N., Mulliken, T. A. & Song, M. Heart of the matter : agarwood use and trade and CITES implementation for Aquilaria malaccensis. (TRAFFIC International, 2000).

10. Gao, M. et al. Overview of sesquiterpenes and chromones of agarwood originating from four main species of the genus Aquilaria. RSC Adv. 9, 4113–4130 (2019).

11. Ran, J. et al. Identification of sesquiterpene synthase genes in the genome of Aquilaria sinensis and characterization of an α-humulene synthase. *J*. For. Res. 34, 1117–1131 (2023).

12. Dada, L. et al. Role of sesquiterpenes in biogenic new particle formation. Sci. Adv. 9, eadi5297 (2023).

13. Tajuddin, S. N., Aizal, C. M. & Yusoff, M. M. Resolution of Complex Sesquiterpene Hydrocarbons in Aquilaria malaccensis Volatile Oils Using Gas Chromatography Technique. in Agarwood: Science Behind the Fragrance (ed. Mohamed, R.) 103–124 (Springer, 2016). doi:10.1007/978-981-10-0833-7_7.

14. Promdonkoy, P., Sornlek, W., Preechakul, T., Tanapongpipat, S. & Runguphan, W. Metabolic Engineering of Saccharomyces cerevisiae for Production of Fragrant Terpenoids from Agarwood and Sandalwood. Fermentation 8, 429 (2022).

15. Kumeta, Y. & Ito, M. Characterization of δ-guaiene synthases from cultured cells of Aquilaria, responsible for the formation of the sesquiterpenes in agarwood. Plant Physiol. 154, 1998–2007 (2010).

16. Kurosaki, F., Kato, T., Misawa, N. & Taura, F. Efficient Production of δ-Guaiene, an Aroma Sesquiterpene Compound Accumulated in Agarwood, by Mevalonate Pathway-Engineered *Escherichia coli* Cells. Adv. Biosci. Biotechnol. 07, 435–445 (2016).

17. Beekwilder, J. et al. Valencene synthase from the heartwood of Nootka cypress (Callitropsis nootkatensis) for biotechnological production of valencene. Plant Biotechnol. J. 12, 174–182 (2014).

18. Hattan, J. ichiro et al. Identification of novel sesquiterpene synthase genes that mediate the biosynthesis of valerianol, which was an unknown ingredient of tea. Sci. Rep. 8, (2018).

19. Sun, Y., Wu, S., Fu, X., Lai, C. & Guo, D. De novo biosynthesis of τ-cadinol in engineered Escherichia coli. Bioresour. Bioprocess. 9, 29 (2022).

20. Felicetti, B. & Cane, D. E. Aristolochene synthase: Mechanistic analysis of active site residues by site-directed mutagenesis. J. Am. Chem. Soc. 126, 7212–7221 (2004).

21. Aguilar, F., Scheper, T. & Beutel, S. Modulating the precursor and terpene synthase supply for the whole-cell biocatalytic production of the sesquiterpene (+)-zizaene in a pathway engineered *E. coli*. Genes 10, 1–15 (2019).

22. Lauersen, K. J. Eukaryotic microalgae as hosts for light-driven heterologous isoprenoid production. Planta 249, 155–180 (2019).

23. Perozeni, F. & Baier, T. Current Nuclear Engineering Strategies in the Green Microalga Chlamydomonas reinhardtii. life 13, (2023).

24. Wichmann, J., Baier, T., Wentnagel, E., Lauersen, K. J. & Kruse, O. Tailored carbon partitioning for phototrophic production of (E)-α-bisabolene from the green microalga Chlamydomonas reinhardtii. Metab. Eng. 45, 211–222 (2018).

25. Yahya, R. Z., Wellman, G. B., Overmans, S. & Lauersen, K. J. Engineered production of isoprene from the model green microalga Chlamydomonas reinhardtii. Metab. Eng. Commun. 16, e00221 (2023).

26. Abdallah, M. N., Wellman, G. B., Overmans, S. & Lauersen, K. J. Combinatorial Engineering Enables Photoautotrophic Growth in High Cell Density Phosphite-Buffered Media to Support Engineered Chlamydomonas reinhardtii Bio-Production Concepts. Front. Microbiol. 13, 885840 (2022).

27. López-Sampson, A. & Page, T. History of Use and Trade of Agarwood. Econ. Bot. 72, 107–129 (2018).

28. Anastas, P. & Eghbali, N. Green Chemistry: Principles and Practice. Chem. Soc. Rev. 39, 301–312 (2009).

29. Koutinas, M., Kiparissides, A., Pistikopoulos, E. N. & Mantalaris, A. BIOPROCESS SYSTEMS ENGINEERING: TRANSFERRING TRADITIONAL PROCESS ENGINEERING PRINCIPLES TO INDUSTRIAL BIOTECHNOLOGY. Comput. Struct. Biotechnol. J. 3, e201210022 (2012).

30. de Freitas, B. B., Overmans, S., Medina, J. S., Hong, P.-Y. & Lauersen, K. J. Biomass generation and heterologous isoprenoid milking from engineered microalgae grown in anaerobic membrane bioreactor effluent. Water Res. 229, 119486 (2023).

31. Lohr, M., Schwender, J. & Polle, J. E. W. Isoprenoid biosynthesis in eukaryotic phototrophs: A spotlight on algae. Plant Sci. 185–186, 9–22 (2012).

32. Lauersen, K. J. et al. Efficient phototrophic production of a high-value sesquiterpenoid from the eukaryotic microalga Chlamydomonas reinhardtii. Metab. Eng. 38, 331–343 (2016).

33. Einhaus, A. et al. Engineering a powerful green cell factory for robust photoautotrophic diterpenoid production. Metab. Eng. 73, 82–90 (2022).

34. Jaeger, D., Baier, T. & Lauersen, K. J. Intronserter, an advanced online tool for design of intron containing transgenes. Algal Res. 42, 101588 (2019).

35. Gutiérrez, S., Wellman, G. B. & Lauersen, K. J. Teaching an old ‘doc’ new tricks for algal biotechnology: Strategic filter use enables multi-scale fluorescent protein signal detection. Front. Bioeng. Biotechnol. 10, 979607 (2022).

36. Freudenberg, R. A., Baier, T., Einhaus, A., Wobbe, L. & Kruse, O. High cell density cultivation enables efficient and sustainable recombinant polyamine production in the microalga Chlamydomonas reinhardtii. Bioresour. Technol. 323, 124542 (2021).

37. Aguilar, F., Scheper, T. & Beutel, S. Improved production and in situ recovery of sesquiterpene (+)-zizaene from metabolically-engineered E. Coli. Molecules 24, (2019).

38. Overmans, S. et al. Continuous extraction and concentration of secreted metabolites from engineered microbes using membrane technology. Green Chem. 10.1039.D2GC00938B (2022) doi:10.1039/D2GC00938B.

39. Overmans, S. & Lauersen, K. J. Biocompatible fluorocarbon liquid underlays for in situ extraction of isoprenoids from microbial cultures. RSC Adv. 12, 16632–16639 (2022).

40. Azzarina, A. B., Mohamed, R., Lee, S. Y. & Nazre, M. Temporal and spatial expression of terpene synthase genes associated with agarwood formation in Aquilaria malaccensis Lam. N. Z. J. For. Sci. 46, (2016).

41. Pateraki, I., Heskes, A. M. & Hamberger, B. Cytochromes P450 for Terpene Functionalisation and Metabolic Engineering. in Advances in biochemical engineering/biotechnology vol. 123 107–139 (2015).

42. Das, A. et al. Genome-wide investigation of Cytochrome P450 superfamily of Aquilaria agallocha: Association with terpenoids and phenylpropanoids biosynthesis. Int. J. Biol. Macromol. 234, 123758 (2023).

43. Balali-Mood, M., Naseri, K., Tahergorabi, Z., Khazdair, M. R. & Sadeghi, M. Toxic Mechanisms of Five Heavy Metals: Mercury, Lead, Chromium, Cadmium, and Arsenic. Front. Pharmacol. 12, (2021).

44. Schroeder, M. Osmium tetraoxide cis hydroxylation of unsaturated substrates. Chem. Rev. 80, 187–213 (1980).

45. Yardley-Jones, A., Anderson, D. & Parke, D. V. The toxicity of benzene and its metabolism and molecular pathology in human risk assessment. Occup. Environ. Med. 48, 437–444 (1991).

46. Winder, C. The Toxicology of Chlorine. Environ. Res. 85, 105–114 (2001).

47. Dalziell, J. & Rogers, W. Are the Ethics of Synthetic Biology Fit for Purpose? A Case Study of Artemisinin [Point of View]. Proc. IEEE 110, 511–517 (2022).

48. Neupert, J., Karcher, D. & Bock, R. Generation of *Chlamydomonas* strains that efficiently express nuclear transgenes. Plant J. 57, 1140–1150 (2009).

49. Baier, T., Wichmann, J., Kruse, O. & Lauersen, K. J. Intron-containing algal transgenes mediate efficient recombinant gene expression in the green microalga *Chlamydomonas reinhardtii*. Nucleic Acids Res. 46, 6909–6919 (2018).

50. Baier, T., Jacobebbinghaus, N., Einhaus, A., Lauersen, K. J. & Kruse, O. Introns mediate post-transcriptional enhancement of nuclear gene expression in the green microalga Chlamydomonas reinhardtii. PLOS Genet. 16, e1008944 (2020).

51. Cazzaniga, S., Perozeni, F., Baier, T. & Ballottari, M. Engineering astaxanthin accumulation reduces photoinhibition and increases biomass productivity under high light in Chlamydomonas reinhardtii. Biotechnol. Biofuels Bioprod. 15, 77 (2022).

52. Kindle, K. L. High-frequency nuclear transformation of *Chlamydomonas reinhardtii*. Proc. Natl. Acad. Sci. U. S. A. 87, 1228–1232 (1990).

53. Lauersen, K. J. et al. Phototrophic production of heterologous diterpenoids and a hydroxy-functionalized derivative from Chlamydomonas reinhardtii. Metab. Eng. 49, 116–127 (2018).

54. Jones, C. G. et al. Sandalwood Fragrance Biosynthesis Involves Sesquiterpene Synthases of Both the Terpene Synthase (TPS)-a and TPS-b Subfamilies, including Santalene Synthases. J. Biol. Chem. 286, 17445–17454 (2011).

55. Di Girolamo, A. et al. The santalene synthase from Cinnamomum camphora: Reconstruction of a sesquiterpene synthase from a monoterpene synthase. Arch. Biochem. Biophys. 695, 108647 (2020).

56. La Guerche, S., Dauphin, B., Pons, M., Blancard, D. & Darriet, P. Characterization of Some Mushroom and Earthy Off-Odors Microbially Induced by the Development of Rot on Grapes. J. Agric. Food Chem. 54, 9193–9200 (2006).

57. Cheeseman, K. et al. Multiple recent horizontal transfers of a large genomic region in cheese making fungi. Nat. Commun. 5, 2876 (2014).

58. Specht, T., Dahlmann, T. A., Zadra, I., Kürnsteiner, H. & Kück, U. Complete Sequencing and Chromosome-Scale Genome Assembly of the Industrial Progenitor Strain P2niaD18 from the Penicillin Producer *Penicillium chrysogenum*. Genome Announc. 2, e00577–14 (2014).

59. Proctor, R. H. & Hohn, T. M. Aristolochene synthase. Isolation, characterization, and bacterial expression of a sesquiterpenoid biosynthetic gene (Ari1) from Penicillium roqueforti. J. Biol. Chem. 268, 4543–4548 (1993).

60. Sharon-Asa, L. et al. Citrus fruit flavor and aroma biosynthesis: isolation, functional characterization, and developmental regulation of Cstps1, a key gene in the production of the sesquiterpene aroma compound valencene. Plant J. 36, 664–674 (2003).

61. Beekwilder, J. et al. Valencene synthase from the heartwood of Nootka cypress (Callitropsis nootkatensis) for biotechnological production of valencene. Plant Biotechnol. J. 12, 174–182 (2014).

62. Xu, Y. et al. Identification of genes related to agarwood formation: transcriptome analysis of healthy and wounded tissues of Aquilaria sinensis. BMC Genomics 14, 227 (2013).

63. Jullien, F. et al. Isolation and functional characterization of a τ-cadinol synthase, a new sesquiterpene synthase from Lavandula angustifolia. Plant Mol. Biol. 84, 227–241 (2014).

64. Zhou, L. et al. Enhancement of Patchoulol Production in *Escherichia coli via* Multiple Engineering Strategies. J. Agric. Food Chem. 69, 7572–7580 (2021).

65. Han, G. H. et al. Fermentative production and direct extraction of (−)-α-bisabolol in metabolically engineered Escherichia coli. Microb. Cell Factories 15, 185 (2016).

