## Supplementary information for "A green alternative to fragrant agarwood sesquiterpenoid production"

### Supplementary Figures

**Supplementary Figure 1.** Optimisation of *Chlamydomonas reinhardtii* metabolism for heterologous sesquiterpene production.

**Supplementary Figure 2.** Growth assessment of engineered *Chlamydomonas reinhardtii* engineered strains.

**Supplementary Figure 3.** Calibration curves of sesquiterpenoid standards.

**Supplementary Figure 4.** Image of ColorChecker Passport.

### Supplementary Tables

**Supplementary Table 1.** Agarwood samples description.

**Supplementary Table 2.** Sesquiterpenoids identified across agarwood (bakhour) samples using GCxGC-TOF/MS.

**Supplementary Table 3.** Sesquiterpenoids identified across agarwood distillates (Oudh) samples using GCxGC-TOF/MS

**Supplementary Table 4.** Terpenoid standard mixture analysed using GCxGC-TOF/MS.

**Supplementary Table 5.** *Chlamydomonas reinhardtii* metabolism optimisation to produce heterologous sesquiterpenes.

**Supplementary Table 6**. List of plasmids.

**Supplementary Table 7.** Sesquiterpenoids quantification (Single transformation).

**Supplementary Table 8.** Sesquiterpenoids quantification (Double transformation).

**Supplementary Table 9.** Summary of sesquiterpenoids quantification.

**Supplementary Table 10.** Concentrated algal-produced sesquiterpenoids identified in ethanol using GCxGC-TOF/MS.

**Supplementary Table 11.** Sesquiterpenoids identified after hydroboration-oxidation reaction using GCxGC-TOF/MS.

**Supplementary Table 12.** Mass spectra of sesquiterpenoids identified by GC-MS.

### Supplementary Files

Files can be downloaded in the following link:

https://datadryad.org/stash/share/lSU89kHS9Poefgh9g65_v6Koerab9ZoBaYyGCcIri98

**Supplementary File 1.** GCxGC-TOF/MS data report of agarwood samples.

**Supplementary File 2.** GCxGC-TOF/MS raw data of agarwood samples.

**Supplementary File 3.** GC-TOF/MS raw data of agarwood samples.

**Supplementary File 4.** GC-MS/FID raw data of engineered algae strains.

**Supplementary File 5.** Genetic constructs.

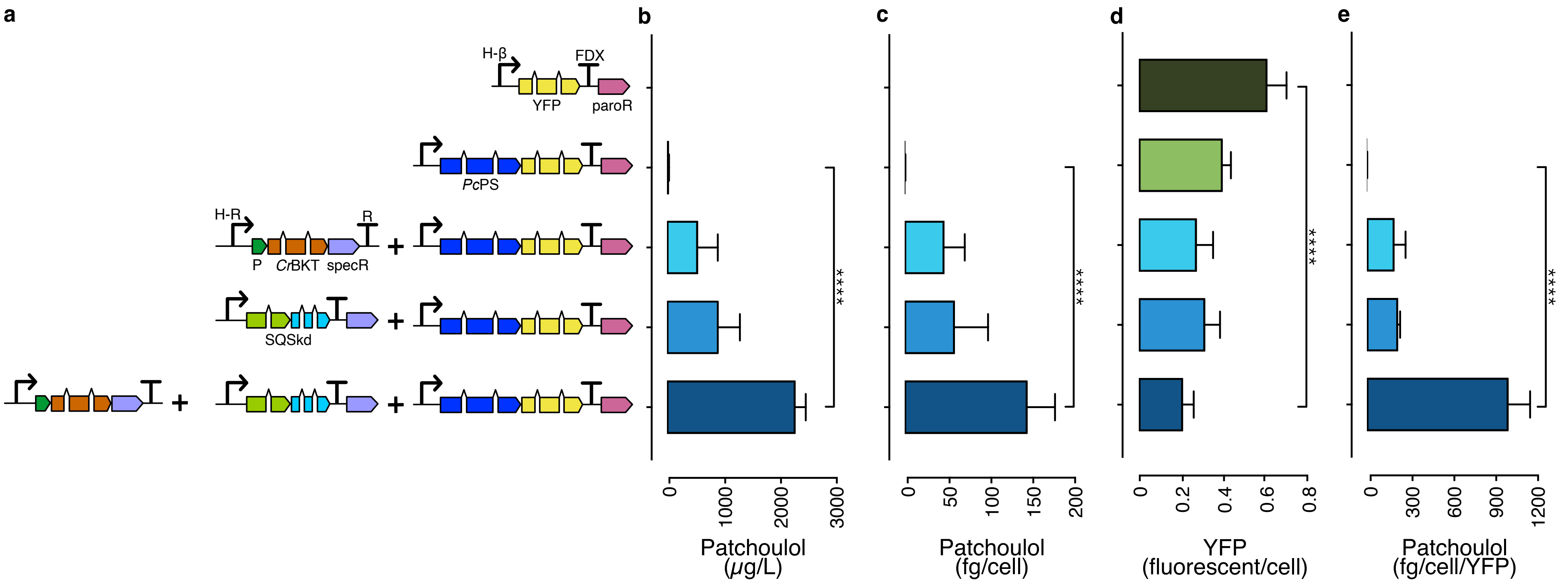


**Supplementary Figure 1.** Optimisation of *Chlamydomonas reinhardtii* metabolism for heterologous sesquiterpene production.

**a**. A variety of vectors, each containing algal-optimized genes such as mVenus (yellow fluorescent protein, YFP), PcPS-mVenus fusion, CrBKT, and CrSQS knockdown (k.d.), all integrated into the C. reinhardtii UPN22 parent strain. **b.** Patchoulol yields (µg/L) achieved with each engineered construct. **c.** Patchoulol yields (fg/cell) for each unique construct. **d.** Yellow fluorescent signal per cell for individual constructs. **e.** Patchoulol production for each construct normalised per cell by YFP fluorescence. **Definitions:** "k.d" refers to knock down; "*Pc*PS" stands for *Pogostemon cablin* patchoulol synthase; "*Cr*BKT" is *Chlamydomonas reinhardtii* β-carotene ketolase; "SQS" designates squalene synthase; "paroR" symbolises paromomycin resistance; "specR" identifies spectinomycin resistance; "H-β" represents the combined HSP70A-β tubulin promoter; "FDX" is the abbreviation for the ferredoxin 1 gene terminator; "P" indicates the photosystem I reaction centre subunit II. Statistical variations between groups were identified using ANOVA with Tukey's method for multiple comparisons. **Statistical significances:** *P < 0.05, **P < 0.01, ***P < 0.001. Comprehensive calculations, cell concentrations, and p-values are available in Supplementary Table 4.

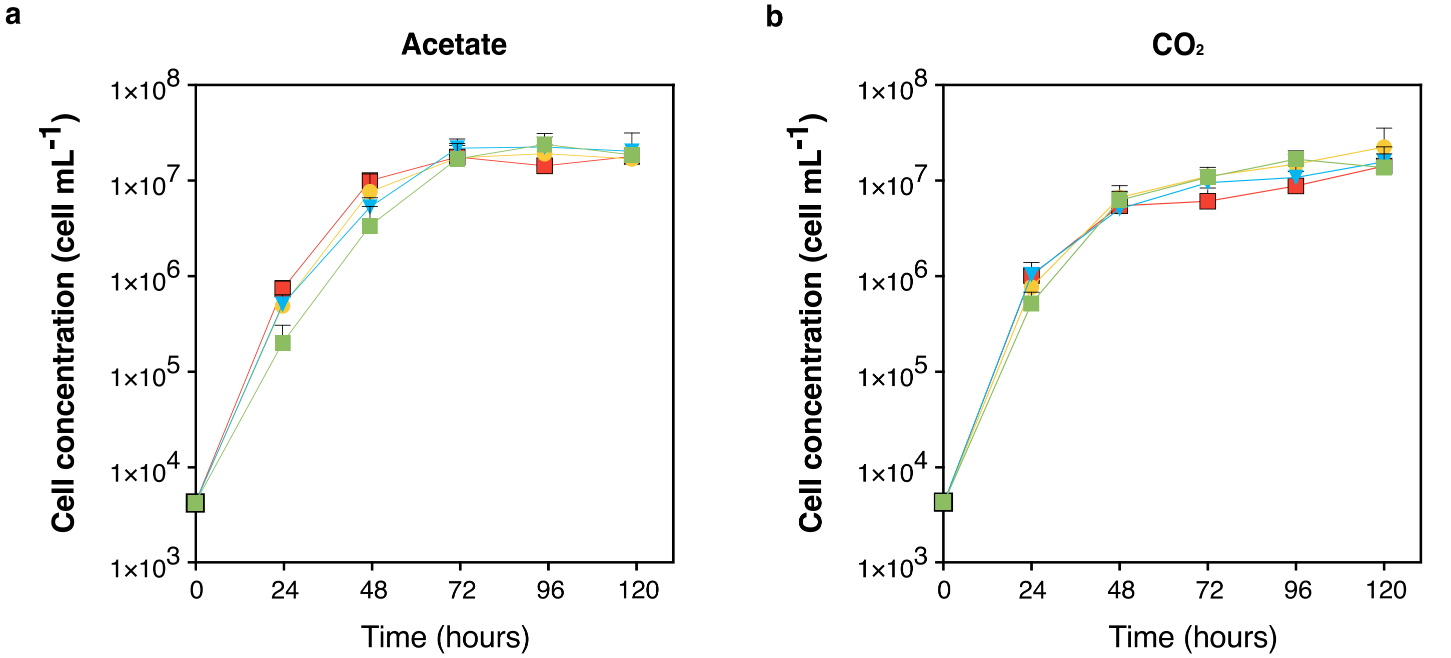


**Supplementary Figure 2.** Growth assessment of engineered *Chlamydomonas reinhardtii* engineered strains.

Analysis of cell concentration over a 120-hour period under two distinct carbon sources: (a) acetate, and (b) carbon dioxide ($CO_2$). Four transformants from each cell line were assessed and compared to the parental strain, *C. reinhardtii* UPN22. Sampling was performed every 24 hours using an automated liquid handler (Opentrons Labworks Inc., NY, USA), with cell counts determined by flow cytometry (Thermo Fisher Scientific, MA, USA). Statistical differences between the groups were identified through ANOVA, utilising Tukey's method for multiple comparisons. Significance levels are marked as follows: *$p < 0.05$, **$p < 0.01$, ***$p < 0.001$.

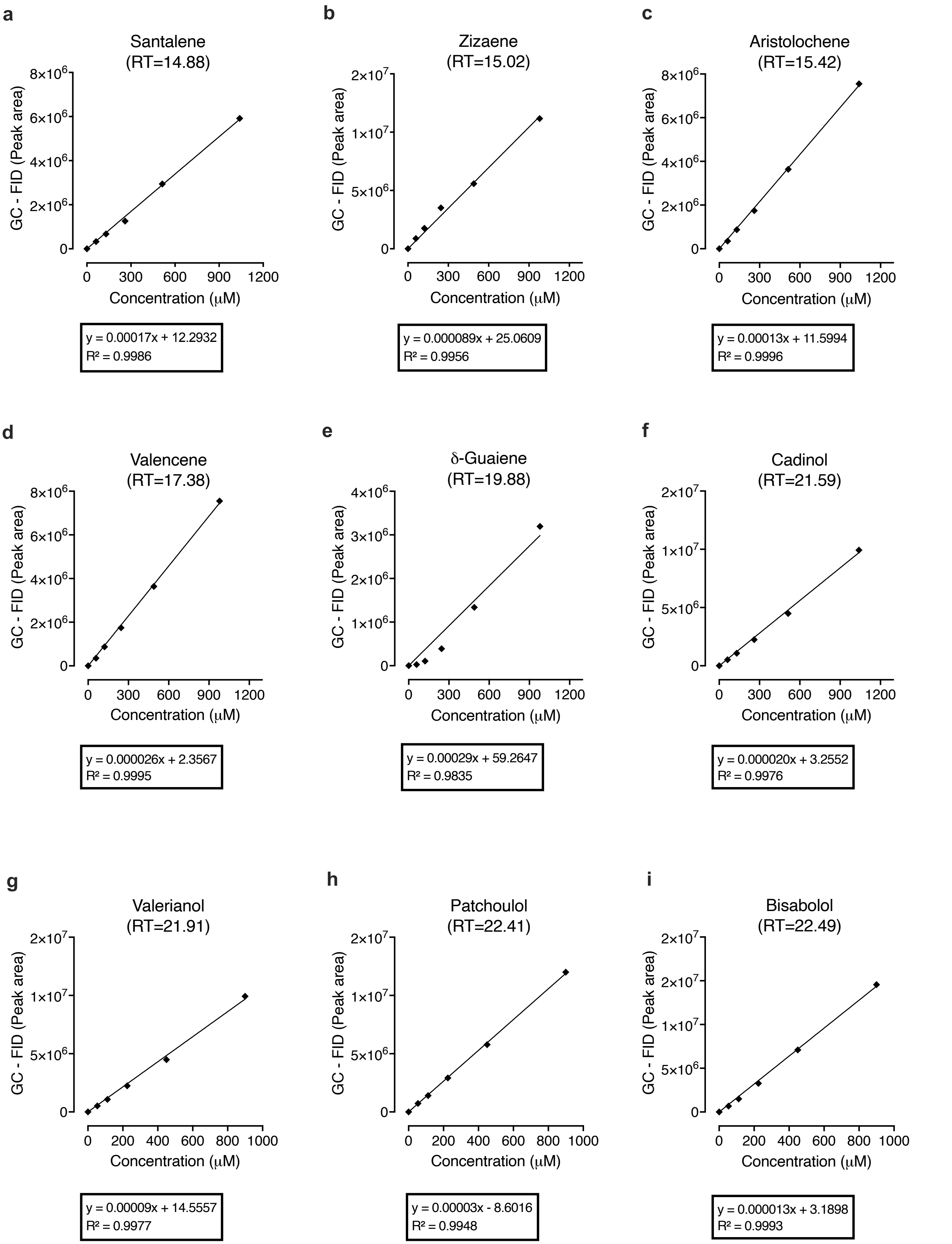


**Supplementary Figure 3**. Calibration curves of sesquiterpenoid standards.

Calibration curves and calculations utilised purified standards, with concentrations ranging from 1–1200 µM in dodecane, of the following sesquiterpenoids: δ-guaiene (CAT#B942760), patchoulol (CAT#P206200), santalene (CAT#S15065), valerianol (CAT#V914000, Toronto Research Chemicals, ON, Canada), bisabolol (CAT#95426), valencene (CAT#06808), and cedrene (CAT#22133, Sigma-Aldrich, MO, USA). For compound identification, retention time acquisition, internal digital library calibration, and method development, we used a set of 12 microampules containing a standard terpene mixture, which covered 98 terpenes at 1.0 mM in methanol (CAT# MSITPN101, MetaSci, ON, Canada).

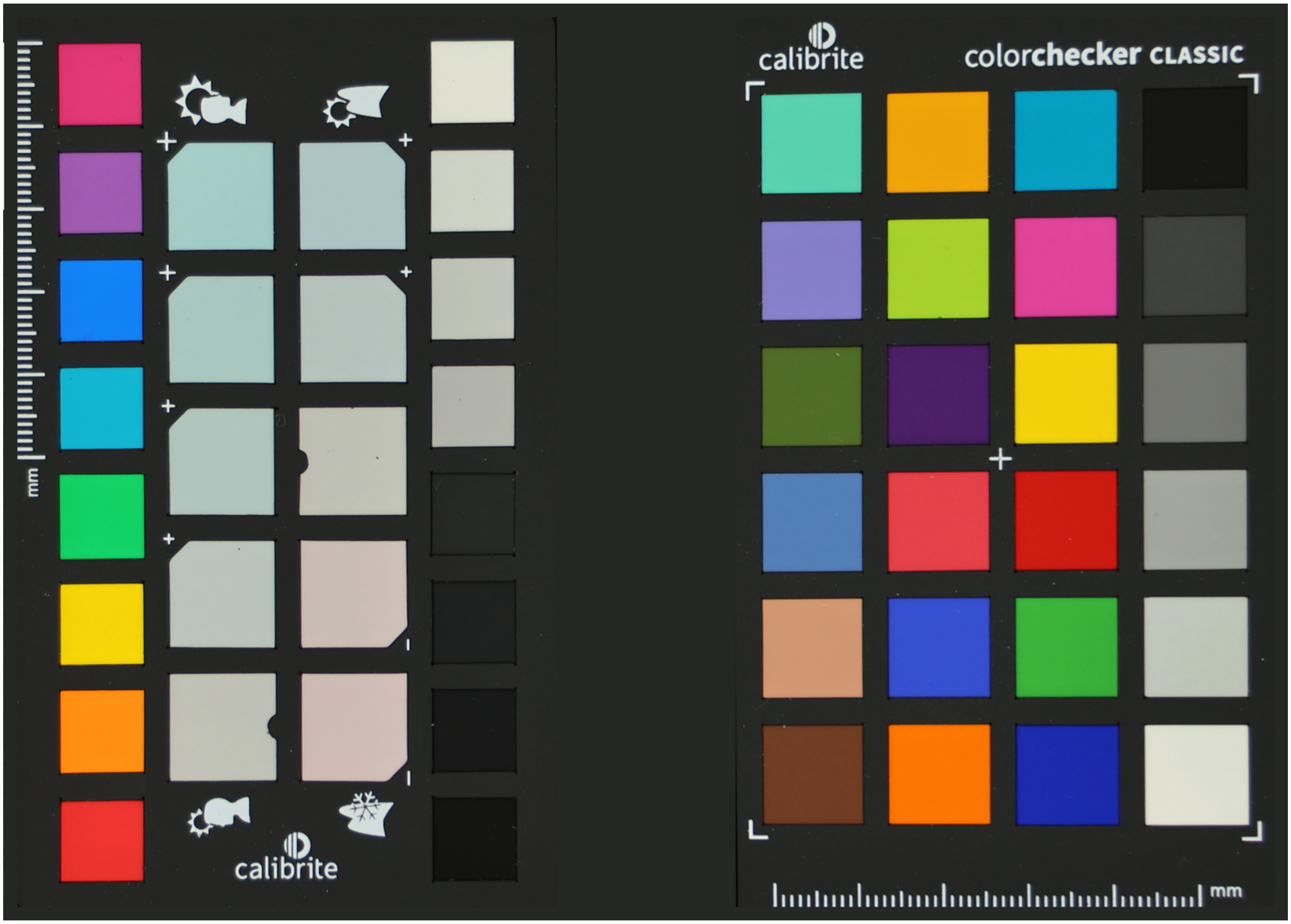


**Supplementary Figure 4.** Image of ColorChecker Passport.

Photograph of the ColorChecker Passport (CCPP2, Calibrite LLC, DE, USA) used for precise colour calibration. The calibration was achieved using the ColorChecker calibration software (Calibrite LLC, DE, USA) in conjunction with Adobe Lightroom (Adobe Inc., CA, USA) for colour correction profiling.

**Supplementary Table 1.** Agarwood samples description.

| No. Sample | Type | Origin (Country) | Code (Country) | Price (USD) (Kg or L) | | Total sesquiterpenoids (n=) | Non-functionalised sesquiterpenoids (n=) | Functionalised sesquiterpenoids (n=) |
|---|---|---|---|---|---|---|---|---|
| 1 | Agarwood | Philippines | PH | $ | 8,888 | 233 | 61 | 172 |
| 2 | Agarwood | Indonesia | ID | $ | 3,555 | 463 | 120 | 343 |
| 3 | Agarwood | Indonesia | ID | $ | 2,666 | 466 | 121 | 345 |
| 4 | Agarwood | Indonesia | ID | $ | 2,666 | 402 | 104 | 297 |
| 5 | Agarwood | Indonesia | ID | $ | 7,111 | 541 | 141 | 400 |
| 6 | Agarwood | Malaysia | MY | $ | 3,288 | 503 | 131 | 372 |
| 7 | Agarwood | Indonesia | ID | $ | 2,666 | 408 | 106 | 302 |
| 8 | Agarwood | Indonesia | ID | $ | 8,888 | 386 | 100 | 286 |
| 9 | Agarwood | Indonesia | ID | $ | 1,688 | 503 | 131 | 372 |
| 10 | Agarwood | Indonesia | ID | $ | 2,666 | 303 | 79 | 224 |
| 11 | Agarwood | Bangladesh | BD | $ | 2,666 | 303 | 79 | 224 |
| 12 | Agarwood | Malaysia | MY | $ | 5,333 | 381 | 99 | 282 |
| 13 | Agarwood | Indonesia | ID | $ | 3,555 | 323 | 84 | 239 |
| 14 | Agarwood | India | IN | $ | 2,666 | 558 | 145 | 413 |
| 15 | Agarwood | India | IN | $ | 8,888 | 294 | 76 | 218 |
| 16 | Agarwood | Thailand | TH | $ | 17,777 | 330 | 86 | 244 |
| 17 | Agarwood | Indonesia | ID | $ | 3,555 | 589 | 153 | 436 |
| 18 | Agarwood | Indonesia | ID | $ | 7,111 | 347 | 90 | 257 |
| 19 | Agarwood | Thailand | TH | $ | 3,288 | 382 | 99 | 283 |
| 20 | Agarwood | India | IN | $ | 2,666 | 382 | 99 | 283 |
| 21 | Agarwood | Indonesia | ID | $ | 21,422 | 477 | 124 | 353 |
| 22 | Agarwood | Indonesia | ID | $ | 1,777 | 369 | 96 | 273 |
| 23 | Agarwood | India | IN | $ | 3,555 | 449 | 117 | 332 |
| 24 | Agarwood | Indonesia | ID | $ | 3,555 | 446 | 116 | 330 |
| 25 | Agarwood | India | IN | $ | 1,777 | 389 | 101 | 287 |
| 26 | Agarwood | Indonesia | ID | $ | 2,666 | 470 | 122 | 348 |
| 27 | Agarwood | Indonesia | ID | $ | 1,333 | 477 | 124 | 353 |
| 28 | Agarwood | Indonesia | ID | $ | 2,666 | 470 | 122 | 348 |
| 29 | Agarwood | India | IN | $ | 1,777 | 426 | 111 | 315 |

**Supplementary Table 1.** (Cont.)

| No. Sample | Type | Origin (Country) | Code (Country) | Price (USD) (Kg or L) | Total sesquiterpenoids (n=) | Non-functionalised sesquiterpenoids (n=) | Functionalised sesquiterpenoids (n=) |
|---|---|---|---|---|---|---|---|
| 30 | Agarwood | India | IN | $ 2,666 | 464 | 121 | 343 |
| 31 | Agarwood | Indonesia | ID | $ 1,600 | 480 | 125 | 355 |
| 32 | Agarwood | Indonesia | ID | $ 2,222 | 340 | 88 | 252 |
| 33 | Agarwood | Indonesia | ID | $ 6,222 | 335 | 87 | 248 |
| 34 | Agarwood | Indonesia | ID | $ 1,777 | 477 | 124 | 353 |
| 35 | Agarwood | Thailand | TH | $ 2,666 | 323 | 84 | 239 |
| 36 | Agarwood | Vietnam | VN | $ 1,244 | 342 | 89 | 253 |
| 37 | Distillate | Bangladesh | BD | $ 8,888 | 497 | 159 | 388 |
| 38 | Distillate | Bangladesh | BD | $ 6,666 | 338 | 108 | 264 |
| 39 | Distillate | Indonesia | ID | $ 7,777 | 524 | 168 | 408 |
| 40 | Distillate | Cambodia | KH | $ 11,111 | 403 | 129 | 314 |
| 41 | Distillate | Indonesia | ID | $ 17,777 | 424 | 136 | 331 |
| 42 | Distillate | Bangladesh | BD | $ 11,111 | 584 | 187 | 455 |
| 43 | Distillate | Malaysia | MY | $ 17,777 | 582 | 186 | 454 |
| 44 | Distillate | Cambodia | KH | $ 8,888 | 629 | 201 | 491 |
| 45 | Distillate | India | IN | $ 11,111 | 756 | 242 | 590 |
| 46 | Distillate | Unknown | NU | $ 6,666 | 619 | 198 | 483 |
| 47 | Distillate | Unknown | NU | $ 8,888 | 672 | 215 | 524 |
| 48 | Distillate | Indonesia | ID | $ 13,333 | 585 | 187 | 456 |
| 49 | Distillate | Indonesia | ID | $ 13,333 | 519 | 166 | 405 |
| 50 | Distillate | Cambodia | KH | $ 8,888 | 314 | 101 | 245 |
| 51 | Distillate | Vietnam | VN | $ 8,888 | 247 | 79 | 193 |
| 52 | Distillate | Indonesia | ID | $ 6,666 | 605 | 194 | 472 |
| 53 | Distillate | Cambodia | KH | $ 8,888 | 619 | 198 | 483 |
| 54 | Distillate | India | IN | $ 17,777 | 436 | 140 | 340 |
| 55 | Distillate | Malaysia | MY | $ 17,777 | 569 | 182 | 444 |
| 56 | Distillate | India | IN | $ 11,111 | 156 | 50 | 122 |
| 57 | Distillate | Indonesia | ID | $ 15,555 | 136 | 44 | 106 |
| 58 | Distillate | Unknown | NU | $ 8,888 | 230 | 73 | 179 |

**Supplementary Table 2.** Sesquiterpenoids identified across agarwood (bakhour) samples using GCxGC-TOF/MS.

| Compound Name | Mol. formula | Clas. | RT I (min) | RT II (sec) | MF | P (%) |
|---|---|---|---|---|---|---|
| Aromadendrene | $C_{15}H_{22}$ | STP | 34.70 | 2.64 | 772 | 94 |
| α-Cubebene | $C_{15}H_{24}$ | STP | 34.70 | 2.22 | 714 | 68 |
| 4a,5-Dimethyl-3-(prop-1-en-2-yl)-1,2,3,4,4a,5,6,7-octahydronaphthalen-1-ol | $C_{15}H_{24}O$ | FSTP | 34.70 | 2.62 | 654 | 48 |
| β-Vatirenene | $C_{15}H_{22}$ | STP | 35.80 | 3.85 | 624 | 84 |
| Germacrene A | $C_{15}H_{24}$ | STP | 37.30 | 2.44 | 633 | 85 |
| β-copaene | $C_{15}H_{24}$ | STP | 37.30 | 2.42 | 674 | 92 |
| 1,5-Cyclodecadiene, 1,5-dimethyl-8-(1-methylethenyl)-, [S-(Z,E)]- | $C_{15}H_{24}$ | STP | 37.30 | 2.46 | 647 | 94 |
| α-gurjunene | $C_{15}H_{24}$ | STP | 38.20 | 2.66 | 641 | 65 |
| (2R,3R,3aR,6R,8aS)-3,7,7-Trimethyl-8-methyleneoctahydro-1H-3a,6-methanoazulen-2-ol | $C_{15}H_{24}O$ | FSTP | 38.70 | 3.17 | 654 | 82 |
| muuroladiene | $C_{15}H_{24}$ | STP | 39.10 | 2.76 | 659 | 75 |
| Caryophyllene | $C_{15}H_{24}$ | STP | 39.30 | 2.68 | 604 | 82 |
| Patchouladiene | $C_{15}H_{22}$ | STP | 39.50 | 3.04 | 643 | 84 |
| 1H-Cycloprop[e]azulene, decahydro-1,1,4,7-tetramethyl- | $C_{15}H_{26}$ | STP | 39.90 | 2.36 | 662 | 86 |
| 4b,5,6,7,8,8a,9,10-Octahydro-1-methylphenanthrene | $C_{15}H_{20}$ | FSTP | 40.20 | 3.31 | 786 | 90 |
| Cycloisolongifolol | $C_{15}H_{24}O$ | FSTP | 40.40 | 2.90 | 709 | 87 |
| β-GURJUNENE | $C_{15}H_{24}$ | STP | 40.80 | 2.74 | 791 | 90 |
| 4,7-Methanoazulene, 1,2,3,4,5,6,7,8-octahydro-1,4,9,9-tetramethyl-, [1S-(1α,4α,7α)]- | $C_{15}H_{24}$ | STP | 40.80 | 2.76 | 665 | 82 |
| Humulene | $C_{15}H_{24}$ | STP | 40.90 | 3.91 | 673 | 89 |
| β-Selinene | $C_{15}H_{24}$ | STP | 40.90 | 2.58 | 693 | 94 |
| α-bisabolene | $C_{15}H_{24}$ | STP | 41.30 | 2.92 | 603 | 90 |
| Alloaromadendrene | $C_{15}H_{24}$ | STP | 41.30 | 2.98 | 716 | 94 |
| Prezizaene | $C_{15}H_{24}$ | STP | 41.40 | 3.00 | 673 | 90 |
| Calamene | $C_{15}H_{22}$ | STP | 41.60 | 3.09 | 650 | 87 |
| 1,4,6-Trimethyl-1,2,3,3a,4,7,8,8a-octahydro-4,7-ethanoazulene | $C_{15}H_{24}$ | STP | 41.70 | 2.88 | 706 | 54 |
| 2-Methyl-3-(3-methyl-but-2-enyl)-2-(4-methyl-pent-3-enyl)-oxetane | $C_{15}H_{26}O$ | FSTP | 41.90 | 2.68 | 688 | 17 |
| Cadinene | $C_{15}H_{28}$ | STP | 42.10 | 2.38 | 514 | 65 |
| (3S,6S)-6-Isopropyl-3-methyl-2-(propan-2-ylidene)-3-vinylcyclohexanone | $C_{15}H_{24}O$ | FSTP | 42.30 | 3.09 | 720 | 89 |
| Diepicedrene oxide | $C_{15}H_{24}O$ | FSTP | 42.60 | 3.43 | 600 | 42 |

**Supplementary Table 2.** (Cont.)

| Compound Name | Mol. formula | Class. | RT I (min.) | RT II (sec.) | MF | P (%) |
|---|---|---|---|---|---|---|
| (R,Z)-2-Methyl-6-(4-methylcyclohexa-1,4-dien-1-yl)hept-2-en-1-ol | $C_{15}H_{24}O$ | FSTP | 42.70 | 3.11 | 610 | 73 |
| 1H-Cycloprop[e]azulene, decahydro-1,1,4,7-tetramethyl-, [1aR-(1aα,4β,4aβ,7β,7aβ,7bα)]- | $C_{15}H_{26}$ | STP | 42.90 | 2.62 | 708 | 81 |
| Caparratriene | $C_{15}H_{26}$ | STP | 43.10 | 2.60 | 700 | 83 |
| Cycloisolongifolene | $C_{15}H_{24}$ | STP | 43.10 | 2.98 | 677 | 91 |
| β-Guaiene | $C_{15}H_{24}$ | STP | 43.10 | 3.04 | 722 | 41 |
| Aristolochene | $C_{15}H_{24}$ | STP | 43.10 | 3.00 | 641 | 43 |
| Eremophilene | $C_{15}H_{24}$ | STP | 43.10 | 2.98 | 733 | 65 |
| α-himachalene | $C_{15}H_{20}$ | FSTP | 43.30 | 3.59 | 777 | 41 |
| β-Bisabolenol | $C_{15}H_{24}O$ | FSTP | 43.30 | 3.21 | 572 | 79 |
| Eudesmatriene | $C_{15}H_{22}$ | STP | 43.40 | 3.17 | 825 | 72 |
| 2-(4a,8-Dimethyl-2,3,4,5,6,7-hexahydro-1H-naphthalen-2-yl)propan-2-ol | $C_{15}H_{26}O$ | FSTP | 43.40 | 2.94 | 752 | 76 |
| 10,11-Epoxycalamenene | $C_{15}H_{20}O$ | FSTP | 43.40 | 3.77 | 753 | 61 |
| β-Vetispirene | $C_{15}H_{22}$ | STP | 43.50 | 3.17 | 588 | 38 |
| (2R,8R,8aS)-8,8a-Dimethyl-2-(prop-1-en-2-yl)-1,2,3,7,8,8a-hexahydronaphthalene | $C_{15}H_{22}$ | STP | 43.50 | 3.13 | 752 | 56 |
| 6-Methyl-2-(4-methylcyclohex-3-en-1-yl)hepta-1,5-dien-4-ol | $C_{15}H_{24}O$ | FSTP | 43.70 | 4.27 | 708 | 38 |
| 1H-3a,7-Methanoazulene, octahydro-1,4,9,9-tetramethyl- | $C_{15}H_{26}$ | STP | 43.90 | 2.64 | 709 | 67 |
| (3R,3aR,4aR,8aR,9aR)-3,8a-Dimethyl-5-methylene-3,3a,4,4a,5,6,9,9a-octahydronaphtho[2,3-b]furan-2(8aH)-one | $C_{15}H_{20}O_2$ | FSTP | 44.80 | 4.70 | 816 | 42 |
| (3R,3aR,3bR,4S,7R,7aR)-4-Isopropyl-3,7-dimethyloctahydro-1H-cyclopenta[1,3]cyclopropa[1,2]benzen-3-ol | $C_{15}H_{26}O$ | FSTP | 44.90 | 2.98 | 824 | 84 |
| Nivalenol | $C_{15}H_{20}O_7$ | FSTP | 45.20 | 3.55 | 769 | 61 |
| Calamenene | $C_{15}H_{22}$ | STP | 45.30 | 3.41 | 625 | 80 |
| Kessane | $C_{15}H_{26}O$ | FSTP | 45.80 | 3.37 | 959 | 71 |
| Cubenene | $C_{15}H_{24}$ | STP | 45.80 | 3.08 | 449 | 77 |
| 1-Formyl-2,2-dimethyl-3-cis-(2-methyl-but-2-enyl)-6-methylidene-cyclohexane | $C_{15}H_{24}O$ | FSTP | 46.00 | 3.85 | 931 | 92 |
| 2,6-Dimethyl-10-methylene-12-oxatricyclo[7.3.1.0(1,6)]tridec-2-ene | $C_{15}H_{22}O$ | FSTP | 46.20 | 3.61 | 829 | 81 |
| α-Calacorene | $C_{15}H_{20}$ | FSTP | 46.50 | 3.73 | 737 | 84 |
| Nerolidol | $C_{15}H_{26}O$ | FSTP | 46.80 | 3.97 | 709 | 70 |
| α-agorofuran | $C_{15}H_{24}O$ | FSTP | 46.90 | 3.99 | 735 | 71 |
| 2-Pentadecen-4-yne, (Z)- | $C_{15}H_{26}$ | STP | 46.90 | 5.22 | 720 | 51 |

**Supplementary Table 2.** (Cont.)

| Compound Name | Mol. formula | Class. | RT I (min.) | RT II (sec.) | MF | P (%) |
|---|---|---|---|---|---|---|
| Longifolene | $C_{15}H_{20}$ | FSTP | 47.20 | 3.69 | 728 | 62 |
| Aromadendran | $C_{15}H_{26}$ | STP | 47.40 | 4.25 | 754 | 77 |
| Dihydroagarofurane | $C_{15}H_{26}O$ | FSTP | 47.50 | 3.47 | 754 | 90 |
| 4-Isopropyl-6-methyl-1-methylene-1,2,3,4-tetrahydronaphthalene | $C_{15}H_{20}$ | FSTP | 47.70 | 3.85 | 749 | 88 |
| 4(15)-Selinene-11,12-diol | $C_{15}H_{26}O_2$ | FSTP | 47.90 | 4.58 | 875 | 41 |
| 2-(4a,8-Dimethyl-2,3,4,4a,5,6-hexahydro-naphthalen-2-yl)-prop-2-en-1-ol | $C_{15}H_{22}O$ | FSTP | 48.20 | 4.86 | 774 | 81 |
| Lanceol | $C_{15}H_{24}O$ | FSTP | 48.60 | 3.55 | 779 | 72 |
| (5S,6R,7S,10R)-7-Isopropyl-2,10-dimethylspiro[4.5]dec-1-en-6-ol | $C_{15}H_{26}O$ | FSTP | 48.80 | 3.27 | 755 | 55 |
| Neoisolongifolene | $C_{15}H_{22}$ | STP | 49.30 | 2.72 | 773 | 56 |
| α-Santalol | $C_{15}H_{24}O$ | FSTP | 49.50 | 4.72 | 768 | 43 |
| 3,7-Cycloundecadien-1-ol, 1,5,5,8-tetramethyl- | $C_{15}H_{26}O$ | FSTP | 49.90 | 3.69 | 729 | 44 |
| isoledene | $C_{15}H_{24}$ | STP | 50.10 | 3.47 | 672 | 36 |
| β-Oplopenone | $C_{15}H_{24}O$ | FSTP | 50.10 | 3.49 | 803 | 60 |
| Eudesmadien | $C_{15}H_{24}O$ | FSTP | 50.20 | 3.69 | 869 | 80 |
| Taylorione | $C_{15}H_{22}O$ | FSTP | 50.50 | 3.95 | 594 | 40 |
| Humulene epoxide I | $C_{15}H_{24}O$ | FSTP | 50.50 | 3.87 | 559 | 80 |
| (1R,4S,5S)-1,8-Dimethyl-4-(prop-1-en-2-yl)spiro[4.5]dec-7-ene | $C_{15}H_{24}$ | STP | 51.00 | 4.03 | 579 | 90 |
| 2-((2S,4aR)-4a,8-Dimethyl-1,2,3,4,4a,5,6,7-octahydronaphthalen-2-yl)propan-2-ol | $C_{15}H_{26}O$ | FSTP | 51.10 | 3.75 | 509 | 60 |
| ɣ-himachalene | $C_{15}H_{24}$ | STP | 51.10 | 4.17 | 564 | 81 |
| 4a(2H)-Naphthalenol, 1,3,4,5,6,8a-hexahydro-4,7-dimethyl-1-(1-methylethyl)-, (1S,4R,4aS,8aR)- | $C_{15}H_{26}O$ | FSTP | 51.40 | 3.51 | 584 | 91 |
| 2(1H)-Naphthalenone, 4a,5,6,7,8,8a-hexahydro-7α-isopropyl-4aβ,8aβ-dimethyl- | $C_{15}H_{24}O$ | FSTP | 51.90 | 3.43 | 662 | 91 |
| δ-Guaiene | $C_{15}H_{24}$ | STP | 52.00 | 3.75 | 669 | 93 |
| β-Caryophyllene | $C_{15}H_{24}$ | STP | 52.10 | 4.09 | 574 | 32 |
| **τ**-Cadinol | $C_{15}H_{26}O$ | FSTP | 52.10 | 3.71 | 640 | 42 |
| Pogostol | $C_{15}H_{26}O$ | FSTP | 52.20 | 3.79 | 574 | 61 |
| 1H-Cycloprop[e]azulen-4-ol, decahydro-1,1,4,7-tetramethyl-, [1aR-(1aα,4β,4aβ,7α,7aβ,7bα)]- | $C_{15}H_{26}O$ | FSTP | 52.20 | 3.75 | 494 | 51 |
| α-Guaiene | $C_{15}H_{24}$ | STP | 52.30 | 3.85 | 641 | 72 |
| 5β,7βH,10α-Eudesm-11-en-1α-ol | $C_{15}H_{26}O$ | FSTP | 52.40 | 0.42 | 651 | 48 |

**Supplementary Table 2.** (Cont.)

| Compound Name | Mol. formula | Class. | RT I (min.) | RT II (sec.) | MF | P (%) |
|---|---|---|---|---|---|---|
| 10s,11s-Himachala-3(12),4-diene | $C_{15}H_{24}$ | STP | 52.40 | 3.85 | 861 | 41 |
| Rosifoliol | $C_{15}H_{26}O$ | FSTP | 52.50 | 4.05 | 591 | 51 |
| Aristolene | $C_{15}H_{24}$ | STP | 52.50 | 4.19 | 731 | 58 |
| Agarospirol | $C_{15}H_{26}O$ | FSTP | 52.80 | 4.07 | 525 | 37 |
| Hedycaryol | $C_{15}H_{26}O$ | FSTP | 52.80 | 3.99 | 592 | 71 |
| Maaliol | $C_{15}H_{26}O$ | FSTP | 52.80 | 4.09 | 690 | 56 |
| Globulol | $C_{15}H_{26}O$ | FSTP | 52.80 | 4.31 | 634 | 59 |
| Cubenol | $C_{15}H_{26}O$ | FSTP | 52.90 | 4.13 | 671 | 66 |
| 2-Naphthalenol, 2,3,4,4a,5,6,7-octahydro-1,4a-dimethyl-7-(2-hydroxy-1-methylethyl) | $C_{15}H_{26}O_2$ | FSTP | 52.90 | 4.52 | 593 | 77 |
| β-BERGAMOTENE | $C_{15}H_{24}$ | STP | 52.90 | 4.48 | 700 | 58 |
| Ledol | $C_{15}H_{26}O$ | FSTP | 52.90 | 5.81 | 735 | 65 |
| 1,2-Naphthalenedione, 3,8-dimethyl-5-(1-methylethyl)- | $C_{15}H_{16}O_2$ | FSTP | 53.00 | 4.58 | 835 | 31 |
| Guaiol | $C_{15}H_{26}O$ | FSTP | 53.00 | 4.09 | 672 | 68 |
| (1S,7S,8aR)-1,8a-Dimethyl-7-(prop-1-en-2-yl)-1,2,3,7,8,8a-hexahydronaphthalene | $C_{15}H_{22}$ | STP | 53.00 | 4.38 | 758 | 95 |
| α-Bisabolol oxide B | $C_{15}H_{26}O_2$ | FSTP | 53.00 | 3.55 | 701 | 77 |
| Terranol | $C_{15}H_{26}O$ | FSTP | 53.10 | 5.28 | 595 | 77 |
| β-Longipinene | $C_{15}H_{24}$ | STP | 53.20 | 4.33 | 704 | 78 |
| Cedrol | $C_{15}H_{26}O$ | FSTP | 53.30 | 4.33 | 712 | 91 |
| α-coastal | $C_{15}H_{22}O$ | FSTP | 53.50 | 4.50 | 892 | 61 |
| Zonarene | $C_{15}H_{24}$ | STP | 53.60 | 5.00 | 685 | 70 |
| 5-Azulenemethanol, 1,2,3,3a,4,5,6,7-octahydro-α,α,3,8-tetramethyl-, [3S-(3α,3aβ,5α)]- | $C_{15}H_{26}O$ | FSTP | 53.60 | 3.93 | 789 | 20 |
| α-copaene | $C_{15}H_{24}O$ | FSTP | 53.90 | 4.44 | 552 | 41 |
| Valerenal | $C_{15}H_{22}O$ | FSTP | 54.00 | 4.58 | 735 | 61 |
| Sesquisabinene | $C_{15}H_{26}O$ | FSTP | 54.10 | 3.89 | 612 | 20 |
| (3bR,4S,7aS)-7,7,8,8-Tetramethyloctahydro-2,3b-methanocyclopenta[1,3]cyclopropa[1,2]benzen-4-ol | $C_{15}H_{24}O$ | FSTP | 54.10 | 4.27 | 619 | 21 |
| 1-Buten-1-ol, 2-methyl-4-(2,6,6-trimethyl-1-cyclohexen-1-yl)-, formate, (E)- | $C_{15}H_{24}O_2$ | FSTP | 54.30 | 4.78 | 558 | 25 |
| α.-Eudesmol | $C_{15}H_{24}O$ | FSTP | 54.60 | 4.21 | 641 | 31 |
| Valerenol | $C_{15}H_{24}O$ | FSTP | 54.70 | 4.36 | 537 | 61 |

**Supplementary Table 2.** (Cont.)

| Compound Name | Mol. formula | Class. | RT I (min.) | RT II (sec.) | MF | P (%) |
|---|---|---|---|---|---|---|
| α-isocomene | $C_{15}H_{24}$ | STP | 54.90 | 4.21 | 589 | 57 |
| Acorenone B | $C_{15}H_{24}O$ | FSTP | 54.90 | 4.23 | 625 | 44 |
| Valencene | $C_{15}H_{24}$ | STP | 54.90 | 4.21 | 602 | 49 |
| Carvyl angelate | $C_{15}H_{22}O_2$ | FSTP | 55.00 | 3.77 | 641 | 34 |
| Spathulenol | $C_{15}H_{24}O$ | FSTP | 55.00 | 4.33 | 641 | 61 |
| Widdrenal | $C_{15}H_{22}O$ | FSTP | 55.00 | 4.42 | 605 | 72 |
| Germacratrienol | $C_{15}H_{24}O$ | FSTP | 55.10 | 4.36 | 658 | 40 |
| Epiprocurcumenol | $C_{15}H_{22}O$ | FSTP | 55.20 | 4.09 | 619 | 11 |
| 2,2,6-Trimethyl-1-(2-methyl-cyclobut-2-enyl)-hepta-4,6-dien-3-one | $C_{15}H_{22}O$ | FSTP | 55.20 | 4.09 | 694 | 61 |
| 2-((3R,3aR,3bS,4R,7R,7aS)-3,7-Dimethyloctahydro-1H-cyclopenta[1,3]cyclopropa[1,2]benzen-4-yl)propan-2-ol | $C_{15}H_{26}O$ | FSTP | 55.90 | 3.81 | 592 | 91 |
| Solstitialin A | $C_{15}H_{20}O_5$ | FSTP | 56.00 | 4.82 | 714 | 59 |
| Calarene epoxide | $C_{15}H_{24}O$ | FSTP | 56.10 | 5.24 | 725 | 51 |
| α-Santalol | $C_{15}H_{24}O$ | FSTP | 56.10 | 4.05 | 760 | 42 |
| α-cyperone | $C_{15}H_{20}O$ | FSTP | 56.60 | 4.38 | 459 | 55 |
| 2,2,5-Trimethyl-1-phenylhexa-3,4-dien-1-one | $C_{15}H_{18}O$ | FSTP | 56.80 | 4.82 | 627 | 64 |
| 6-Dehydropetasol | $C_{15}H_{20}O_2$ | FSTP | 56.80 | 4.86 | 683 | 41 |
| (R)-1,5,8-Trimethyl-6,7,8,9-tetrahydronaphtho[2,1-b]furan | $C_{15}H_{18}O$ | FSTP | 57.20 | 4.72 | 714 | 4 |
| Cycloseychellene | $C_{15}H_{24}$ | STP | 57.20 | 5.59 | 665 | 7 |
| Elemol | $C_{15}H_{26}O$ | FSTP | 57.30 | 5.73 | 668 | 20 |
| Thujopsene-I3 | $C_{15}H_{24}$ | STP | 57.40 | 5.93 | 722 | 8 |
| α-Hexylcinnamaldehyde | $C_{15}H_{20}O$ | FSTP | 57.50 | 4.44 | 726 | 20 |
| Vetiselinenol | $C_{15}H_{24}O$ | FSTP | 57.50 | 4.52 | 692 | 4 |
| Eudesmenol | $C_{15}H_{26}O$ | FSTP | 57.60 | 5.46 | 652 | 6 |
| Khusimyl acid | $C_{15}H_{22}O$ | FSTP | 57.90 | 4.86 | 585 | 6 |
| 7-Isopropenyl-1,4a-dimethyl-4,4a,5,6,7,8-hexahydro-3H-naphthalen-2-one | $C_{15}H_{22}O$ | FSTP | 58.00 | 4.64 | 669 | 10 |
| 3,7-Cyclodecadien-1-one, 3,7-dimethyl-10-(1-methylethylidene)-, (E,E)- | $C_{15}H_{22}O$ | FSTP | 58.20 | 5.04 | 721 | 15 |
| 5(1H)-Azulenone, 2,4,6,7,8,8a-hexahydro-3,8-dimethyl-4-(1-methylethylidene)-, (8S-cis)- | $C_{15}H_{22}O$ | FSTP | 58.30 | 5.63 | 615 | 11 |
| Valerenic acid | $C_{15}H_{22}O_2$ | FSTP | 58.60 | 5.73 | 741 | 9 |

**Supplementary Table 2.** (Cont.)

| Compound Name | Mol. formula | Class. | RT I (min.) | RT II (sec.) | MF | P (%) |
|---|---|---|---|---|---|---|
| 6-Isopropenyl-4,8a-dimethyl-4a,5,6,7,8,8a-hexahydro-1H-naphthalen-2-one | $C_{15}H_{22}O$ | FSTP | 58.70 | 5.14 | 676 | 13 |
| 1H-3a,7-Methanoazulene-6-methanol, 2,3,4,7,8,8a-hexahydro-3,8,8-trimethyl-, [3R-(3α,3aβ,7β,8aα)]- | $C_{15}H_{24}O$ | FSTP | 58.80 | 0.69 | 729 | 12 |
| 4-(3,3-Dimethyl-but-1-ynyl)-4-hydroxy-2,6,6-trimethylcyclohex-2-enone | $C_{15}H_{22}O_2$ | FSTP | 58.90 | 5.32 | 687 | 4 |
| (S,Z)-2-Methyl-6-(p-tolyl)hept-2-en-1-ol | $C_{15}H_{22}O$ | FSTP | 58.90 | 4.94 | 677 | 9 |
| 7-(1,3-Dimethylbuta-1,3-dienyl)-1,6,6-trimethyl-3,8-dioxatricyclo[5.1.0.0(2,4)]octane | $C_{15}H_{22}O_2$ | FSTP | 59.40 | 4.98 | 652 | 6 |
| Valerenic acid | $C_{15}H_{22}O_2$ | FSTP | 58.60 | 5.73 | 741 | 9 |
| 6-Isopropenyl-4,8a-dimethyl-4a,5,6,7,8,8a-hexahydro-1H-naphthalen-2-one | $C_{15}H_{22}O$ | FSTP | 58.70 | 5.14 | 676 | 13 |
| 1H-3a,7-Methanoazulene-6-methanol, 2,3,4,7,8,8a-hexahydro-3,8,8-trimethyl-, [3R-(3α,3aβ,7β,8aα)]- | $C_{15}H_{24}O$ | FSTP | 58.80 | 0.69 | 729 | 12 |
| γ-Gurjunenepoxide | $C_{15}H_{24}O$ | FSTP | 59.50 | 4.64 | 690 | 19 |
| (E,Z)-α-Farnesene | $C_{15}H_{24}$ | STP | 59.80 | 0.26 | 657 | 8 |
| 1(2H)Phenanthrenone, 3,4,4a,9,10,10a-hexahydro-4a-methyl- | $C_{15}H_{18}O$ | FSTP | 60.00 | 4.80 | 759 | 10 |
| (R)-3-Methylene-6-((S)-1,2,2-trimethylcyclopentyl)cyclohex-1-ene | $C_{15}H_{24}$ | STP | 60.20 | 5.95 | 763 | 5 |
| 1H-Benzocyclohepten-7-ol, 2,3,4,4a,5,6,7,8-octahydro-1,1,4a,7-tetramethyl-, cis- | $C_{15}H_{26}O$ | FSTP | 60.30 | 5.93 | 674 | 12 |
| Ylangenal | $C_{15}H_{22}O$ | FSTP | 60.40 | 5.77 | 516 | 22 |
| Proximadiol | $C_{15}H_{28}O_2$ | FSTP | 60.60 | 5.16 | 711 | 23 |
| 3H-Cyclodeca[b]furan-2-one, 4,9-dihydroxy-6-methyl-3,10-dimethylene-3a,4,7,8,9,10,11,11a-octahydro- | $C_{15}H_{20}O_4$ | FSTP | 60.80 | 4.70 | 666 | 10 |
| Nootkatone | $C_{15}H_{22}O$ | FSTP | 61.00 | 5.28 | 740 | 12 |
| β-acoradienol | $C_{15}H_{24}O$ | FSTP | 61.00 | 4.23 | 689 | 29 |
| 2H-Cyclopropa[a]naphthalen-2-one, 1,1a,4,5,6,7,7a,7b-octahydro-1,1,7,7a-tetramethyl-, (1aα,7α,7aα,7bα)- | $C_{15}H_{22}O$ | FSTP | 61.00 | 5.20 | 703 | 29 |
| Dehydrofukinone | $C_{15}H_{22}O$ | FSTP | 61.10 | 5.38 | 674 | 41 |
| Caryophylladienol | $C_{15}H_{26}O$ | FSTP | 61.10 | 6.01 | 667 | 16 |
| Elemadiol | $C_{15}H_{26}O_2$ | FSTP | 61.30 | 4.82 | 656 | 14 |
| β-Eudesmol | $C_{15}H_{26}O$ | FSTP | 61.60 | 5.12 | 656 | 12 |
| 4,4-Dimethyl-3-(3-methylbut-2-enylidene)octane-2,7-dione | $C_{15}H_{24}O_2$ | FSTP | 61.70 | 5.63 | 584 | 7 |
| 4,6,6-Trimethyl-2-(3-methylbuta-1,3-dienyl)-3-oxatricyclo[5.1.0.0(2,4)]octane | $C_{15}H_{22}O$ | FSTP | 61.90 | 6.09 | 609 | 15 |
| Cyperotundone | $C_{15}H_{22}O$ | FSTP | 62.10 | 4.88 | 677 | 7 |
| 2-Cyclohexene-1-carboxaldehyde, 2,6-dimethyl-6-(4-methyl-3-pentenyl)- | $C_{15}H_{24}O$ | FSTP | 62.20 | 4.33 | 618 | 25 |
| 6-Isopropenyl-4,8a-dimethyl-1,2,3,5,6,7,8,8a-octahydro-naphthalen-2-ol | $C_{15}H_{24}O$ | FSTP | 62.60 | 1.96 | 666 | 18 |

**Supplementary Table 2.** (Cont.)

| Compound Name | Mol. formula | Class. | RT I (min.) | RT II (sec.) | MF | P (%) |
|---|---|---|---|---|---|---|
| Hexahydronaphthalenone | $C_{15}H_{22}O$ | FSTP | 62.60 | 5.83 | 708 | 11 |
| Isovelleral | $C_{15}H_{20}O_2$ | FSTP | 62.70 | 5.77 | 572 | 94 |
| 6-(p-Tolyl)-2-methyl-2-heptenol, trans- | $C_{15}H_{22}O$ | FSTP | 62.70 | 0.99 | 675 | 18 |
| 1(2H)-Naphthalenone, octahydro-4,8a-dimethyl-6-(1-methylethenyl)-, (4α,4aβ,6α,8aβ)- | $C_{15}H_{24}O$ | FSTP | 63.00 | 0.87 | 608 | 65 |
| Alloaromadendrene oxide | $C_{15}H_{24}O$ | FSTP | 63.00 | 5.08 | 639 | 10 |
| Uvidin C | $C_{15}H_{26}O_2$ | FSTP | 63.10 | 5.85 | 553 | 13 |
| Costal | $C_{15}H_{22}O$ | FSTP | 63.10 | 5.95 | 590 | 15 |
| 1,1,4,7-Tetramethyl-1a,2,3,4,6,7,7a,7b-octahydro-1H-cyclopropa[e]azulene | $C_{15}H_{24}$ | STP | 63.20 | 5.08 | 677 | 17 |
| α-Farnesene | $C_{15}H_{24}$ | STP | 63.50 | 0.40 | 659 | 15 |
| Isoaromadendrene epoxide | $C_{15}H_{24}O$ | FSTP | 63.50 | 5.24 | 529 | 32 |
| 2-((2R,4aR,8aR)-4a,8-Dimethyl-1,2,3,4,4a,5,6,8a-octahydronaphthalen-2-yl)prop-2-en-1-ol | $C_{15}H_{24}O$ | FSTP | 63.50 | 5.20 | 580 | 40 |
| 4-Isopropyl-6-methyl-3,4,4a,7,8,8a-hexahydronaphthalene-1-carbaldehyde | $C_{15}H_{22}O$ | FSTP | 63.50 | 5.20 | 547 | 15 |
| Bergamoto | $C_{15}H_{24}O$ | FSTP | 63.80 | 0.73 | 607 | 23 |
| Silphiperfola-4,7(14)-diene | $C_{15}H_{22}$ | STP | 63.80 | 0.65 | 504 | 25 |
| 1-Cyclopropene-1-pentanol, α,ε,ε,2-tetramethyl-3-(1-methylethenyl)- | $C_{15}H_{26}O$ | FSTP | 63.90 | 5.54 | 698 | 15 |
| Ledene oxide | $C_{15}H_{24}O$ | FSTP | 64.30 | 0.77 | 552 | 7 |
| Carabrol | $C_{15}H_{22}O_3$ | FSTP | 64.30 | 4.72 | 549 | 14 |
| ɣ-gurjunene | $C_{15}H_{24}$ | STP | 64.30 | 5.20 | 534 | 26 |
| 10-Epigazaniolide | $C_{15}H_{18}O_2$ | FSTP | 64.40 | 1.55 | 593 | 15 |
| 14-Hydroxycaryophyllene | $C_{15}H_{24}O$ | FSTP | 64.40 | 5.65 | 557 | 9 |
| 7-Tetracyclo[6.2.1.0(3.8)0(3.9)]undecanol, 4,4,11,11-tetramethyl- | $C_{15}H_{24}O$ | FSTP | 64.80 | 5.87 | 562 | 24 |
| Isospathulenol | $C_{15}H_{24}O$ | FSTP | 65.50 | 5.50 | 653 | 29 |
| 2,4a,5,8a-Tetramethyl-1,2,3,4,4a,7,8,8a-octahydronaphthalen-1-yl formate (isomer 1) | $C_{15}H_{24}O_2$ | FSTP | 65.50 | 0.52 | 547 | 11 |
| Cumanin | $C_{15}H_{22}O_4$ | FSTP | 66.50 | 1.88 | 528 | 20 |
| 11,11-Dimethyl-4,8-dimethylenebicyclo[7.2.0]undecan-3-ol | $C_{15}H_{24}O$ | FSTP | 66.70 | 0.58 | 598 | 5 |
| (2S,4aR,8aR)-4a,8-Dimethyl-2-(prop-1-en-2-yl)-1,2,3,4,4a,5,6,8a-octahydronaphthalene | $C_{15}H_{24}$ | STP | 66.80 | 0.65 | 549 | 46 |
| α-Panasinsen | $C_{15}H_{24}$ | STP | 66.80 | 0.62 | 547 | 26 |
| 3-Cyclohexene-1-ethanol, α-ethenyl-α,3-dimethyl-6-(1-methylethylidene)- | $C_{15}H_{24}O$ | FSTP | 66.90 | 5.56 | 556 | 28 |

**Supplementary Table 2.** (Cont.)

| Compound Name | Mol. formula | Class. | RT I (min.) | RT II (sec.) | MF | P (%) |
|---|---|---|---|---|---|---|
| 2,3,3-Trimethyl-2-(3-methylbuta-1,3-dienyl)-6-methylenecyclohexanone | $C_{15}H_{22}O$ | FSTP | 66.90 | 6.19 | 686 | 19 |
| Jaeskeanadiol | $C_{15}H_{26}O_2$ | FSTP | 67.00 | 1.09 | 590 | 60 |
| Parthenolide | $C_{15}H_{20}O_3$ | FSTP | 67.00 | 6.13 | 614 | 31 |
| Valerenolic acid | $C_{15}H_{22}O_3$ | FSTP | 67.00 | 5.57 | 556 | 91 |
| 7-(hydroxymethyl)-2,2,4-trimethyl-3,3a,4,8a-tetrahydro-1H-azulen-6-yl]methanol | $C_{15}H_{24}O_2$ | FSTP | 67.10 | 6.13 | 578 | 29 |
| 7-Oxabicyclo[4.1.0]heptane, 2,2,6-trimethyl-1-(3-methyl-1,3-butadienyl)-5-methylene- | $C_{15}H_{22}O$ | FSTP | 67.10 | 6.11 | 583 | 26 |
| α-Bisabolene epoxide | $C_{15}H_{24}O$ | FSTP | 67.30 | 6.21 | 591 | 50 |
| Aromadendrene oxide | $C_{15}H_{24}O$ | FSTP | 67.30 | 5.85 | 610 | 66 |
| Squamulosone | $C_{15}H_{22}O$ | FSTP | 67.60 | 5.97 | 570 | 64 |
| Verrucarol | $C_{15}H_{22}O_4$ | FSTP | 67.80 | 0.40 | 680 | 79 |
| 12-Hydroxy-6-epi-albrassitriol | $C_{15}H_{26}O_2$ | FSTP | 67.90 | 6.09 | 596 | 52 |
| Curcumenol | $C_{15}H_{22}O_2$ | FSTP | 67.90 | 0.44 | 638 | 28 |
| Shizukanolide | $C_{15}H_{18}O_2$ | FSTP | 68.20 | 0.18 | 600 | 30 |
| Longiverbenone | $C_{15}H_{22}O$ | FSTP | 68.30 | 6.21 | 598 | 59 |
| Asperilin | $C_{15}H_{20}O_3$ | FSTP | 68.30 | 5.75 | 619 | 65 |
| Dihydrocolumellarin | $C_{15}H_{22}O_2$ | FSTP | 68.50 | 5.93 | 912 | 30 |
| Solavetivone | $C_{15}H_{22}O$ | FSTP | 68.60 | 6.15 | 593 | 71 |
| Occidentalol | $C_{15}H_{24}O$ | FSTP | 68.60 | 3.87 | 727 | 39 |
| β-Cyclodihydrocostunolide | $C_{15}H_{22}O_2$ | FSTP | 68.70 | 0.62 | 565 | 66 |
| Arctiol | $C_{15}H_{26}O_2$ | FSTP | 68.80 | 3.75 | 585 | 58 |
| γ-Elemene | $C_{15}H_{24}$ | STP | 69.10 | 5.83 | 604 | 65 |
| (2R,3R,4aR,5S,8aS)-2-Hydroxy-4a,5-dimethyl-3-(prop-1-en-2-yl)-2,3,4,4a,5,6-hexahydronaphthalen-1(8aH)-one | $C_{15}H_{22}O_2$ | FSTP | 69.10 | 0.36 | 605 | 35 |
| α-Humulen-14-oic acid | $C_{15}H_{22}O_2$ | FSTP | 69.30 | 0.79 | 603 | 54 |
| Nordeoxynivalenol | $C_{15}H_{22}O_3$ | FSTP | 69.30 | 1.33 | 654 | 33 |
| Humulenol-II | $C_{15}H_{24}O$ | FSTP | 69.40 | 6.17 | 654 | 61 |
| Velleral | $C_{15}H_{20}O_2$ | FSTP | 69.50 | 0.46 | 585 | 40 |
| α-selinene | $C_{15}H_{24}O$ | FSTP | 69.60 | 6.19 | 628 | 19 |
| (3R,3aR,5R,6R,7aR)-3,6-Dimethyl-5-(prop-1-en-2-yl)-6-vinylhexahydrobenzofuran-2(3H)-one | $C_{15}H_{22}O_2$ | FSTP | 69.70 | 5.81 | 638 | 48 |

**Supplementary Table 2.** (Cont.)

| Compound Name | Mol. formula | Class. | RT I (min.) | RT II (sec.) | MF | P (%) |
|---|---|---|---|---|---|---|
| Calarene | $C_{15}H_{24}$ | STP | 69.70 | 4.07 | 609 | 71 |
| 4aH-Cycloprop[e]azulen-4a-ol, decahydro-1,1,4,7-tetramethyl-, [1aR-(1aα,4β,4aβ,7α,7aβ,7bα)]- | $C_{15}H_{26}O$ | FSTP | 70.10 | 1.39 | 479 | 25 |
| Isoalantolactone | $C_{15}H_{20}O_2$ | FSTP | 70.50 | 1.27 | 610 | 18 |
| Aromaticin | $C_{15}H_{20}O_2$ | FSTP | 70.60 | 6.19 | 694 | 37 |
| (E)-6-Hydroxy-2-methyl-6-(4-methylphenyl)hept-2-enoic acid | $C_{15}H_{20}O_3$ | FSTP | 70.70 | 2.36 | 619 | 81 |
| Ambrosin | $C_{15}H_{18}O_3$ | FSTP | 70.80 | 1.21 | 488 | 64 |
| (2S)-2-((1R,3aR,4R,5S,7aS)-1,7a-Dimethyloctahydro-1H-1,2,4-(epimethanetriyl)inden-5-yl)propan-1-ol | $C_{15}H_{24}O$ | FSTP | 70.80 | 3.81 | 639 | 97 |
| Reynosin | $C_{15}H_{20}O_3$ | FSTP | 70.80 | 0.69 | 493 | 81 |
| β-Cyclocostunolide | $C_{15}H_{20}O_2$ | FSTP | 70.90 | 1.43 | 497 | 94 |
| Velleral | $C_{15}H_{20}O_2$ | FSTP | 69.50 | 0.46 | 585 | 40 |
| α-selinene | $C_{15}H_{24}O$ | FSTP | 69.60 | 6.19 | 628 | 19 |
| 6-(2-Hydroxypropan-2-yl)-4,8a-dimethyl-2,3,4,6,7,8-hexahydro-1H-naphthalen-1-ol | $C_{15}H_{26}O_2$ | FSTP | 71.30 | 1.90 | 641 | 44 |
| 2H-2,4a-Ethanonaphthalen-8(5H)-one, hexahydro-2,5,5-trimethyl- | $C_{15}H_{24}O$ | FSTP | 71.40 | 0.28 | 632 | 49 |
| 2,5-Dimethylbenzophenone | $C_{15}H_{14}O$ | FSTP | 71.40 | 0.69 | 738 | 20 |
| 4a,7-Methano-4aH-naphth[1,8a-b]oxirene, octahydro-4,4,8,8-tetramethyl- | $C_{15}H_{24}O$ | FSTP | 71.40 | 0.36 | 632 | 61 |
| 2,4-Dimethylbenzophenone | $C_{15}H_{14}O$ | FSTP | 71.40 | 0.69 | 654 | 22 |
| (1S,4aR,7R)-1,4a-Dimethyl-7-(prop-1-en-2-yl)-1,2,3,4,4a,5,6,7-octahydronaphthalene | $C_{15}H_{24}$ | STP | 71.50 | 1.31 | 567 | 86 |
| Bohlmann k2631 | $C_{15}H_{20}O_2$ | FSTP | 71.50 | 1.35 | 668 | 18 |
| Nootkaton epoxide | $C_{15}H_{22}O_2$ | FSTP | 71.70 | 1.09 | 682 | 21 |
| 11,13-Dihydrolactucin | $C_{15}H_{18}O_3$ | FSTP | 71.80 | 5.73 | 869 | 80 |
| Isopetasol | $C_{15}H_{22}O_2$ | FSTP | 72.20 | 1.07 | 675 | 47 |
| α-Eudesmol | $C_{15}H_{22}O_2$ | FSTP | 72.40 | 1.03 | 675 | 48 |
| Limonenol | $C_{15}H_{24}O_2$ | FSTP | 72.70 | 1.96 | 535 | 83 |
| Tourneforin | $C_{15}H_{18}O_3$ | FSTP | 72.80 | 2.34 | 782 | 42 |
| 1-Penten-3-one, 4-methyl-1-[2,6,6-trimethyl-2-cyclohexen-1-yl]- | $C_{15}H_{24}O$ | FSTP | 72.90 | 3.61 | 720 | 28 |
| Spirafolide | $C_{15}H_{18}O_3$ | FSTP | 72.90 | 2.46 | 701 | 78 |
| 7-Methyl-3-methylidene-6-(3-oxobutyl)-4,7,8,8a-tetrahydro-3aH-cyclohepta[b]furan-2-one | $C_{15}H_{20}O_3$ | FSTP | 73.30 | 5.97 | 755 | 91 |
| Hydroxymuurolene | $C_{15}H_{24}O$ | FSTP | 73.60 | 6.33 | 649 | 81 |

**Supplementary Table 2.** (Cont.)

| Compound Name | Mol. formula | Class. | RT I (min.) | RT II (sec.) | MF | P (%) |
|---|---|---|---|---|---|---|
| Cedrandiol | $C_{15}H_{26}O_2$ | FSTP | 73.90 | 1.41 | 545 | 21 |
| α-cedrene | $C_{15}H_{24}$ | STP | 74.00 | 2.04 | 698 | 82 |
| 4,7-Methanofuro[3,2-c]oxacycloundecin-6(4H)-one, 7,8,9,12-tetrahydro-3,11-dimethyl- | $C_{15}H_{18}O_3$ | FSTP | 74.00 | 3.09 | 717 | 72 |
| Germacrene D | $C_{15}H_{24}$ | STP | 74.20 | 1.23 | 774 | 76 |
| Phomenone | $C_{15}H_{20}O_4$ | FSTP | 74.20 | 0.89 | 525 | 94 |
| Shyobunol | $C_{15}H_{26}O$ | FSTP | 75.40 | 0.54 | 710 | 93 |
| 1,8-Cyclopentadecadiyne | $C_{15}H_{22}$ | STP | 76.00 | 0.46 | 750 | 21 |
| 3-Oxo-10(14)-epoxyguai-11(13)-en-6,12-olide | $C_{15}H_{18}O_3$ | FSTP | 76.20 | 3.17 | 742 | 54 |
| 2,7-Cyclodecadiene-1-methanol, α,α,4,8-tetramethyl- | $C_{15}H_{26}O$ | FSTP | 76.60 | 1.43 | 546 | 91 |
| Aureonitol | $C_{15}H_{20}O_3$ | FSTP | 76.70 | 4.60 | 551 | 92 |
| 7-Methoxy-3,4-dihydro-1(2H)-phenanthrenone | $C_{15}H_{14}O_2$ | FSTP | 77.20 | 2.12 | 488 | 48 |
| Santamarine | $C_{15}H_{20}O_3$ | FSTP | 77.30 | 3.13 | 750 | 92 |
| Spirafolide | $C_{15}H_{18}O_3$ | FSTP | 72.90 | 2.46 | 701 | 78 |
| 7-Methyl-3-methylidene-6-(3-oxobutyl)-4,7,8,8a-tetrahydro-3aH-cyclohepta[b]furan-2-one | $C_{15}H_{20}O_3$ | FSTP | 73.30 | 5.97 | 755 | 91 |
| Hydroxymuurolene | $C_{15}H_{24}O$ | FSTP | 73.60 | 6.33 | 649 | 81 |
| Cedrandiol | $C_{15}H_{26}O_2$ | FSTP | 73.90 | 1.41 | 545 | 21 |
| α-cedrene | $C_{15}H_{24}$ | STP | 74.00 | 2.04 | 698 | 82 |
| 4,7-Methanofuro[3,2-c]oxacycloundecin-6(4H)-one, 7,8,9,12-tetrahydro-3,11-dimethyl- | $C_{15}H_{18}O_3$ | FSTP | 74.00 | 3.09 | 717 | 72 |
| Graveolide | $C_{15}H_{20}O_3$ | FSTP | 77.30 | 2.12 | 549 | 81 |
| 1-n-Pentyladamantane | $C_{15}H_{26}$ | STP | 78.70 | 2.86 | 741 | 94 |
| Ivalin | $C_{15}H_{20}O_4$ | FSTP | 79.20 | 1.61 | 552 | 51 |
| Costunolide | $C_{15}H_{20}O_2$ | FSTP | 79.40 | 1.77 | 752 | 45 |
| Achalensolide | $C_{15}H_{18}O_3$ | FSTP | 80.40 | 2.74 | 580 | 72 |
| T-2 Tetraol | $C_{15}H_{22}O_6$ | FSTP | 80.70 | 1.31 | 836 | 32 |
| Gazaniolide | $C_{15}H_{18}O_2$ | FSTP | 80.80 | 5.65 | 565 | 36 |
| Cyperanic acid | $C_{15}H_{22}O_4$ | FSTP | 80.90 | 3.39 | 629 | 95 |
| Pallensin | $C_{15}H_{20}O_4$ | FSTP | 82.10 | 2.52 | 544 | 40 |
| Caryophyllene oxide | $C_{15}H_{24}O$ | FSTP | 82.40 | 3.65 | 717 | 36 |

**Supplementary Table 2.** (Cont.)

| Compound Name | Mol. formula | Class. | RT I (min.) | RT II (sec.) | MF | P (%) |
|---|---|---|---|---|---|---|
| δ-Elemene | $C_{15}H_{24}$ | STP | 82.80 | 4.01 | 561 | 96 |
| 6-Hydroxy-5a,9-dimethyl-3-methylidene-4,5,6,7,9a,9b-hexahydro-3aH-benzo[g][1]benzofuran-2-one | $C_{15}H_{20}O_3$ | FSTP | 82.90 | 3.17 | 887 | 25 |
| Achillin | $C_{15}H_{18}O_3$ | FSTP | 86.90 | 3.89 | 929 | 51 |
| Achillicin | $C_{15}H_{18}O_3$ | FSTP | 86.90 | 4.11 | 729 | 32 |
| 5-[2-(Benzyloxy)ethyl]cyclohex-2-en-1-one | $C_{15}H_{18}O_2$ | FSTP | 87.40 | 4.03 | 831 | 95 |
| Pentadecanal | $C_{15}H_{30}O$ | FSTP | 87.60 | 3.17 | 772 | 97 |
| 3,7,7-Trimethyl-8-(2-methyl-propenyl)-bicyclo[4.2.0]oct-2-ene | $C_{15}H_{24}$ | STP | 88.90 | 5.12 | 615 | 66 |
| sesquithujene | $C_{15}H_{24}$ | STP | 91.80 | 3.67 | 541 | 71 |
| 2-Ethyl-5-methyl-2-phenyl-4-hexen-1-ol | $C_{15}H_{22}O$ | FSTP | 92.40 | 1.71 | 538 | 91 |
| 1-(2-Methylenecyclohexyl)-1-phenylethanol | $C_{15}H_{20}O$ | FSTP | 95.00 | 3.39 | 587 | 78 |
| 1-Cyclohexylnonene | $C_{15}H_{28}$ | STP | 95.10 | 3.27 | 580 | 40 |
| 1,4-Dimethyl 2-[(3,4-dimethoxyphenyl)methyl]butanedioate | $C_{15}H_{20}O_6$ | FSTP | 100.80 | 5.93 | 578 | 27 |
| 3-Pentadecanone | $C_{15}H_{30}O$ | FSTP | 101.10 | 3.37 | 572 | 97 |
| 1-Methylene-2b-hydroxymethyl-3,3-dimethyl-4b-(3-methylbut-2-enyl)-cyclohexane | $C_{15}H_{26}O$ | FSTP | 102.50 | 4.03 | 784 | 81 |
| 2,6,10-Dodecatrien-1-ol, 3,7,11-trimethyl-, (Z,E)- | $C_{15}H_{26}O$ | FSTP | 102.60 | 3.99 | 769 | 52 |
| 2,6,10-Dodecatrien-1-ol, 3,7,11-trimethyl- | $C_{15}H_{26}O$ | FSTP | 102.60 | 3.85 | 608 | 52 |
| Ylangenol | $C_{15}H_{24}O$ | FSTP | 103.10 | 1.15 | 541 | 96 |
| 2-Pentadecanone | $C_{15}H_{30}O$ | FSTP | 104.60 | 3.49 | 441 | 26 |
| 1-Formyl-2,2,6-trimethyl-3-cis-(3-methylbut-2-enyl)-5-cyclohexene | $C_{15}H_{24}O$ | FSTP | 107.30 | 4.74 | 582 | 60 |
| 1,7-Dimethyl-4-(1-methylethyl)cyclodecane | $C_{15}H_{30}$ | STP | 108.90 | 3.11 | 530 | 43 |
| 7-Oxabicyclo[4.1.0]heptane, 1-(1,3-dimethyl-1,3-butadienyl)-2,2,6-trimethyl-, (E)- | $C_{15}H_{24}O$ | FSTP | 110.50 | 0.93 | 574 | 97 |
| 1H-Indene, 2,3,3a,4,7,7a-hexahydro-2,2,4,4,7,7-hexamethyl- | $C_{15}H_{26}$ | STP | 113.40 | 2.02 | 597 | 32 |
| 5,7-Decadien-3-yne, 2,9-dihydroxy-5-(1-hydroxy-1-methylethyl)-2,9-dimethyl-, (Z,E)- | $C_{15}H_{24}O_3$ | FSTP | 114.60 | 1.51 | 545 | 67 |
| Guaiadiene | $C_{15}H_{24}$ | STP | 118.10 | 1.17 | 607 | 96 |
| Anisole | $C_{15}H_{24}O$ | FSTP | 120.90 | 4.64 | 667 | 72 |

**Supplementary Table 3.** Sesquiterpenoids identified across agarwood distillates (Oudh) samples using GCxGC-TOF/MS.

| Compound Name | Mol. formula | Clas. | RT I (min) | RT II (sec) | MF | P (%) |
|---|---|---|---|---|---|---|
| Aureonitol | $C_{15}H_{20}O_3$ | FSTP | 33.50 | 3.51 | 787 | 60 |
| 1,2-Dihydrothujopsene-(I1) | $C_{15}H_{26}$ | STP | 34.30 | 4.03 | 889 | 63 |
| (8R,8aS)-8,8a-Dimethyl-2-(propan-2-ylidene)-1,2,3,7,8,8a-hexahydronaphthalene | $C_{15}H_{22}$ | STP | 36.60 | 2.78 | 832 | 95 |
| Germacrene D | $C_{15}H_{24}$ | STP | 36.60 | 2.44 | 432 | 92 |
| Copaene | $C_{15}H_{24}$ | STP | 36.70 | 2.46 | 799 | 56 |
| α-Bergamotene | $C_{15}H_{24}$ | STP | 37.30 | 2.54 | 933 | 81 |
| 4b,5,6,7,8,8a,9,10-Octahydro-1-methylphenanthrene | $C_{15}H_{20}$ | FSTP | 37.90 | 3.25 | 471 | 79 |
| (R)-1-Methyl-4-(6-methylhept-5-en-2-yl)cyclohexa-1,4-diene | $C_{15}H_{24}$ | STP | 38.00 | 3.09 | 662 | 71 |
| isoledene | $C_{15}H_{24}$ | STP | 38.70 | 2.56 | 916 | 65 |
| 1H-Cycloprop[e]azulene, 1a,2,3,4,4a,5,6,7b-octahydro-1,1,4,7-tetramethyl-, [1aR-(1aα,4α,4aβ,7bα)]- | $C_{15}H_{24}$ | STP | 38.70 | 2.58 | 944 | 69 |
| 1H-Cyclopropa[a]naphthalene, decahydro-1,1,3a-trimethyl-7-methylene-, [1aS-(1aα,3aα,7aβ,7bα)]- | $C_{15}H_{24}$ | STP | 38.80 | 3.13 | 695 | 54 |
| Cyperene | $C_{15}H_{24}$ | STP | 38.80 | 2.68 | 738 | 68 |
| Bicyclo[5.3.0]decane, 2-methylene-5-(1-methylvinyl)-8-methyl- | $C_{15}H_{24}$ | STP | 39.00 | 2.64 | 712 | 64 |
| α-Bergamotene | $C_{15}H_{24}$ | STP | 39.00 | 2.76 | 736 | 41 |
| Cubenene | $C_{15}H_{24}$ | STP | 39.20 | 2.74 | 889 | 36 |
| 1H-3a,7-Methanoazulene, 2,3,4,7,8,8a-hexahydro-3,6,8,8-tetramethyl-, [3R-(3α,3aβ,7β,8aα)]- | $C_{15}H_{24}$ | STP | 39.30 | 2.78 | 588 | 81 |
| Caryophyllene | $C_{15}H_{24}$ | STP | 39.40 | 2.72 | 947 | 83 |
| 6-Methyl-2-(4-methylcyclohex-3-en-1-yl)hepta-1,5-dien-4-ol | $C_{15}H_{24}O$ | FSTP | 39.60 | 2.94 | 655 | 24 |
| 3-methyl-5-(2,6-dimethylheptyl)-1,5-Pent-2-enolide | $C_{15}H_{26}O_2$ | FSTP | 39.60 | 3.97 | 734 | 19 |
| Bicyclo[4.1.0]-3-heptene, 2-isopropenyl-5-isopropyl-7,7-dimethyl- | $C_{15}H_{24}$ | STP | 39.60 | 2.82 | 915 | 56 |
| 1,3,7,11-Cyclotetradecatetraene, 2-methyl- | $C_{15}H_{22}$ | STP | 39.80 | 3.13 | 733 | 62 |
| γ-Elemene | $C_{15}H_{24}$ | STP | 40.10 | 3.55 | 790 | 48 |
| β-Calarene | $C_{15}H_{24}$ | STP | 40.20 | 2.80 | 769 | 62 |
| 4a,5-Dimethyl-3-(prop-1-en-2-yl)-1,2,3,4,4a,5,6,7-octahydronaphthalen-1-ol | $C_{15}H_{24}O$ | FSTP | 40.30 | 3.15 | 640 | 28 |
| α-Guaiene | $C_{15}H_{24}$ | STP | 40.60 | 2.62 | 604 | 41 |
| 2-Pentadecen-4-yne, (Z)- | $C_{15}H_{26}$ | STP | 40.60 | 4.23 | 862 | 98 |
| 6,10-Dodecadien-1-yn-3-ol, 3,7,11-trimethyl- | $C_{15}H_{24}O$ | FSTP | 40.80 | 3.15 | 841 | 41 |
| Caryophyllenyl alcohol | $C_{15}H_{26}O$ | FSTP | 41.00 | 3.67 | 754 | 72 |

**Supplementary Table 3.** (Cont.)

| Compound Name | Mol. formula | Clas. | RT I (min) | RT II (sec) | MF | P (%) |
|---|---|---|---|---|---|---|
| Cycloisolongifolene | $C_{15}H_{24}$ | STP | 41.30 | 3.04 | 846 | 41 |
| Humulene | $C_{15}H_{24}$ | STP | 41.40 | 2.88 | 588 | 44 |
| 6,10-Dodecadien-3-ol, 3,7,11-trimethyl- | $C_{15}H_{28}$ | FSTP | 41.50 | 2.32 | 666 | 29 |
| 1,5-Cyclodecadiene, 1,5-dimethyl-8-(1-methylethenyl)-, [S-(Z,E)]- | $C_{15}H_{24}$ | STP | 41.50 | 3.08 | 719 | 49 |
| 1R,3Z,9S-4,11,11-Trimethyl-8-methylenebicyclo[7.2.0]undec-3-ene | $C_{15}H_{24}$ | STP | 41.50 | 2.84 | 630 | 50 |
| Prezizaene | $C_{15}H_{24}$ | STP | 41.60 | 3.11 | 787 | 24 |
| 1H-3a,7-Methanoazulene, octahydro-1,4,9,9-tetramethyl-, (1α,3aα,4β,7α,8aβ)- | $C_{15}H_{26}$ | STP | 41.80 | 2.70 | 248 | 95 |
| Rotundene | $C_{15}H_{24}$ | STP | 41.90 | 3.04 | 841 | 43 |
| Alloaromadendrene | $C_{15}H_{24}$ | STP | 41.90 | 2.90 | 764 | 31 |
| Aromandendrene | $C_{15}H_{24}$ | STP | 41.90 | 2.88 | 679 | 49 |
| β-Caryophyllene | $C_{15}H_{24}$ | STP | 42.00 | 2.98 | 828 | 33 |
| Sesquithujene | $C_{15}H_{24}$ | STP | 42.10 | 2.94 | 637 | 67 |
| β-Acoradiene | $C_{15}H_{24}$ | STP | 42.30 | 2.94 | 706 | 33 |
| (4R,4aS,6S)-4,4a-Dimethyl-6-(prop-1-en-2-yl)-1,2,3,4,4a,5,6,7-octahydronaphthalene | $C_{15}H_{24}$ | STP | 42.40 | 2.90 | 828 | 52 |
| (R)-3-Methylene-6-((S)-1,2,2-trimethylcyclopentyl)cyclohex-1-ene | $C_{15}H_{24}$ | STP | 42.50 | 3.93 | 749 | 49 |
| ɣ-Gurjunene | $C_{15}H_{24}$ | STP | 42.50 | 2.86 | 542 | 10 |
| 4a,8-Dimethyl-2-(prop-1-en-2-yl)-1,2,3,4,4a,5,6,7-octahydronaphthalene | $C_{15}H_{24}$ | STP | 42.60 | 2.84 | 545 | 42 |
| Cedrene oxide | $C_{15}H_{24}O$ | FSTP | 42.70 | 3.57 | 661 | 33 |
| Spathulenol | $C_{15}H_{24}O$ | FSTP | 42.70 | 3.55 | 722 | 14 |
| β-Acoradienol | $C_{15}H_{24}O$ | FSTP | 42.70 | 3.19 | 616 | 27 |
| α-Eremophilane | $C_{15}H_{28}$ | STP | 42.80 | 2.62 | 762 | 60 |
| 1,2,4-Metheno-1H-indene, octahydro-1,7a-dimethyl-5-(1-methylethyl)-, [1S-(1α,2α,3aβ,4α,5α,7aβ,8S*)]- | $C_{15}H_{24}$ | STP | 42.80 | 2.96 | 558 | 11 |
| Cedrenol | $C_{15}H_{24}O$ | FSTP | 42.80 | 3.00 | 837 | 84 |
| α-Amorphene | $C_{15}H_{24}$ | STP | 42.80 | 3.02 | 679 | 98 |
| Zizanene | $C_{15}H_{24}$ | STP | 42.90 | 3.00 | 808 | 35 |
| Muuroladiene | $C_{15}H_{24}$ | STP | 43.10 | 3.93 | 714 | 48 |
| 1,4-Dimethyl-7-(prop-1-en-2-yl)decahydroazulen-4-ol | $C_{15}H_{26}O$ | FSTP | 43.20 | 3.09 | 734 | 48 |
| Aristolochene | $C_{15}H_{24}$ | STP | 43.20 | 3.02 | 833 | 65 |

**Supplementary Table 3.** (Cont.)

| Compound Name | Mol. formula | Clas. | RT I (min) | RT II (sec) | MF | P (%) |
|---|---|---|---|---|---|---|
| Selinadiene | $C_{15}H_{24}$ | STP | 43.20 | 3.06 | 805 | 66 |
| β-Selinene | $C_{15}H_{24}$ | STP | 43.40 | 3.06 | 785 | 41 |
| Epoxycalamenene | $C_{15}H_{20}O$ | FSTP | 43.50 | 3.81 | 581 | 41 |
| β-Vetispirene | $C_{15}H_{22}$ | STP | 43.60 | 3.21 | 659 | 91 |
| β-Longipinene | $C_{15}H_{24}$ | STP | 43.70 | 3.08 | 731 | 42 |
| δ-Guaiene | $C_{15}H_{24}$ | STP | 44.00 | 3.04 | 589 | 31 |
| Valencene | $C_{15}H_{24}$ | STP | 44.00 | 2.92 | 811 | 21 |
| α-Farnesene | $C_{15}H_{24}$ | STP | 44.00 | 3.65 | 690 | 81 |
| Muurolane | $C_{15}H_{24}$ | STP | 44.00 | 2.96 | 511 | 62 |
| Guaiadiene | $C_{15}H_{24}$ | STP | 44.40 | 3.25 | 759 | 53 |
| Cuparene | $C_{15}H_{22}$ | STP | 44.50 | 3.75 | 649 | 91 |
| α-Curcumene | $C_{15}H_{22}$ | STP | 44.60 | 3.69 | 462 | 59 |
| 1,1,4,7-Tetramethyldecahydro-1H-cyclopropa[e]azulene-4,7-diol | $C_{15}H_{26}O_2$ | FSTP | 44.70 | 3.31 | 752 | 69 |
| Humulenol-II | $C_{15}H_{24}O$ | FSTP | 44.90 | 3.29 | 928 | 49 |
| Nootkatene | $C_{15}H_{22}$ | STP | 45.10 | 3.23 | 769 | 21 |
| γ-Muurolene | $C_{15}H_{24}$ | STP | 45.20 | 3.15 | 778 | 54 |
| 4aH-Cycloprop[e]azulen-4a-ol, decahydro-1,1,4,7-tetramethyl-, [1aR-(1aα,4β,4aβ,7α,7aβ,7bα)]- | $C_{15}H_{26}O$ | FSTP | 45.20 | 2.96 | 687 | 44 |
| Carotol | $C_{15}H_{26}O$ | FSTP | 45.20 | 3.11 | 563 | 41 |
| Caparratriene | $C_{15}H_{26}$ | STP | 45.30 | 4.38 | 770 | 61 |
| α-Panasinsen | $C_{15}H_{24}$ | STP | 45.30 | 3.11 | 791 | 24 |
| β-Cadinene | $C_{15}H_{24}$ | STP | 45.40 | 3.15 | 676 | 22 |
| 1-Isopropyl-4,7-dimethyl-1,2,3,5,6,8a-hexahydronaphthalene | $C_{15}H_{24}$ | STP | 45.40 | 3.06 | 607 | 95 |
| Calamenene | $C_{15}H_{22}$ | STP | 45.50 | 3.47 | 634 | 12 |
| 1-Isopropyl-4,7-dimethyl-1,2,3,5,6,8a-hexahydronaphthalene | $C_{15}H_{24}$ | STP | 45.60 | 3.13 | 738 | 19 |
| α-Muurolene | $C_{15}H_{24}O$ | FSTP | 45.60 | 3.63 | 745 | 52 |
| ɣ-Vetivenene | $C_{15}H_{22}$ | STP | 45.70 | 3.37 | 530 | 10 |
| Kessane | $C_{15}H_{26}O$ | FSTP | 45.80 | 3.29 | 652 | 12 |
| Cyperene epoxide | $C_{15}H_{24}O$ | FSTP | 46.10 | 3.75 | 614 | 15 |

**Supplementary Table 3.** (Cont.)

| Compound Name | Mol. formula | Clas. | RT I (min) | RT II (sec) | MF | P (%) |
|---|---|---|---|---|---|---|
| α-Cadinene | $C_{15}H_{24}$ | STP | 46.20 | 3.06 | 705 | 17 |
| (4aR,8aS)-4a-Methyl-1-methylene-7-(propan-2-ylidene)decahydronaphthalene | $C_{15}H_{24}$ | STP | 46.30 | 3.21 | 621 | 26 |
| Himachalene | $C_{15}H_{22}$ | STP | 46.30 | 3.65 | 586 | 18 |
| 4-Isopropyl-6-methyl-1-methylene-1,2,3,4-tetrahydronaphthalene | $C_{15}H_{20}$ | FSTP | 46.60 | 3.75 | 611 | 31 |
| α-Calacorene | $C_{15}H_{20}$ | FSTP | 46.60 | 3.89 | 780 | 62 |
| Cyclopentadecadiyne | $C_{15}H_{22}$ | STP | 46.80 | 5.65 | 754 | 36 |
| Selina-3,7(11)-diene | $C_{15}H_{24}$ | STP | 46.80 | 3.19 | 743 | 35 |
| Longiverbenone | $C_{15}H_{22}O$ | FSTP | 46.90 | 3.85 | 725 | 15 |
| 2-(4a,8-Dimethyl-1,2,3,4,4a,5,6,7-octahydro-naphthalen-2-yl)-prop-2-en-1-ol | $C_{15}H_{24}O$ | FSTP | 47.00 | 3.87 | 795 | 53 |
| α-Agarofuran | $C_{15}H_{24}O$ | FSTP | 47.00 | 3.93 | 591 | 27 |
| α-Calacorene | $C_{15}H_{20}$ | FSTP | 47.00 | 4.07 | 720 | 18 |
| (3aS,8aS)-6,8a-Dimethyl-3-(propan-2-ylidene)-1,2,3,3a,4,5,8,8a-octahydroazulene | $C_{15}H_{24}$ | STP | 47.10 | 4.29 | 601 | 20 |
| Coronene | $C_{15}H_{20}$ | FSTP | 47.30 | 3.75 | 793 | 77 |
| Nerolidol | $C_{15}H_{26}O$ | FSTP | 47.40 | 3.08 | 724 | 43 |
| 1,6,10-Dodecatrien-3-ol, 3,7,11-trimethyl-, (E)- | $C_{15}H_{26}O$ | FSTP | 47.40 | 3.00 | 666 | 14 |
| Hydroxycaryophyllene | $C_{15}H_{24}O$ | FSTP | 47.70 | 3.59 | 702 | 13 |
| α-Vetivone | $C_{15}H_{22}O$ | FSTP | 47.70 | 3.63 | 779 | 37 |
| Bicyclo[4.4.0]dec-2-ene-4-ol, 2-methyl-9-(prop-1-en-3-ol-2-yl)- | $C_{15}H_{24}O_2$ | FSTP | 47.80 | 3.65 | 729 | 18 |
| Bicyclo[4.4.0]dec-5-ene, 1,5-dimethyl-3-hydroxy-8-(1-methylene-2-hydroxyethyl-1)- | $C_{15}H_{24}O_2$ | FSTP | 47.80 | 3.79 | 817 | 69 |
| 4(15)-Selinene-11,12-diol | $C_{15}H_{26}O_2$ | FSTP | 48.00 | 4.76 | 684 | 18 |
| 1H-Cycloprop[e]azulene, decahydro-1,1,4,7-tetramethyl-, [1aR-(1aα,4β,4aβ,7β,7aβ,7bα)]- | $C_{15}H_{26}$ | STP | 48.10 | 4.40 | 702 | 14 |
| 2H-3,9a-Methano-1-benzoxepin, octahydro-2,2,5a,9-tetramethyl-, [3R-(3α,5aα,9α,9aα)]- | $C_{15}H_{26}O$ | FSTP | 48.30 | 4.80 | 652 | 12 |
| 4aH-cycloprop[e]azulen-4a-ol, decahydro-1,1,4,7-tetramethyl- | $C_{15}H_{26}O$ | FSTP | 48.30 | 3.69 | 722 | 28 |
| (S,1Z,6Z)-8-Isopropyl-1-methyl-5-methylenecyclodeca-1,6-diene | $C_{15}H_{24}$ | STP | 48.40 | 4.31 | 758 | 66 |
| 2-(4a,8-Dimethyl-2,3,4,4a,5,6-hexahydro-naphthalen-2-yl)-prop-2-en-1-ol | $C_{15}H_{22}O$ | FSTP | 48.40 | 3.89 | 748 | 16 |
| Isolongifolene | $C_{15}H_{20}$ | FSTP | 48.40 | 3.87 | 729 | 26 |
| 1H-3a,7-Methanoazulene, octahydro-1,4,9,9-tetramethyl- | $C_{15}H_{26}$ | STP | 48.60 | 4.48 | 750 | 26 |
| 2-((2R,4aR)-4a,8-Dimethyl-1,2,3,4,4a,5,6,7-octahydronaphthalen-2-yl)prop-2-en-1-ol | $C_{15}H_{24}O$ | FSTP | 48.90 | 3.93 | 759 | 14 |

**Supplementary Table 3.** (Cont.)

| Compound Name | Mol. formula | Clas. | RT I (min) | RT II (sec) | MF | P (%) |
|---|---|---|---|---|---|---|
| 1,4-Dihydrothujopsene-(I1) | $C_{15}H_{26}$ | STP | 49.00 | 0.46 | 861 | 28 |
| Humulene epoxide I | $C_{15}H_{24}O$ | FSTP | 49.00 | 0.34 | 799 | 28 |
| Humulene oxide II | $C_{15}H_{24}O$ | FSTP | 49.40 | 0.50 | 682 | 36 |
| 5-Azulenemethanol, 1,2,3,4,5,6,7,8-octahydro-α,α,3,8-tetramethyl- | $C_{15}H_{26}O$ | FSTP | 49.70 | 3.57 | 824 | 16 |
| 2-((2R,4aR,8aR)-4a,8-Dimethyl-1,2,3,4,4a,5,6,8a-octahydronaphthalen-2-yl)prop-2-en-1-ol | $C_{15}H_{24}O$ | FSTP | 49.70 | 3.93 | 785 | 31 |
| 1,4-Methanoazulen-9-ol, decahydro-1,5,5,8a-tetramethyl-, [1R-(1α .. | $C_{15}H_{26}O$ | FSTP | 49.90 | 3.91 | 552 | 34 |
| Isoaromadendrene epoxide | $C_{15}H_{24}O$ | FSTP | 50.00 | 3.79 | 794 | 24 |
| (R)-3,5,8a-Trimethyl-7,8,8a,9-tetrahydronaphtho[2,3-b]furan-4(6H)-one | $C_{15}H_{18}O_2$ | FSTP | 50.10 | 4.72 | 720 | 13 |
| α-Isocomene | $C_{15}H_{24}$ | STP | 50.10 | 4.42 | 769 | 17 |
| α-cyperone | $C_{15}H_{20}O$ | FSTP | 50.60 | 4.60 | 873 | 43 |
| 4a(2H)-Naphthalenol, 1,3,4,5,6,8a-hexahydro-4,7-dimethyl-1-(1-methylethyl)-, (1S,4R,4aS,8aR)- | $C_{15}H_{26}O$ | FSTP | 50.70 | 3.63 | 783 | 35 |
| 2H-2,4a-Ethanonaphthalen-8(5H)-one, hexahydro-2,5,5-trimethyl- | $C_{15}H_{24}O$ | FSTP | 50.70 | 4.27 | 517 | 21 |
| Neointermedeol | $C_{15}H_{26}O$ | FSTP | 50.70 | 3.75 | 612 | 65 |
| 1H-Benzocyclohepten-7-ol, 2,3,4,4a,5,6,7,8-octahydro-1,1,4a,7-tetramethyl-, cis- | $C_{15}H_{26}O$ | FSTP | 50.80 | 3.53 | 562 | 18 |
| Cyperen-8-one | $C_{15}H_{22}O$ | FSTP | 50.90 | 4.34 | 803 | 12 |
| Maaliol | $C_{15}H_{26}O$ | FSTP | 50.90 | 3.73 | 618 | 27 |
| Junenol | $C_{15}H_{26}O$ | FSTP | 51.10 | 3.85 | 872 | 58 |
| β-Guaiene | $C_{15}H_{24}$ | STP | 51.10 | 4.15 | 657 | 69 |
| 1,3a-Ethano-3aH-indene, 1,2,3,6,7,7a-hexahydro-2,2,4,7a-tetramethyl-, [1R-(1α,3aα,7aα)]- | $C_{15}H_{24}$ | STP | 51.30 | 5.46 | 615 | 48 |
| 2H-Cyclopropa[a]naphthalen-2-one, 1,1a,4,5,6,7,7a,7b-octahydro-1,1,7,7a-tetramethyl-, (1aα,7α,7aα,7bα)- | $C_{15}H_{22}O$ | FSTP | 51.30 | 4.68 | 757 | 16 |
| β-Acorenol | $C_{15}H_{26}O$ | FSTP | 51.50 | 0.36 | 795 | 23 |
| 2-(4a,8-Dimethyl-2,3,4,5,6,7-hexahydro-1H-naphthalen-2-yl)propan-2-ol | $C_{15}H_{26}O$ | FSTP | 51.80 | 3.93 | 840 | 83 |
| 2-(4a,8-Dimethyl-2,3,4,5,6,7-hexahydro-1H-naphthalen-2-yl)propan-2-ol | $C_{15}H_{26}O$ | FSTP | 51.80 | 3.93 | 840 | 83 |
| Aristolene | $C_{15}H_{24}$ | STP | 51.80 | 5.22 | 770 | 53 |
| Curcumenol | $C_{15}H_{22}O_2$ | FSTP | 51.90 | 4.74 | 625 | 34 |
| Zonarene | $C_{15}H_{24}$ | STP | 52.00 | 4.40 | 569 | 44 |
| Agarospirol | $C_{15}H_{26}O$ | FSTP | 52.00 | 3.93 | 821 | 52 |
| 4,7-Methanoazulene, 1,2,3,4,5,6,7,8-octahydro-1,4,9,9-tetramethyl-, [1S-(1α,4α,7α)]- | $C_{15}H_{24}$ | STP | 52.00 | 4.50 | 823 | 59 |

**Supplementary Table 3.** (Cont.)

| Compound Name | Mol. formula | Clas. | RT I (min) | RT II (sec) | MF | P (%) |
|---|---|---|---|---|---|---|
| 1-Naphthalenol, 1,2,3,4,4a,7,8,8a-octahydro-1,6-dimethyl-4-(1-methylethyl)-, [1R-(1α,4β,4aβ,8aβ)]- | $C_{15}H_{26}O$ | FSTP | 52.10 | 3.77 | 831 | 38 |
| γ-Gurjunenepoxide-(2) | $C_{15}H_{24}O$ | FSTP | 52.40 | 3.95 | 764 | 38 |
| Rosifoliol | $C_{15}H_{26}O$ | FSTP | 52.50 | 3.99 | 604 | 45 |
| Ambrosin | $C_{15}H_{18}O_3$ | FSTP | 52.60 | 4.54 | 548 | 82 |
| β-Cyclocostunolide | $C_{15}H_{20}O_2$ | FSTP | 52.70 | 4.50 | 788 | 40 |
| 4a,trans-8a-Perhydro-cis-2-(2-hydroxy-2-propyl)-4a,cis-8-dimethylnaphthalene | $C_{15}H_{28}$ | FSTP | 52.80 | 3.75 | 807 | 34 |
| β-Eudesmol | $C_{15}H_{24}O$ | FSTP | 52.80 | 4.19 | 674 | 32 |
| 2-Naphthalenemethanol, decahydro-α,α,4a-trimethyl-8-methylene-, [2R-(2α,4aα,8aβ)]- | $C_{15}H_{26}O$ | FSTP | 52.90 | 4.01 | 768 | 13 |
| Guaiol | $C_{15}H_{26}O$ | FSTP | 52.90 | 4.07 | 788 | 52 |
| α-Eudesmol | $C_{15}H_{26}O$ | FSTP | 53.00 | 4.27 | 592 | 47 |
| Cubenol | $C_{15}H_{26}O$ | FSTP | 53.00 | 4.11 | 590 | 37 |
| Viridiflorol | $C_{15}H_{26}O$ | FSTP | 53.10 | 3.95 | 798 | 16 |
| Hedycaryol | $C_{15}H_{26}O$ | FSTP | 53.10 | 3.95 | 808 | 30 |
| 2-Furanmethanol, tetrahydro-α,α,5-trimethyl-5-(4-methyl-3-cyclohexen-1-yl)-, [2S-[2α,5β(R*)]]- | $C_{15}H_{26}O_2$ | FSTP | 53.20 | 3.55 | 709 | 19 |
| Globulol | $C_{15}H_{26}O$ | FSTP | 53.30 | 4.56 | 807 | 23 |
| Longipinene epoxide | $C_{15}H_{24}O$ | FSTP | 53.40 | 2.22 | 728 | 37 |
| δ-Neoclovene | $C_{15}H_{26}$ | STP | 53.40 | 4.31 | 675 | 17 |
| 4-[(E)-5-Hydroxy-3-methylpent-3-enyl]-3,5,5-trimethylcyclohex-2-en-1-one | $C_{15}H_{24}O_2$ | FSTP | 53.40 | 0.60 | 808 | 24 |
| Khusiol | $C_{15}H_{26}O$ | FSTP | 53.50 | 4.38 | 742 | 14 |
| Patchouli alcohol | $C_{15}H_{26}O$ | FSTP | 53.50 | 4.38 | 746 | 17 |
| 5-Azulenemethanol, 1,2,3,3a,4,5,6,7-octahydro-α,α,3,8-tetramethyl-, [3S-(3α,3aβ,5α)]- | $C_{15}H_{26}O$ | FSTP | 53.60 | 3.99 | 663 | 63 |
| 2-Naphthalenemethanol, 1,2,3,4,4a,5,6,8a-octahydro-α,α,4a,8-tetramethyl-, (2α,4aα,8aα)- | $C_{15}H_{26}O$ | FSTP | 53.60 | 3.87 | 670 | 74 |
| α-Bisabolene epoxide | $C_{15}H_{24}O$ | FSTP | 53.90 | 4.21 | 730 | 32 |
| (7a-Isopropenyl-4,5-dimethyloctahydroinden-4-yl)methanol | $C_{15}H_{26}O$ | FSTP | 54.00 | 3.73 | 796 | 78 |
| 3-Cyclohexen-1-ol, 1-(1,5-dimethyl-4-hexenyl)-4-methyl- | $C_{15}H_{26}O$ | FSTP | 54.00 | 3.55 | 832 | 32 |
| 1H-3a,7-Methanoazulene, octahydro-1,4,9,9-tetramethyl-, (1α,3aα,4β,7α,8aβ)- | $C_{15}H_{26}$ | STP | 54.00 | 3.71 | 597 | 16 |
| Guaiazulene | $C_{15}H_{18}$ | STP | 54.00 | 4.64 | 830 | 30 |
| 1H-Benzocycloheptene, 2,4a,5,6,7,8,9,9a-octahydro-3,5,5-trimethyl-9-methylene-, (4aS-cis)- | $C_{15}H_{24}$ | STP | 54.20 | 4.90 | 791 | 9 |

**Supplementary Table 3.** (Cont.)

| Compound Name | Mol. formula | Clas. | RT I (min) | RT II (sec) | MF | P (%) |
|---|---|---|---|---|---|---|
| Germacraol | $C_{15}H_{24}O$ | FSTP | 54.20 | 4.44 | 783 | 11 |
| 2,6,10-Dodecatrien-1-ol, 3,7,11-trimethyl- | $C_{15}H_{26}O$ | FSTP | 54.30 | 3.51 | 572 | 22 |
| Mustakone | $C_{15}H_{22}O$ | FSTP | 54.40 | 4.58 | 785 | 14 |
| 1-(1,3-Dimethyl-buta-1,3-dienyl)-3,7,7-trimethyl-2-oxa-bicyclo[3.2.0]hept-3-ene | $C_{15}H_{22}O$ | FSTP | 54.50 | 1.25 | 768 | 11 |
| α-Bisabolol | $C_{15}H_{26}O$ | FSTP | 54.70 | 3.61 | 626 | 23 |
| Epizizanone | $C_{15}H_{22}O$ | FSTP | 54.70 | 4.82 | 823 | 26 |
| Taylorione | $C_{15}H_{22}O$ | FSTP | 54.90 | 4.84 | 819 | 33 |
| Longifolene | $C_{15}H_{24}$ | STP | 55.00 | 5.00 | 816 | 30 |
| 1-Methylene-2b-hydroxymethyl-3,3-dimethyl-4b-(3-methylbut-2-enyl)-cyclohexane | $C_{15}H_{26}O$ | FSTP | 55.00 | 3.39 | 649 | 17 |
| Alloaromadendrene oxide | $C_{15}H_{24}O$ | FSTP | 55.10 | 4.36 | 526 | 13 |
| Aromadendrene oxide-(1) | $C_{15}H_{24}O$ | FSTP | 55.30 | 4.27 | 705 | 28 |
| 3,5,11-Eudesmatriene | $C_{15}H_{22}$ | STP | 55.30 | 5.20 | 809 | 16 |
| Cyperotundone | $C_{15}H_{22}O$ | FSTP | 55.50 | 5.30 | 779 | 24 |
| 7-(1,3-Dimethylbuta-1,3-dienyl)-1,6,6-trimethyl-3,8-dioxatricyclo[5.1.0.0(2,4)]octane | $C_{15}H_{22}O_2$ | FSTP | 55.60 | 5.10 | 738 | 21 |
| 1H-Cyclopropa[a]naphthalene, 1a,2,6,7,7a,7b-hexahydro-1,1,7,7a-tetramethyl-, [1aR-(1aα,7α,7aα,7bα)]- | $C_{15}H_{22}$ | STP | 55.60 | 5.46 | 779 | 21 |
| Eudesmatriene | $C_{15}H_{22}$ | STP | 55.70 | 5.63 | 789 | 18 |
| 4-isopropyl-1,6-dimethyl-1,2,3,4-tetrahydronaphthalene | $C_{15}H_{22}$ | STP | 55.70 | 5.48 | 548 | 21 |
| 2,4a,5,8a-Tetramethyl-1,2,3,4,4a,7,8,8a-octahydronaphthalen-1-yl formate (isomer 1) | $C_{15}H_{24}O_2$ | FSTP | 55.90 | 1.15 | 821 | 20 |
| Eucalyptol | $C_{15}H_{22}O$ | FSTP | 56.00 | 4.58 | 785 | 17 |
| 2-((2R,4aR,8aR)-4a,8-Dimethyl-1,2,3,4,4a,5,6,8a-octahydronaphthalen-2-yl)acrylaldehyde | $C_{15}H_{22}O$ | FSTP | 56.00 | 4.66 | 737 | 16 |
| 1H-Cyclopropa[a]naphthalene, 1a,2,3,3a,4,5,6,7b-octahydro-1,1,3a,7-tetramethyl-, [1aR-(1aα,3aα,7bα)]- | $C_{15}H_{24}$ | STP | 56.10 | 5.04 | 585 | 16 |
| Isocaryophyllene | $C_{15}H_{24}$ | STP | 56.20 | 1.25 | 725 | 32 |
| 4,6,6-Trimethyl-2-(3-methylbuta-1,3-dienyl)-3-oxatricyclo[5.1.0.0(2,4)]octane | $C_{15}H_{22}O$ | FSTP | 56.20 | 4.52 | 749 | 30 |
| 2-Pentanone, 4-cyclohexylidene-3,3-diethyl- | $C_{15}H_{26}O$ | FSTP | 56.20 | 3.65 | 772 | 23 |
| 2(1H)Naphthalenone, 3,5,6,7,8,8a-hexahydro-4,8a-dimethyl-6-(1-methylethenyl)- | $C_{15}H_{22}O$ | FSTP | 56.30 | 4.58 | 713 | 87 |
| 2-((2S,4aR)-4a,8-Dimethyl-1,2,3,4,4a,5,6,7-octahydronaphthalen-2-yl)propan-2-ol | $C_{15}H_{26}O$ | FSTP | 56.50 | 5.22 | 798 | 65 |
| (R)-2-((4aS,8aR)-4a-Methyl-8-methylene-1,4,4a,5,6,7,8,8a-octahydronaphthalen-2-yl)propan-1-ol | $C_{15}H_{24}O$ | FSTP | 56.60 | 4.40 | 839 | 77 |
| T-2 Tetraol | $C_{15}H_{22}O_6$ | FSTP | 56.70 | 4.96 | 583 | 14 |

**Supplementary Table 3.** (Cont.)

| Compound Name | Mol. formula | Clas. | RT I (min) | RT II (sec) | MF | P (%) |
|---|---|---|---|---|---|---|
| Verrucarol | $C_{15}H_{22}O_4$ | FSTP | 56.70 | 4.86 | 812 | 22 |
| Naphtho[1,2-b]furan-2-one, 2,3,3a,4,5,5a,6,7,9a,9b-decahydro-3,5a,9-trimethyl-7,9a-peroxy- | $C_{15}H_{20}O_4$ | FSTP | 56.70 | 3.67 | 672 | 12 |
| Gossonorol | $C_{15}H_{22}O$ | FSTP | 56.90 | 5.83 | 751 | 19 |
| 6-Isopropenyl-4,8a-dimethyl-1,2,3,5,6,7,8,8a-octahydro-naphthalen-2-ol | $C_{15}H_{24}O$ | FSTP | 56.90 | 4.46 | 697 | 68 |
| Sesquisabinene hydrate | $C_{15}H_{26}O$ | FSTP | 57.30 | 4.11 | 832 | 29 |
| Ledol | $C_{15}H_{26}O$ | FSTP | 57.50 | 3.93 | 748 | 25 |
| α-Santalol | $C_{15}H_{24}O$ | FSTP | 57.60 | 4.05 | 571 | 53 |
| α-Vetivol | $C_{15}H_{24}O$ | FSTP | 57.60 | 4.44 | 804 | 18 |
| Cadinene | $C_{15}H_{22}O$ | FSTP | 57.60 | 4.92 | 680 | 15 |
| Valerenol | $C_{15}H_{24}O$ | FSTP | 57.60 | 4.46 | 846 | 33 |
| 1-Cyclopropene-1-pentanol, α,ε,ε,2-tetramethyl-3-(1-methylethenyl)- | $C_{15}H_{26}O$ | FSTP | 57.70 | 5.79 | 741 | 18 |
| Khusimol | $C_{15}H_{24}O$ | FSTP | 57.80 | 4.86 | 770 | 29 |
| 1(2H)-Naphthalenone, 3,4,4a,5,6,7-hexahydro-4a,5-dimethyl-3-(1-methylethenyl)-, [3S-(3α,4aα,5α)]- | $C_{15}H_{22}O$ | FSTP | 57.90 | 5.30 | 748 | 42 |
| 7-Oxabicyclo[4.1.0]heptane, 1-(1,3-dimethyl-1,3-butadienyl)-2,2,6-trimethyl-, (E)- | $C_{15}H_{24}O$ | FSTP | 57.90 | 4.44 | 628 | 23 |
| 1-Naphthalenol, decahydro-1,4a-dimethyl-7-(1-methylethylidene)-, [1R-(1α,4aβ,8aα)]- | $C_{15}H_{26}O$ | FSTP | 58.00 | 4.52 | 797 | 16 |
| 7-Isopropenyl-1,4a-dimethyl-4,4a,5,6,7,8-hexahydro-3H-naphthalen-2-one | $C_{15}H_{22}O$ | FSTP | 58.20 | 4.82 | 702 | 16 |
| β-Oplopenone | $C_{15}H_{24}O$ | FSTP | 58.50 | 3.06 | 672 | 72 |
| Squamulosone | $C_{15}H_{22}O$ | FSTP | 58.80 | 5.28 | 610 | 36 |
| Santalcamphor | $C_{15}H_{24}O_2$ | FSTP | 59.00 | 5.02 | 779 | 20 |
| Calarene epoxide | $C_{15}H_{24}O$ | FSTP | 59.00 | 4.96 | 666 | 55 |
| Epiglobulol | $C_{15}H_{26}O$ | FSTP | 59.20 | 4.86 | 605 | 22 |
| 1-Formyl-2,2-dimethyl-3-trans-(3-methyl-but-2-enyl)-6-methylidene-cyclohexane | $C_{15}H_{24}O$ | FSTP | 59.30 | 2.68 | 742 | 21 |
| Isolongifolol | $C_{15}H_{26}O$ | FSTP | 59.40 | 4.21 | 745 | 20 |
| Silphiperfoladiene | $C_{15}H_{22}$ | STP | 59.50 | 5.26 | 681 | 24 |
| Arctiol | $C_{15}H_{26}O_2$ | FSTP | 59.70 | 3.95 | 739 | 21 |
| 6-(1-Hydroxymethylvinyl)-4,8a-dimethyl-3,5,6,7,8,8a-hexahydro-1H-naphthalen-2-one | $C_{15}H_{22}O_2$ | FSTP | 59.70 | 5.32 | 746 | 15 |
| Parthenolide | $C_{15}H_{20}O_3$ | FSTP | 59.80 | 5.22 | 769 | 19 |
| (E)-2-((8R,8aS)-8,8a-Dimethyl-3,4,6,7,8,8a-hexahydronaphthalen-2(1H)-ylidene)propan-1-ol | $C_{15}H_{24}O$ | FSTP | 60.00 | 4.94 | 687 | 30 |

**Supplementary Table 3.** (Cont.)

| Compound Name | Mol. formula | Clas. | RT I (min) | RT II (sec) | MF | P (%) |
|---|---|---|---|---|---|---|
| 2-Methyl-3-(3-methyl-but-2-enyl)-2-(4-methyl-pent-3-enyl)-oxetane | $C_{15}H_{26}O$ | FSTP | 60.00 | 2.84 | 772 | 27 |
| 5,6-Azulenedimethanol, 1,2,3,3a,8,8a-hexahydro-2,2,8-trimethyl-, (3aα,8β,8aα)- | $C_{15}H_{24}O_2$ | FSTP | 60.10 | 4.92 | 727 | 93 |
| 4,4-Dimethyl-3-(3-methylbut-3-enylidene)-2-methylenebicyclo[4.1.0]heptane | $C_{15}H_{22}$ | STP | 60.30 | 5.61 | 693 | 16 |
| α-Isonootkatol | $C_{15}H_{24}O$ | FSTP | 60.40 | 5.52 | 754 | 16 |
| Spiro[tricyclo[4.4.0.0(5,9)]decane-10,2'-oxirane], 1-methyl-4-isopropyl-7,8-dihydroxy-, (8S)- | $C_{15}H_{24}O_3$ | FSTP | 60.40 | 5.34 | 732 | 94 |
| Bicyclo[5.2.0]nonane, 4-methylene-2,8,8-trimethyl-2-vinyl- | $C_{15}H_{24}$ | STP | 60.50 | 5.81 | 660 | 23 |
| 2,4a-Methanonaphthalen-7(4aH)-one, 1,2,3,4,5,6-hexahydro-1,1,5,5-tetramethyl-, (2s-cis)- | $C_{15}H_{22}O$ | FSTP | 60.60 | 4.88 | 733 | 41 |
| Linderane | $C_{15}H_{16}O_4$ | FSTP | 60.70 | 5.63 | 790 | 27 |
| Ylangenal | $C_{15}H_{22}O$ | FSTP | 60.90 | 5.12 | 775 | 20 |
| 3-Isopropyl-6,7-dimethyltricyclo[4.4.0.0(2,8)]decane-9,10-diol | $C_{15}H_{26}O_2$ | FSTP | 61.00 | 4.21 | 759 | 39 |
| 7-Oxabicyclo[4.1.0]heptane, 2,2,6-trimethyl-1-(3-methyl-1,3-butadienyl)-5-methylene- | $C_{15}H_{22}O$ | FSTP | 61.00 | 5.32 | 700 | 54 |
| 1H-3a,7-Methanoazulene-6-methanol, 2,3,4,7,8,8a-hexahydro-3,8,8-trimethyl-, [3R-(3α,3aβ,7β,8aα)]- | $C_{15}H_{24}O$ | FSTP | 61.10 | 4.34 | 593 | 24 |
| Nootkatone | $C_{15}H_{22}O$ | FSTP | 61.20 | 5.26 | 768 | 53 |
| 1H-3a,7-Methanoazulene, 2,3,6,7,8,8a-hexahydro-1,4,9,9-tetramethyl-, (1α,3aα,7α,8aβ)- | $C_{15}H_{24}$ | STP | 61.20 | 4.46 | 854 | 96 |
| 4,4-Dimethyl-3-(3-methylbut-2-enylidene)octane-2,7-dione | $C_{15}H_{24}O_2$ | FSTP | 61.40 | 3.73 | 793 | 17 |
| (5R,10R)-6,10-Dimethyl-2-(propan-2-ylidene)spiro[4.5]dec-6-en-8-one | $C_{15}H_{22}O$ | FSTP | 61.50 | 5.16 | 783 | 21 |
| Cyperolactone | $C_{15}H_{22}O_2$ | FSTP | 61.70 | 5.52 | 595 | 30 |
| Cumanin | $C_{15}H_{22}O_4$ | FSTP | 62.10 | 5.50 | 689 | 68 |
| 1,4-Methanonaphthalene, 6,7-diethyldecahydro-, cis- | $C_{15}H_{26}$ | STP | 62.30 | 3.91 | 624 | 69 |
| 5-Hydroxymethyl-1,1,4a-trimethyl-6-methylenedecahydronaphthalen-2-ol | $C_{15}H_{26}O_2$ | FSTP | 62.40 | 4.09 | 787 | 20 |
| Uvidin C | $C_{15}H_{26}O_3$ | FSTP | 62.50 | 4.66 | 848 | 85 |
| Cyperadione | $C_{15}H_{24}O_2$ | FSTP | 62.70 | 5.75 | 773 | 19 |
| Ledene oxide-(I) | $C_{15}H_{24}O$ | FSTP | 62.70 | 3.21 | 787 | 14 |
| Nootkaton epoxide | $C_{15}H_{22}O_2$ | FSTP | 62.80 | 5.65 | 477 | 42 |
| (4aR,5S)-1-Hydroxy-4a,5-dimethyl-3-(propan-2-ylidene)-4,4a,5,6,7,8-hexahydronaphthalen-2(3H)-one | $C_{15}H_{22}O_2$ | FSTP | 63.20 | 5.22 | 722 | 13 |
| Corymbolone | $C_{15}H_{24}O_2$ | FSTP | 63.70 | 5.59 | 774 | 17 |
| Neoisolongifolene | $C_{15}H_{22}O$ | FSTP | 63.90 | 0.87 | 627 | 15 |
| Cedrol | $C_{15}H_{26}O$ | FSTP | 64.10 | 5.56 | 789 | 12 |

**Supplementary Table 3.** (Cont.)

| Compound Name | Mol. formula | Clas. | RT I (min) | RT II (sec) | MF | P (%) |
|---|---|---|---|---|---|---|
| 2(3H)-Benzofuranone, 6-ethenylhexahydro-6-methyl-3-methylene-7-(1-methylethenyl)-, [3aS-(3aα,6α,7β,7aβ)]- | $C_{15}H_{20}O_2$ | FSTP | 64.20 | 5.75 | 822 | 27 |
| 5,7-Decadien-3-yne, 2,9-dihydroxy-5-(1-hydroxy-1-methylethyl)-2,9-dimethyl-, (Z,E)- | $C_{15}H_{24}O_3$ | FSTP | 64.50 | 4.15 | 774 | 43 |
| 7-Tetracyclo[6.2.1.0(3.8)0(3.9)]undecanol, 4,4,11,11-tetramethyl- | $C_{15}H_{24}O$ | FSTP | 64.50 | 0.77 | 776 | 21 |
| 1-Acetyl-4,6,8-trimethylazulene | $C_{15}H_{16}O$ | FSTP | 64.50 | 0.46 | 584 | 50 |
| Clovanediol | $C_{15}H_{26}O_2$ | FSTP | 64.60 | 5.83 | 709 | 16 |
| Psilostachyin B | $C_{15}H_{18}O_4$ | FSTP | 64.60 | 5.81 | 791 | 16 |
| 2-Cyclohexene-1-carboxaldehyde, 2,6-dimethyl-6-(4-methyl-3-pentenyl)- | $C_{15}H_{24}O$ | FSTP | 64.60 | 4.54 | 659 | 61 |
| α-Hydroxyculmorin | $C_{15}H_{26}O_3$ | FSTP | 64.80 | 6.15 | 774 | 15 |
| Shyobunone | $C_{15}H_{24}O$ | FSTP | 64.80 | 2.96 | 810 | 21 |
| Solstitialin A | $C_{15}H_{20}O_5$ | FSTP | 64.90 | 0.22 | 664 | 23 |
| 3-Methoxymethyl-2,5,5,8a-tetramethyl-6,7,8,8a-tetrahydro-5H-chromene | $C_{15}H_{24}O_2$ | FSTP | 65.10 | 5.85 | 734 | 18 |
| Callitrin | $C_{15}H_{22}O_2$ | FSTP | 65.40 | 5.42 | 799 | 36 |
| 1,4-Methanoazulen-3-ol, decahydro-1,5,5,8a-tetramethyl-, [1S-(1α,3β,3aβ,4α,8aβ)]- | $C_{15}H_{26}O$ | FSTP | 65.50 | 6.35 | 736 | 11 |
| Valerenic acid | $C_{15}H_{22}O_2$ | FSTP | 65.70 | 5.57 | 901 | 20 |
| 8-Deoxylactucin | $C_{15}H_{16}O_4$ | FSTP | 66.30 | 1.01 | 687 | 34 |
| 1H-Indene, 2,3,3a,4,7,7a-hexahydro-2,2,4,4,7,7-hexamethyl-, trans- | $C_{15}H_{26}$ | STP | 66.30 | 2.98 | 757 | 14 |
| 5,9b-Dihydroxy-6,6,9a-trimethyl-5,5a,8,9-tetrahydro-3H-benzo[g][2]benzofuran-1,7-dione | $C_{15}H_{20}O_5$ | FSTP | 66.40 | 6.01 | 560 | 52 |
| 2(3H)-Benzofuranone, 6-ethenylhexahydro-3,6-dimethyl-7-(1-methylethenyl)-, [3S-(3α,3aα,6α,7β,7aβ)]- | $C_{15}H_{22}O_2$ | FSTP | 66.40 | 5.67 | 820 | 50 |
| Asperilin | $C_{15}H_{20}O_3$ | FSTP | 66.60 | 6.13 | 766 | 25 |
| Cubedol | $C_{15}H_{26}O$ | FSTP | 66.60 | 0.34 | 801 | 19 |
| 2(1H)-Naphthalenone, 4a,5,6,7,8,8a-hexahydro-6-[1-(hydroxymethyl)ethenyl]-4,8a-dimethyl-, [4ar-(4aα,6α,8aβ)]- | $C_{15}H_{22}O_2$ | FSTP | 66.70 | 0.32 | 632 | 15 |
| Cyclolongifolene oxide | $C_{15}H_{22}O$ | FSTP | 66.90 | 1.43 | 793 | 32 |
| Dihydrolactucin | $C_{15}H_{18}O_5$ | FSTP | 66.90 | 4.94 | 584 | 16 |
| Carissone | $C_{15}H_{24}O_2$ | FSTP | 67.10 | 5.79 | 814 | 52 |
| β-Cyclodihydrocostunolide | $C_{15}H_{22}O_2$ | FSTP | 67.40 | 6.49 | 746 | 72 |
| Costunolide | $C_{15}H_{20}O_2$ | FSTP | 67.60 | 5.91 | 739 | 36 |
| 4-(3,3-Dimethyl-but-1-ynyl)-4-hydroxy-2,6,6-trimethylcyclohex-2-enone | $C_{15}H_{22}O_2$ | FSTP | 68.10 | 6.21 | 797 | 15 |
| 2,3,3-Trimethyl-2-(3-methylbuta-1,3-dienyl)-6-methylenecyclohexanone | $C_{15}H_{22}O$ | FSTP | 68.60 | 6.67 | 737 | 20 |

**Supplementary Table 3.** (Cont.)

| Compound Name | Mol. formula | Clas. | RT I (min) | RT II (sec) | MF | P (%) |
|---|---|---|---|---|---|---|
| Dihydrocolumellarin | $C_{15}H_{22}O_2$ | FSTP | 68.60 | 6.03 | 663 | 27 |
| Cedrandiol | $C_{15}H_{26}O_2$ | FSTP | 68.70 | 6.61 | 782 | 28 |
| Zizanoic acid | $C_{15}H_{22}O_2$ | FSTP | 69.40 | 3.17 | 656 | 32 |
| 4-(Hydroxymethyl)-3,4a,8,8-tetramethyl-1,2,5,6,7,8a-hexahydronaphthalene-1,2-diol | $C_{15}H_{26}O_3$ | FSTP | 69.90 | 0.50 | 704 | 15 |
| 2-((2R,4aR,8aS)-4a-Methyl-8-methylenedecahydronaphthalen-2-yl)acrylaldehyde | $C_{15}H_{22}O$ | FSTP | 70.00 | 0.32 | 697 | 10 |
| Furanoeremophilone | $C_{15}H_{20}O_2$ | FSTP | 70.40 | 0.67 | 775 | 23 |
| Columellarin | $C_{15}H_{20}O_2$ | FSTP | 71.00 | 6.33 | 706 | 24 |
| 1-Formyl-2,2,6-trimethyl-3-cis-(3-methylbut-2-enyl)-5-cyclohexene | $C_{15}H_{24}O$ | FSTP | 71.50 | 3.69 | 823 | 61 |
| 5,6-Azulenedicarboxaldehyde, 1,2,3,3a,8,8a-hexahydro-2,2,8-trimethyl-, (3aα,8α,8aα)-(-)- | $C_{15}H_{20}O_2$ | FSTP | 71.70 | 1.09 | 698 | 37 |
| 2-Acetoxy-1,1,10-trimethyl-6,9-epidioxydecalin | $C_{15}H_{24}O_4$ | FSTP | 71.90 | 6.25 | 553 | 39 |
| Lapachol | $C_{15}H_{14}O_3$ | FSTP | 71.90 | 6.19 | 732 | 44 |
| 3α,4α,9β,11-Diepoxymuurolan-10-ol | $C_{15}H_{24}O_3$ | FSTP | 72.20 | 6.07 | 779 | 39 |
| Naphtho[2,3-b]furan-2(4H)-one, 4a,5,6,7,8,8a,9,9a-octahydro-3,8a-dimethyl-5-methylene- | $C_{15}H_{20}O_2$ | FSTP | 72.20 | 0.56 | 686 | 99 |
| Reynosin | $C_{15}H_{20}O_3$ | FSTP | 73.10 | 0.79 | 492 | 44 |
| Spirojatamol | $C_{15}H_{26}O$ | FSTP | 73.50 | 5.65 | 787 | 37 |
| Hydroxyvalerenic acid | $C_{15}H_{22}O_3$ | FSTP | 73.80 | 0.65 | 756 | 22 |
| Cyperanic acid | $C_{15}H_{22}O_4$ | FSTP | 73.90 | 5.89 | 639 | 68 |
| Nonanophenone | $C_{15}H_{22}O$ | FSTP | 74.00 | 0.89 | 749 | 40 |
| Gansongone | $C_{15}H_{22}O$ | FSTP | 74.10 | 1.33 | 728 | 29 |
| Humulaneol | $C_{15}H_{26}O$ | FSTP | 75.20 | 4.21 | 789 | 83 |
| 1H-3a,7-Methanoazulene, octahydro-3,8,8-trimethyl-6-methylene-, [3R-(3α,3aβ,7β,8aα)]- | $C_{15}H_{24}$ | STP | 75.40 | 1.63 | 714 | 25 |
| 2H-Cyclopentacyclooctene, 4,5,6,7,8,9-hexahydro-1,2,2,3-tetramethyl- | $C_{15}H_{24}$ | STP | 76.60 | 4.33 | 745 | 43 |
| Alantolactone | $C_{15}H_{20}O_3$ | FSTP | 76.60 | 5.36 | 775 | 37 |
| Naphtho[2,3-b]furan-2(3H)-one, 4a,5,6,7,8,8a-hexahydro-3,8a-dimethyl-5-methylene- | $C_{15}H_{18}O_2$ | FSTP | 76.90 | 0.89 | 740 | 29 |
| Ageratriol | $C_{15}H_{24}O_3$ | FSTP | 76.90 | 4.86 | 634 | 47 |
| α-Santalol | $C_{15}H_{24}O$ | FSTP | 78.40 | 4.13 | 612 | 47 |
| Dihydroartemisinin | $C_{15}H_{22}O_3$ | FSTP | 78.90 | 1.77 | 794 | 52 |
| Lanceol | $C_{15}H_{24}O$ | FSTP | 78.90 | 4.27 | 695 | 38 |

**Supplementary Table 3.** (Cont.)

| Compound Name | Mol. formula | Clas. | RT I (min) | RT II (sec) | MF | P (%) |
|---|---|---|---|---|---|---|
| 4,11-Dimethyl-8-(propan-2-yl)-5,12-dioxatricyclo[9.1.0.04,6]dodecan-7-ol | $C_{15}H_{26}O_3$ | FSTP | 79.00 | 4.13 | 699 | 67 |
| β-Copaenol | $C_{15}H_{24}O$ | FSTP | 79.10 | 4.07 | 595 | 27 |
| 2H-2,4a-Ethanonaphthalene, 1,3,4,5,6,7-hexahydro-2,5,5-trimethyl- | $C_{15}H_{24}$ | STP | 79.50 | 4.54 | 583 | 46 |
| Thujopsene | $C_{15}H_{24}$ | STP | 80.30 | 3.95 | 588 | 34 |
| Hinesol | $C_{15}H_{26}O$ | FSTP | 82.00 | 4.09 | 593 | 40 |
| Himachaladiene | $C_{15}H_{24}$ | STP | 83.60 | 4.15 | 716 | 31 |
| α-Selinene | $C_{15}H_{24}$ | STP | 84.00 | 4.15 | 670 | 15 |
| [5-(Hydroxymethyl)-2,5,8a-trimethyl-1,4,4a,6,7,8-hexahydronaphthalen-1-yl]methanol | $C_{15}H_{26}O_2$ | FSTP | 84.20 | 0.52 | 794 | 29 |
| 3,4,8,8-Tetramethyl-4,5,6,7,8,8a-hexahydro-1H-3a,7-methanoazulen-4-ol | $C_{15}H_{24}O$ | FSTP | 85.90 | 4.74 | 769 | 21 |
| 5β,7βH,10α-Eudesm-11-en-1α-ol | $C_{15}H_{26}O$ | FSTP | 86.60 | 4.21 | 747 | 16 |
| Limonenol | $C_{15}H_{24}O_2$ | FSTP | 86.80 | 4.36 | 633 | 15 |
| β-Santalol | $C_{15}H_{24}O$ | FSTP | 86.80 | 4.27 | 755 | 54 |
| 1,4-Methano-1H-indene, octahydro-4-methyl-8-methylene-7-(1-methylethyl)-, [1S-(1α,3aβ,4α,7α,7aβ)]- | $C_{15}H_{24}$ | STP | 86.80 | 1.71 | 671 | 42 |
| 9-(3,3-Dimethyloxiran-2-yl)-2,7-dimethylnona-2,6-dien-1-ol | $C_{15}H_{26}O_2$ | FSTP | 87.10 | 4.48 | 681 | 28 |
| Eudesmaol | $C_{15}H_{24}O$ | FSTP | 87.40 | 3.97 | 712 | 44 |
| 6S-2,3,8,8-Tetramethyltricyclo[5.2.2.0(1,6)]undec-2-ene | $C_{15}H_{24}$ | STP | 89.50 | 5.20 | 644 | 29 |
| Hydroxy albrassitriol | $C_{15}H_{26}O_4$ | FSTP | 89.70 | 2.44 | 702 | 53 |
| 1,7-Dimethyl-4-(1-methylethyl)cyclodecane | $C_{15}H_{30}$ | STP | 92.40 | 4.86 | 667 | 25 |
| Picrotoxinin | $C_{15}H_{16}O_6$ | FSTP | 94.80 | 3.69 | 625 | 84 |
| Isopetasol | $C_{15}H_{22}O_2$ | FSTP | 96.70 | 4.96 | 630 | 88 |
| 3H-Cyclodeca[b]furan-2-one, 4,9-dihydroxy-6-methyl-3,10-dimethylene-3a,4,7,8,9,10,11,11a-octahydro- | $C_{15}H_{20}O_4$ | FSTP | 100.80 | 4.29 | 681 | 44 |
| 2H-Cyclopropa[g]benzofuran, 4,5,5a,6,6a,6b-hexahydro-4,4,6b-trimethyl-2-(1-methylethenyl)- | $C_{15}H_{22}O$ | FSTP | 103.50 | 5.14 | 776 | 69 |
| Coronopilin | $C_{15}H_{20}O_4$ | FSTP | 107.30 | 5.50 | 632 | 89 |
| 3-Oxo-10(14)-epoxyguai-11(13)-en-6,12-olide | $C_{15}H_{18}O_4$ | FSTP | 110.90 | 4.36 | 750 | 30 |
| Longipinocarveol | $C_{15}H_{24}O$ | FSTP | 111.10 | 4.96 | 695 | 32 |
| 1H-Indene, 2,3,3a,4,7,7a-hexahydro-2,2,4,4,7,7-hexamethyl- | $C_{15}H_{26}$ | STP | 113.50 | 5.28 | 622 | 80 |
| 4-Hydroxy-5-(5-hydroxypentan-2-yl)-6-methyl-3-methylidene-3a,4,7,7a-tetrahydro-1-benzofuran-2-one | $C_{15}H_{22}O_4$ | FSTP | 114.50 | 4.56 | 690 | 67 |
| 2,4a,8,8-Tetramethyldecahydrocyclopropa[d]naphthalene | $C_{15}H_{26}$ | STP | 116.40 | 4.25 | 766 | 71 |

**Supplementary Table 3.** (Cont.)

| Compound Name | Mol. formula | Clas. | RT I (min) | RT II (sec) | MF | P (%) |
|---|---|---|---|---|---|---|
| γ-Himachalene | $C_{15}H_{24}$ | STP | 116.70 | 5.00 | 742 | 79 |
| Spiro[4.5]decan-7-one, 1,8-dimethyl-8,9-epoxy-4-isopropyl- | $C_{15}H_{24}O_2$ | FSTP | 119.60 | 4.80 | 597 | 54 |
| 1,5,9,9-Tetramethyl-2-oxatricyclo[6.4.0.0(4,8)]dodecane | $C_{15}H_{26}O$ | FSTP | 122.90 | 4.05 | 690 | 44 |

**Supplementary Table 4.** Terpenoid standard mixture analysed using GCxGC-TOF/MS.

| Compound Name | Molecular formula | Classification | RT I (min) | RT II (min) | MF | P (%) |
|---|---|---|---|---|---|---|
| Butanoic acid, 3-methyl-, 1-ethenyl-1,5-dimethyl-4-hexenyl ester | $C_{15}H_{26}O_2$ | Sesquiterpenoid | 12.00 | 1.07 | 726 | 73 |
| Camphene | $C_{10}H_{16}$ | Monoterpenoid | 12.50 | 1.29 | 879 | 88 |
| Bicyclo[3.1.1]heptane, 6,6-dimethyl-2-methylene-, (1S)- | $C_{10}H_{16}$ | Monoterpenoid | 13.60 | 1.43 | 780 | 78 |
| β-Myrcene | $C_{10}H_{16}$ | Monoterpenoid | 13.90 | 1.37 | 780 | 78 |
| Bicyclo[3.1.0]hex-2-ene, 4-methylene-1-(1-methylethyl)- | $C_{10}H_{14}$ | Monoterpenoid | 14.20 | 1.57 | 603 | 60 |
| Bicyclo[3.1.0]hex-2-ene, 2-methyl-5-(1-methylethyl)- | $C_{10}H_{16}$ | Monoterpenoid | 14.70 | 1.63 | 883 | 88 |
| 5-Isopropyl-2-methylbicyclo[3.1.0]hexan-2-ol # | $C_{10}H_{18}O$ | Monoterpenoid | 15.20 | 1.61 | 543 | 54 |
| p-Cymene | $C_{10}H_{14}$ | Monoterpenoid | 15.60 | 1.86 | 572 | 57 |
| Limonene | $C_{10}H_{16}$ | Monoterpenoid | 15.80 | 1.63 | 951 | 95 |
| Eucalyptol | $C_{10}H_{18}O$ | Monoterpenoid | 16.00 | 1.85 | 676 | 68 |
| 3-Carene | $C_{10}H_{16}$ | Monoterpenoid | 16.50 | 1.65 | 937 | 94 |
| γ-Terpinene | $C_{10}H_{16}$ | Monoterpenoid | 17.20 | 1.85 | 896 | 90 |
| Bicyclo[3.1.0]hexan-2-ol, 2-methyl-5-(1-methylethyl)-, (1α,2α,5α)- | $C_{10}H_{18}O$ | Monoterpenoid | 17.70 | 2.00 | 838 | 84 |
| Cyclohexene, 1-methyl-4-(1-methylethylidene)- | $C_{10}H_{16}$ | Monoterpenoid | 18.80 | 1.98 | 918 | 92 |
| Fenchone | $C_{10}H_{16}O$ | Monoterpenoid | 19.00 | 2.56 | 949 | 95 |
| 3-Octanol, 3,7-dimethyl- | $C_{10}H_{22}O$ | Monoterpenoid | 19.10 | 1.73 | 949 | 95 |
| Fenchol | $C_{10}H_{18}O$ | Monoterpenoid | 20.40 | 2.44 | 883 | 88 |
| 1-Pentene, 5-(2,2-dimethylcyclopropyl)-2-methyl-4-methylene- | $C_{12}H_{20}$ | Ketone | 21.00 | 2.64 | 592 | 59 |
| 1,2-Dihydrolinalool | $C_{10}H_{20}O$ | Monoterpenoid | 21.10 | 2.12 | 840 | 84 |
| cis-Verbenol | $C_{10}H_{16}O$ | Monoterpenoid | 21.90 | 2.72 | 789 | 79 |
| (+)-2-Bornanone | $C_{10}H_{16}O$ | Monoterpenoid | 22.20 | 3.23 | 867 | 87 |
| Cyclohexanol, 5-methyl-2-(1-methylethenyl)- | $C_{10}H_{18}O$ | Monoterpenoid | 22.20 | 2.48 | 978 | 98 |
| Cyclohexanone, 5-methyl-2-(1-methylethyl)-, cis- | $C_{10}H_{18}O$ | Monoterpenoid | 22.60 | 2.68 | 938 | 94 |
| Isoborneol | $C_{10}H_{18}O$ | Monoterpenoid | 22.90 | 2.90 | 935 | 93 |
| endo-Borneol | $C_{10}H_{18}O$ | Monoterpenoid | 23.40 | 2.96 | 946 | 95 |
| Cyclohexanol, 5-methyl-2-(1-methylethyl)-, [1S-(1α,2α,5β)]- | $C_{10}H_{20}O$ | Monoterpenoid | 23.70 | 2.50 | 764 | 76 |
| Terpinen-4-ol | $C_{10}H_{18}O$ | Monoterpenoid | 24.00 | 2.74 | 930 | 93 |

**Supplementary Table 4.** (Cont.)

| Compound Name | Molecular formula | Classification | RT I (min) | RT II (min) | MF | P (%) |
|---|---|---|---|---|---|---|
| α-Terpineol | $C_{10}H_{18}O$ | Monoterpenoid | 24.80 | 2.92 | 935 | 93 |
| Estragole | $C_{10}H_{12}O$ | Monoterpenoid | 25.20 | 3.43 | 796 | 80 |
| Bicyclo[3.1.1]hept-2-ene-2-methanol, 6,6-dimethyl- | $C_{10}H_{16}O$ | Monoterpenoid | 25.20 | 3.09 | 883 | 88 |
| Bicyclo[3.1.1]hept-3-en-2-one, 4,6,6-trimethyl- | $C_{10}H_{14}O$ | Monoterpenoid | 26.00 | 3.89 | 630 | 63 |
| 2,6-Octadiene, 1-(1-ethoxyethoxy)-3,7-dimethyl- | $C_{14}H_{26}O_2$ | Ketone | 26.80 | 2.76 | 885 | 89 |
| Cyclohexanone, 5-methyl-2-(1-methylethenyl)- | $C_{10}H_{16}O$ | Monoterpenoid | 27.80 | 3.53 | 845 | 85 |
| Carvone | $C_{10}H_{14}O$ | Monoterpenoid | 28.00 | 3.75 | 925 | 93 |
| Bicyclo[3.1.1]heptan-3-one, 2-hydroxy-2,6,6-trimethyl- | $C_{10}H_{16}O_2$ | Monoterpenoid | 28.50 | 4.17 | 529 | 53 |
| 2-Cyclohexen-1-one, 3-methyl-6-(1-methylethyl)- | $C_{10}H_{16}O$ | Monoterpenoid | 28.70 | 3.75 | 938 | 94 |
| Bicyclo[2.2.1]heptan-2-ol, 1,7,7-trimethyl-, acetate, (1S-endo)- | $C_{12}H_{20}O_2$ | Ketone | 30.70 | 2.96 | 895 | 89 |
| Phenol, 2-methyl-5-(1-methylethyl)- | $C_{10}H_{14}O$ | Monoterpenoid | 30.90 | 3.71 | 757 | 76 |
| Cyclohexanol, 5-methyl-2-(1-methylethyl)-, acetate, (1α,2α,5β)- | $C_{12}H_{22}O_2$ | Ketone | 30.90 | 2.54 | 841 | 84 |
| Phenol, 2-methyl-5-(1-methylethyl)- | $C_{10}H_{14}O$ | Monoterpenoid | 31.50 | 3.83 | 908 | 91 |
| Bicyclo[2.2.1]heptane-2,3-dione, 1,7,7-trimethyl-, (1S)- | $C_{10}H_{14}O_2$ | Monoterpenoid | 31.80 | 5.38 | 973 | 97 |
| Santolina alcohol | $C_{10}H_{18}O$ | Monoterpenoid | 31.80 | 3.97 | 929 | 93 |
| (1R,2R,3S,5R)-(-)-2,3-Pinanediol | $C_{10}H_{18}O_2$ | Monoterpenoid | 33.00 | 4.34 | 908 | 91 |
| Bicyclo[2.2.1]heptane, 2-chloro-2,3,3-trimethyl- | $C_{10}H_{17}Cl$ | Monoterpenoid | 33.40 | 2.40 | 886 | 89 |
| 6-Octen-1-ol, 3,7-dimethyl-, acetate | $C_{12}H_{22}O_2$ | Ketone | 34.30 | 2.58 | 797 | 80 |
| Geranyl isovalerate | $C_{15}H_{26}O_2$ | Sesquiterpenoid | 35.10 | 2.88 | 809 | 81 |
| 4-Hexen-1-ol, 5-methyl-2-(1-methylethenyl)-, acetate | $C_{12}H_{20}O_2$ | Ketone | 36.20 | 3.00 | 995 | 100 |
| α-Damascone | $C_{13}H_{20}O$ | Ketone | 37.20 | 3.45 | 834 | 83 |
| 1,3-Cyclohexadiene, 5-(1,5-dimethyl-4-hexenyl)-2-methyl-, [S-(R*,S*)]- | $C_{15}H_{24}$ | Sesquiterpenoid | 38.70 | 2.64 | 644 | 64 |
| Caryophyllene | $C_{15}H_{24}$ | Sesquiterpenoid | 39.10 | 2.68 | 923 | 92 |
| cis-β-Farnesene | $C_{15}H_{24}$ | Sesquiterpenoid | 40.80 | 2.40 | 901 | 90 |
| Humulene | $C_{15}H_{24}$ | Sesquiterpenoid | 41.20 | 2.84 | 903 | 90 |
| 3-Buten-2-one, 4-(2,6,6-trimethyl-1-cyclohexen-1-yl)- | $C_{13}H_{20}O$ | Ketone | 42.80 | 3.61 | 849 | 85 |
| 1,3,6,10-Dodecatetraene, 3,7,11-trimethyl-, (Z,E)- | $C_{15}H_{24}$ | Sesquiterpenoid | 43.00 | 2.58 | 877 | 88 |

**Supplementary Table 4.** (Cont.)

| Compound Name | Molecular formula | Classification | RT I (min) | RT II (min) | MF | P (%) |
|---|---|---|---|---|---|---|
| 1,3,6,10-Dodecatetraene, 3,7,11-trimethyl-, (Z,E)- | C15H24 | Sesquiterpenoid | 43.80 | 2.58 | 846 | 85 |
| (E)-β-Farnesene | C15H24 | Sesquiterpenoid | 44.00 | 2.60 | 855 | 85 |
| Butylated Hydroxytoluene | C15H24O | Sesquiterpenoid | 44.30 | 3.35 | 953 | 95 |
| 1-Isopropyl-4,7-dimethyl-1,2,3,5,6,8a-hexahydronaphthalene | C15H24 | Sesquiterpenoid | 45.10 | 2.96 | 980 | 98 |
| 1,5-Diphenyl-2H-1,2,4-triazoline-3-thione | C14H11N3S | Ketone | 46.00 | 2.74 | 885 | 89 |
| (E)-β-Farnesene | C15H24 | Sesquiterpenoid | 47.00 | 3.57 | 953 | 95 |
| 1,6,10-Dodecatrien-3-ol, 3,7,11-trimethyl-, (E)- | C15H26O | Sesquiterpenoid | 47.10 | 2.92 | 898 | 90 |
| Caryophyllene oxide | C15H24O | Sesquiterpenoid | 48.80 | 3.69 | 954 | 95 |
| Cedrol | C15H26O | Sesquiterpenoid | 50.00 | 3.87 | 838 | 84 |
| Cyclopentaneacetic acid, 3-oxo-2-(2-pentenyl)-, methyl ester, [1α,2α(Z)]- | C13H20O3 | Ketone | 52.00 | 4.72 | 957 | 96 |
| α-Bisabolol | C15H26O | Sesquiterpenoid | 54.00 | 3.53 | 970 | 97 |
| 2-Butenoic acid, 2-methyl-, 3,7-dimethyl-2,6-octadienyl ester, (E,Z)- | C15H24O2 | Sesquiterpenoid | 54.60 | 3.49 | 788 | 79 |
| 1-Methylene-2b-hydroxymethyl-3,3-dimethyl-4b-(3-methylbut-2-enyl)-cyclohexane | C15H26O | Sesquiterpenoid | 55.90 | 3.39 | 912 | 91 |
| Cyclopropane, 1-methyl-2-(3-methylpentyl)- | C10H20 | Monoterpenoid | 56.00 | 1.81 | 931 | 93 |
| Azulene, 1,4-dimethyl-7-(1-methylethyl)- | C15H18 | Sesquiterpenoid | 59.20 | 5.14 | 915 | 91 |
| Nootkatone | C15H22O | Sesquiterpenoid | 60.80 | 5.14 | 811 | 81 |
| Neophytadiene | C20H38 | Diterpenoid | 61.60 | 2.08 | 948 | 95 |
| (E,E)-7,11,15-Trimethyl-3-methylene-hexadeca-1,6,10,14-tetraene | C20H32 | Diterpenoid | 65.60 | 2.84 | 879 | 88 |
| 3,7,11,15-Tetramethyl-2-hexadecen-1-ol | C20H40O | Diterpenoid | 66.90 | 2.44 | 835 | 83 |
| 1-Methylene-2b-hydroxymethyl-3,3-dimethyl-4b-(3-methylbut-2-enyl)-cyclohexane | C15H26O | Sesquiterpenoid | 68.00 | 2.98 | 602 | 60 |
| Musk ketone | C14H18N2O5 | Ketone | 69.30 | 5.93 | 954 | 95 |
| Phytol | C20H40O | Diterpenoid | 74.50 | 2.90 | 935 | 93 |
| α-Santonin | C15H18O3 | Sesquiterpenoid | 79.60 | 3.11 | 631 | 63 |
| 2-Methyl-3-(3-methyl-but-2-enyl)-2-(4-methyl-pent-3-enyl)-oxetane | C15H26O | Sesquiterpenoid | 101.10 | 3.61 | 499 | 50 |
| Supraene | C30H50 | Triterpenoid | 101.40 | 3.55 | 881 | 88 |
| Squalene | C30H50 | Triterpenoid | 101.70 | 3.65 | 690 | 69 |

**Supplementary Table 5.** *Chlamydomonas reinhardtii* metabolism optimisation to produce heterologous sesquiterpenes.

| Construct transformed | Cell concentration (cells/L) | | | Patchoulol titer (µg/L) | | | | | Patchoulol titer (fg/cell) | | | | |
|---|---|---|---|---|---|---|---|---|---|---|---|---|---|
| | n | Mean | SD | n | Mean | SD | Fold change | p-value | n | Mean | SD | Fold change | p-value |
| PcPS_YFP | 2.1E+10 | 2.3E+10 | 4.5E+09 | 77 | 119 | 37 | - | - | 3.7 | 5.4 | 2.1 | - | - |
| | 2.1E+10 | | | 109 | | | | | 5.3 | | | | |
| | 2.0E+10 | | | 165 | | | | | 8.4 | | | | |
| | 2.9E+10 | | | 125 | | | | | 4.3 | | | | |
| PcPS_YFP + CrBKT | 2.2E+10 | 2.2E+10 | 2.1E+08 | 969 | 906 | 126 | 8 | 0.0003 | 44.6 | 41.2 | 5.9 | 8 | 0.0114 |
| | 2.2E+10 | | | 993 | | | | | 45.3 | | | | |
| | 2.2E+10 | | | 719 | | | | | 32.6 | | | | |
| | 2.2E+10 | | | 941 | | | | | 42.4 | | | | |
| PcPS_YFP + SQSkd | 2.2E+10 | 2.3E+10 | 2.1E+08 | 1502 | 1622 | 174 | 14 | <0.0001 | 67.1 | 71.7 | 7.8 | 12 | 0.0110 |
| | 2.3E+10 | | | 1809 | | | | | 80.3 | | | | |
| | 2.3E+10 | | | 1729 | | | | | 76.2 | | | | |
| | 2.3E+10 | | | 1449 | | | | | 63.4 | | | | |
| PcPS_YFP + CrBKT + SQSkd | 2.3E+10 | 2.8E+10 | 3.7E+09 | 2171 | 2215 | 272 | 21 | <0.0001 | 110.7 | 132.7 | 41.1 | 25 | <0.0001 |
| | 2.6E+10 | | | 2385 | | | | | 156.4 | | | | |
| | 2.9E+10 | | | 2074 | | | | | 87.1 | | | | |
| | 3.1E+10 | | | 2231 | | | | | 176.7 | | | | |

**Supplementary Table 6.** List of plasmids.

| Construct number | Construct name | Gene name | Product | Gene length (bp) | Protein size (kDa) | Intron copies (i1RBCS2) | Reporter (FP) | Selection (Antibiotic Resistance) |
|---|---|---|---|---|---|---|---|---|
| 01 | P007 | *mVenus* | Yellow fluorescent protein | 1004 | 27 | 3 | YFP | Paromomycin |
| 02 | P117 | *Sa*SS | Santalene | 2441 | 92 | 5 | YFP | Paromomycin |
| 03 | P118 | *Cl*SS | Santalene | 2387 | 91 | 5 | YFP | Paromomycin |
| 04 | P119 | *Cc*SS | Santalene | 2393 | 91 | 5 | YFP | Paromomycin |
| 05 | P124 | *Cz*ZS | Zizaene | 2399 | 91 | 4 | YFP | Paromomycin |
| 06 | P125 | *Cz*ZS | Zizaene | 2390 | 91 | 4 | YFP | Paromomycin |
| 07 | P107 | *Pe*AS | Aristolochene | 1446 | 65 | 3 | YFP | Paromomycin |
| 08 | P108 | *Pc*AS | Aristolochene | 1470 | 66 | 3 | YFP | Paromomycin |
| 09 | P109 | *Pc*AS | Aristolochene | 1521 | 68 | 3 | YFP | Paromomycin |
| 10 | P110 | *Pr*AS | Aristolochene | 1470 | 66 | 3 | YFP | Paromomycin |
| 11 | P120 | *Cs*VS | Valencene | 2378 | 91 | 5 | YFP | Paromomycin |
| 12 | P121 | *Cn*VS | Valencene | 2501 | 96 | 5 | YFP | Paromomycin |
| 13 | P111 | *Ac*DGSI | δ-guaiene | 2375 | 91 | 5 | YFP | Paromomycin |
| 14 | P112 | *Ac*DGSII | δ-guaiene | 2375 | 91 | 5 | YFP | Paromomycin |
| 15 | P113 | *Ac*DGSIII | δ-guaiene | 2375 | 91 | 5 | YFP | Paromomycin |
| 16 | P114 | *As*DGSI | δ-guaiene | 2375 | 95 | 5 | YFP | Paromomycin |
| 17 | P115 | *As*DGSII | δ-guaiene | 2375 | 91 | 5 | YFP | Paromomycin |
| 18 | P116 | *As*DGSIII | δ-guaiene | 2375 | 91 | 5 | YFP | Paromomycin |
| 19 | P126 | *La*CS | $\tau$-Cadinol | 2426 | 92 | 5 | YFP | Paromomycin |
| 20 | P122 | *Cs*VL | Valerioanol | 2453 | 94 | 5 | YFP | Paromomycin |
| 21 | P123 | *Ch*VL | Valerioanol | 2396 | 91 | 5 | YFP | Paromomycin |
| 22 | P139 | ispA_*Pc*PS(C415F_H454A) | Patchoulol | 3707 | 122 | 8 | YFP | Paromomycin |
| 23 | P140 | *Pc*PS(C415F_H454A) | Patchoulol | 2378 | 72 | 6 | YFP | Paromomycin |
| 24 | P127 | *Cc*BS | Bisabolol | 2468 | 93 | 5 | YFP | Paromomycin |
| 25 | B005 | mTFP1 | Cyan-green florescent protein | 1140 | 27 | 3 | TFP | Bleomycin |
| 26 | B036 | *ClSS* | Santalene | 1650 | 91 | 5 | BFP | Bleomycin |
| 27 | B039 | *CzZS* | Zizaene | 2399 | 91 | 4 | BFP | Bleomycin |
| 28 | B033 | *PcAS* | Aristolochene | 1078 | 67 | 3 | BFP | Bleomycin |
| 29 | B037 | *CnVS* | Valencene | 1764 | 96 | 5 | BFP | Bleomycin |

**Supplementary Table 6.** (Cont.)

| Construct number | Construct name | Gene name | Product | Gene length (bp) | Protein size (kDa) | Intron copies (i1RBCS2) | Reporter (FP) | Selection (Antibiotic Resistance) |
|---|---|---|---|---|---|---|---|---|
| 30 | B034 | *AcDGSI* | δ-guaiene | 1638 | 91 | 5 | BFP | Bleomycin |
| 31 | B035 | *AsDGSIII* | δ-guaiene | 1638 | 91 | 5 | BFP | Bleomycin |
| 32 | B040 | *LaCS* | τ-Cadinol | 1662 | 92 | 5 | BFP | Bleomycin |
| 33 | B038 | *ChVL* | Valerioanol | 1659 | 91 | 5 | BFP | Bleomycin |
| 34 | B031 | ispA_*Pc*PS(C415F_H454A) | Patchoulol | 3707 | 122 | 8 | BFP | Bleomycin |
| 35 | B032 | *Pc*PS(C415F_H454A) | Patchoulol | 2378 | 72 | 6 | BFP | Bleomycin |
| 36 | B041 | *Cc*BS | Bisabolol | 1704 | 93 | 5 | BFP | Bleomycin |
| 37 | P011 | *Pc*PS | Patchoulol | 2097 | 72 | 5 | YFP | Paromomycin |
| 38 | MS021 | BKT | Ketocarotenoids | 2419 | 93 | 3 | - | Spectinomycin |
| 39 | S001 | SQSkd | Squalene synthase knock-down | 1382 | 30 | 3 | gLuc | Spectinomycin |

**Supplementary Table 7.** Sesquiterpenoid quantifications (Single transformation).

| Sesquiterpenoid | Construct transformed | Cell concentration (cells/L) | | | Product titer ($\mu$g/L culture) | | | Product titer (fg/cell) | | |
|---|---|---|---|---|---|---|---|---|---|---|
| | | n | Mean | SD | n | Mean | SD | n | Mean | SD |
| Santalene | 3 | 2.3E+10 | 2.3E+10 | 2.1E+09 | 78 | 64 | 10 | 3.4 | 2.7 | 0.5 |
| | | 2.1E+10 | | | 55 | | | 2.7 | | |
| | | 2.6E+10 | | | 58 | | | 2.3 | | |
| | | 2.4E+10 | | | 63 | | | 2.6 | | |
| Zizaene | 6 | 2.5E+10 | 2.5E+10 | 4.1E+09 | 503 | 416 | 77 | 19.8 | 16.6 | 2.5 |
| | | 3.1E+10 | | | 448 | | | 14.5 | | |
| | | 2.2E+10 | | | 324 | | | 14.7 | | |
| | | 2.2E+10 | | | 390 | | | 17.4 | | |
| Aristolochene | 9 | 1.9E+10 | 1.7E+10 | 2.4E+09 | 32 | 25 | 6 | 1.6 | 1.4 | 0.2 |
| | | 1.7E+10 | | | 29 | | | 1.7 | | |
| | | 1.9E+10 | | | 22 | | | 1.2 | | |
| | | 1.4E+10 | | | 18 | | | 1.3 | | |
| Valencene | 12 | 2.3E+10 | 2.3E+10 | 0.0E+00 | 57 | 52 | 5 | 2.4 | 2.5 | 0.1 |
| | | 2.3E+10 | | | 54 | | | 2.5 | | |
| | | 2.3E+10 | | | 52 | | | 2.6 | | |
| | | 2.3E+10 | | | 46 | | | 2.5 | | |
| δ-Guaiene | 13 | 2.1E+10 | 2.1E+10 | 3.4E+08 | 26 | 25 | 4 | 1.3 | 1.2 | 0.2 |
| | | 2.1E+10 | | | 30 | | | 1.4 | | |
| | | 2.1E+10 | | | 21 | | | 1.0 | | |
| | | 2.1E+10 | | | 22 | | | 1.0 | | |
| δ-Guaiene | 18 | 2.1E+10 | 2.5E+10 | 3.2E+09 | 1747 | 1447 | 312 | 83.6 | 60.3 | 20.4 |
| | | 2.4E+10 | | | 1668 | | | 70.9 | | |
| | | 2.7E+10 | | | 1279 | | | 46.8 | | |
| | | 2.7E+10 | | | 1094 | | | 39.9 | | |
| τ-Cadinol | 19 | 2.1E+10 | 2.4E+10 | 2.6E+09 | 30 | 43 | 14 | 1.4 | 1.8 | 0.6 |
| | | 2.7E+10 | | | 37 | | | 1.4 | | |
| | | 2.2E+10 | | | 43 | | | 2.0 | | |
| | | 2.4E+10 | | | 62 | | | 2.6 | | |
| Valerianol | 21 | 2.2E+10 | 2.0E+10 | 1.9E+09 | 40 | 41 | 13 | 1.8 | 2.1 | 0.6 |
| | | 1.8E+10 | | | 44 | | | 2.4 | | |
| | | 2.0E+10 | | | 56 | | | 2.7 | | |
| | | 1.8E+10 | | | 25 | | | 1.4 | | |
| Patchoulol | 22 | 2.4E+10 | 2.1E+10 | 2.6E+09 | 1813 | 1729 | 132 | 77.1 | 82.1 | 4.4 |
| | | 2.3E+10 | | | 1871 | | | 80.4 | | |
| | | 1.9E+10 | | | 1618 | | | 83.5 | | |
| | | 1.8E+10 | | | 1614 | | | 87.5 | | |
| Bisabolol | 24 | 2.2E+10 | 2.2E+10 | 9.6E+08 | 28 | 21 | 5 | 1.3 | 1.0 | 0.3 |
| | | 2.1E+10 | | | 23 | | | 1.1 | | |
| | | 2.3E+10 | | | 19 | | | 0.8 | | |
| | | 2.3E+10 | | | 15 | | | 0.6 | | |

**Supplementary Table 8.** Sesquiterpenoid quantifications (Double transformation).

| Sesquiterpenoid | Construct transformed | Cell concentration (cells/L) | | | Product titer ($\mu$g/L culture) | | | Product titer (fg/cell) | | |
|---|---|---|---|---|---|---|---|---|---|---|
| | | n | Mean | SD | n | Mean | SD | n | Mean | SD |
| Santalene | 3+26 | 2.1E+10 | 2.4E+10 | 2.7E+09 | 211 | 250 | 27 | 10.0 | 11 | 1.1 |
| | | 2.7E+10 | | | 257 | | | 9.4 | | |
| | | 2.4E+10 | | | 269 | | | 11.3 | | |
| | | 2.2E+10 | | | 265 | | | 11.9 | | |
| Zizaene | 6+27 | 2.4E+10 | 2.4E+10 | 6.0E+08 | 739 | 642 | 80 | 30.4 | 26 | 3.0 |
| | | 2.4E+10 | | | 625 | | | 25.9 | | |
| | | 2.5E+10 | | | 659 | | | 26.2 | | |
| | | 2.4E+10 | | | 547 | | | 23.1 | | |
| Aristolochene | 9+28 | 2.2E+10 | 1.4E+10 | 6.0E+09 | 36 | 38 | 7 | 1.6 | 3 | 1.7 |
| | | 1.4E+10 | | | 39 | | | 2.8 | | |
| | | 8.4E+09 | | | 47 | | | 5.5 | | |
| | | 1.2E+10 | | | 30 | | | 2.6 | | |
| Valencene | 12+29 | 2.4E+10 | 2.1E+10 | 2.5E+09 | 332 | 313 | 17 | 14.3 | 13 | 0.7 |
| | | 2.2E+10 | | | 309 | | | 13.3 | | |
| | | 2.0E+10 | | | 292 | | | 12.5 | | |
| | | 1.8E+10 | | | 319 | | | 13.7 | | |
| δ-Guaiene | 13+30 | 2.1E+10 | 2.2E+10 | 9.3E+08 | 54 | 53 | 13 | 2.5 | 2 | 0.6 |
| | | 2.2E+10 | | | 67 | | | 3.0 | | |
| | | 2.3E+10 | | | 57 | | | 2.5 | | |
| | | 2.3E+10 | | | 36 | | | 1.6 | | |
| δ-Guaiene | 18+31 | 2.3E+10 | 2.4E+10 | 1.8E+09 | 3236 | 2692 | 482 | 137.8 | 113 | 25.3 |
| | | 2.7E+10 | | | 2092 | | | 78.1 | | |
| | | 2.3E+10 | | | 2861 | | | 122.0 | | |
| | | 2.3E+10 | | | 2580 | | | 113.0 | | |
| τ-Cadinol | 19+32 | 2.6E+10 | 2.5E+10 | 8.7E+08 | 374 | 399 | 36 | 14.5 | 16 | 1.2 |
| | | 2.5E+10 | | | 434 | | | 17.1 | | |
| | | 2.4E+10 | | | 363 | | | 14.9 | | |
| | | 2.6E+10 | | | 425 | | | 16.1 | | |
| Valerianol | 21+33 | 2.6E+10 | 2.4E+10 | 4.0E+09 | 446 | 538 | 85 | 17.1 | 23 | 7.2 |
| | | 2.9E+10 | | | 494 | | | 17.3 | | |
| | | 2.1E+10 | | | 640 | | | 31.0 | | |
| | | 2.1E+10 | | | 573 | | | 27.8 | | |
| Patchoulol | 22+34 | 2.6E+10 | 2.4E+10 | 2.6E+09 | 3007 | 2767 | 201 | 116.9 | 115 | 9.3 |
| | | 2.4E+10 | | | 2578 | | | 107.7 | | |
| | | 2.1E+10 | | | 2625 | | | 127.2 | | |
| | | 2.7E+10 | | | 2858 | | | 107.6 | | |
| Bisabolol | 24+35 | 2.2E+10 | 2.4E+10 | 1.5E+09 | 330 | 275 | 48 | 15.0 | 12 | 2.6 |
| | | 2.5E+10 | | | 293 | | | 11.7 | | |
| | | 2.3E+10 | | | 256 | | | 11.1 | | |
| | | 2.5E+10 | | | 219 | | | 8.8 | | |

**Supplementary Table 9.** Summary of sesquiterpenoids quantification.

| Sesquiterpenoid | Titer ($\mu$g/cell) | | | | Titer (fg/cell) | | | | Fold change |
|---|---|---|---|---|---|---|---|---|---|
| | Single transformation | | Double transformation | | Single transformation | | Double transformation | | |
| | Mean | SD | Mean | SD | Mean | SD | mean | SD | |
| Santalene | 63.5 | 10.4 | 250.5 | 26.9 | 2.7 | 0.5 | 10.6 | 1.1 | 3.9 |
| Zizaene | 416.3 | 76.8 | 642.3 | 79.5 | 16.6 | 2.5 | 26.4 | 3.0 | 1.6 |
| Aristolochene | 25.1 | 6.3 | 38.1 | 6.8 | 1.4 | 0.2 | 3.1 | 1.7 | 2.2 |
| Valencene | 52.4 | 4.7 | 313.1 | 16.9 | 2.5 | 0.1 | 13.4 | 0.7 | 5.4 |
| δ-Guaiene | 1447.0 | 311.5 | 2692.4 | 482.2 | 60.3 | 20.4 | 112.7 | 25.3 | 1.9 |
| τ-Cadinol | 43.2 | 13.9 | 398.8 | 35.8 | 1.8 | 0.6 | 15.7 | 1.2 | 8.7 |
| Valerianol | 41.3 | 12.5 | 538.2 | 85.4 | 2.1 | 0.6 | 23.3 | 7.2 | 11.1 |
| Patchoulol | 1729.1 | 132.4 | 2766.9 | 201.4 | 82.1 | 4.4 | 114.8 | 9.3 | 1.4 |
| Bisabolol | 21.2 | 5.5 | 274.9 | 47.8 | 1.0 | 0.3 | 11.7 | 2.6 | 11.7 |

**Supplementary Table 10**. Concentrated algal-produced sesquiterpenoids identified in ethanol using GCxGC-TOF/MS.

| Compound Name | Molecular formula | Classification | RT I (min) | RT II (min) | MF | P (%) |
|---|---|---|---|---|---|---|
| α-Santalene | $C_{15}H_{24}$ | STP | 31.20 | 2.72 | 888 | 89 |
| Aromadendrene | $C_{15}H_{22}$ | STP | 34.70 | 2.64 | 821 | 82 |
| α-Cubebene | $C_{15}H_{24}$ | STP | 36.30 | 2.42 | 752 | 75 |
| β-Bourbonene | $C_{15}H_{24}$ | STP | 36.90 | 2.54 | 718 | 72 |
| β-Elemene | $C_{15}H_{24}$ | STP | 37.20 | 2.50 | 728 | 73 |
| β-copaene | $C_{15}H_{24}$ | STP | 37.30 | 2.42 | 775 | 78 |
| α-gurjunene | $C_{15}H_{24}$ | STP | 38.20 | 2.66 | 744 | 74 |
| Isocaryophyllene | $C_{15}H_{24}$ | STP | 39.10 | 2.76 | 737 | 74 |
| Caryophyllene | $C_{15}H_{24}$ | STP | 39.30 | 2.68 | 752 | 75 |
| Seychellene | $C_{15}H_{24}$ | STP | 40.60 | 3.15 | 758 | 76 |
| β-Santalene | $C_{15}H_{24}$ | STP | 41.30 | 2.74 | 695 | 69 |
| Alloaromadendrene | $C_{15}H_{24}$ | STP | 41.30 | 2.98 | 739 | 74 |
| Zizaene | $C_{15}H_{24}$ | STP | 41.40 | 3.00 | 761 | 76 |
| Cadinene | $C_{15}H_{28}$ | STP | 42.10 | 2.38 | 904 | 90 |
| α-Curcumene | $C_{15}H_{22}$ | STP | 42.50 | 3.04 | 815 | 82 |
| α-Farnesene | $C_{15}H_{24}$ | STP | 42.90 | 3.11 | 910 | 91 |
| β-Guaiene | $C_{15}H_{24}$ | STP | 43.10 | 3.04 | 765 | 76 |
| Aristolochene | $C_{15}H_{24}$ | STP | 43.10 | 3.00 | 774 | 77 |
| Eremophilene | $C_{15}H_{24}$ | STP | 43.10 | 2.98 | 797 | 80 |
| Rosiglitazone | $C_{15}H_{24}$ | STP | 44.60 | 3.09 | 693 | 69 |
| Humulene | $C_{15}H_{24}$ | STP | 47.90 | 3.91 | 823 | 82 |
| α-Santalol | $C_{15}H_{24}O$ | FSTP | 49.50 | 4.72 | 774 | 77 |
| α-Guaiene | $C_{15}H_{24}$ | STP | 50.40 | 2.60 | 748 | 75 |
| γ-himachalene | $C_{15}H_{24}$ | STP | 51.10 | 4.17 | 812 | 81 |
| β-Caryophyllene | $C_{15}H_{24}$ | STP | 52.10 | 4.09 | 791 | 79 |
| β-Longipinene | $C_{15}H_{24}$ | STP | 53.20 | 4.33 | 591 | 59 |
| α-Santalol | $C_{15}H_{24}O$ | FSTP | 53.70 | 3.85 | 828 | 83 |

**Supplementary Table 10.** (Cont.)

| Compound Name | Molecular formula | Classification | RT I (min) | RT II (min) | MF | P (%) |
|---|---|---|---|---|---|---|
| α-copaene | $C_{15}H_{24}O$ | FSTP | 53.90 | 4.44 | 690 | 69 |
| α-Eudesmol | $C_{15}H_{24}O$ | FSTP | 54.60 | 4.21 | 702 | 70 |
| Valencene | $C_{15}H_{24}$ | STP | 54.90 | 4.21 | 814 | 81 |
| Elemol | $C_{15}H_{24}O$ | FSTP | 57.30 | 5.73 | 805 | 81 |
| δ-Guaiene | $C_{15}H_{24}$ | STP | 59.67 | 3.75 | 779 | 78 |
| Nootkatone | $C_{15}H_{22}O$ | FSTP | 61.00 | 5.28 | 830 | 83 |
| α-Patchoulene | $C_{15}H_{24}$ | STP | 61.40 | 3.04 | 737 | 74 |
| τ-Cadinol | $C_{15}H_{24}O$ | FSTP | 62.10 | 3.71 | 843 | 84 |
| γ-gurjunene | $C_{15}H_{24}$ | STP | 64.30 | 5.20 | 894 | 89 |
| Valerianol | $C_{15}H_{24}O$ | FSTP | 64.70 | 4.36 | 658 | 66 |
| α-Selinene | $C_{15}H_{24}O$ | FSTP | 69.60 | 6.19 | 949 | 95 |
| Patchouli alcohol | $C_{15}H_{24}O$ | FSTP | 73.20 | 4.52 | 865 | 86 |
| α-bisabol | $C_{15}H_{24}$ | STP | 78.30 | 2.92 | 866 | 87 |
| α-Bisabolol oxide B | $C_{15}H_{26}O_2$ | FSTP | 81.00 | 3.55 | 676 | 68 |
| δ-Elemene | $C_{15}H_{24}$ | STP | 82.80 | 4.01 | 865 | 86 |

**Supplementary Table 11.** Sesquiterpenoids identified after hydroboration-oxidation reaction using GCxGC-TOF/MS.

| Compound Name | Molecular formula | Classification | RT I (min) | RT II (min) | MF | P (%) |
|---|---|---|---|---|---|---|
| α-Bisabolol oxide B | $C_{15}H_{26}O_2$ | FSTP | 31.20 | 2.72 | 888 | 89 |
| α-Santalene | $C_{15}H_{24}$ | STP | 31.20 | 2.72 | 834 | 83 |
| α-cedrene | $C_{15}H_{24}$ | STP | 34.00 | 2.04 | 1005 | 101 |
| Zizaene | $C_{15}H_{24}$ | STP | 34.40 | 3.00 | 883 | 88 |
| Aromadendrene | $C_{15}H_{22}$ | STP | 34.70 | 2.64 | 888 | 89 |
| β-Vatirenene | $C_{15}H_{22}$ | STP | 35.80 | 3.85 | 728 | 73 |
| α-Cubebene | $C_{15}H_{24}$ | STP | 36.30 | 2.42 | 775 | 78 |
| β-Bourbonene | $C_{15}H_{24}$ | STP | 36.90 | 2.54 | 744 | 74 |
| β-Elemene | $C_{15}H_{24}$ | STP | 37.20 | 2.50 | 737 | 74 |
| Germacrene A | $C_{15}H_{24}$ | STP | 37.30 | 2.44 | 752 | 75 |
| β-copaene | $C_{15}H_{24}$ | STP | 37.30 | 2.42 | 758 | 76 |
| Isocaryophyllene | $C_{15}H_{24}$ | STP | 39.10 | 2.76 | 910 | 91 |
| Caryophyllene | $C_{15}H_{24}$ | STP | 39.30 | 2.68 | 797 | 80 |
| Patchouladiene | $C_{15}H_{22}$ | STP | 39.50 | 3.04 | 693 | 69 |
| Cycloisolongifolol | $C_{15}H_{24}O$ | FSTP | 40.40 | 2.90 | 812 | 81 |
| Seychellene | $C_{15}H_{24}$ | STP | 40.60 | 3.15 | 591 | 59 |
| Humulene | $C_{15}H_{24}$ | STP | 40.90 | 3.91 | 702 | 70 |
| β-Selinene | $C_{15}H_{24}$ | STP | 40.90 | 2.58 | 805 | 81 |
| β-Santalene | $C_{15}H_{24}$ | STP | 41.30 | 2.74 | 779 | 78 |
| Alloaromadendrene | $C_{15}H_{24}$ | STP | 41.30 | 2.98 | 843 | 84 |
| α-Patchoulene | $C_{15}H_{24}$ | STP | 41.40 | 3.04 | 658 | 66 |
| α-gurjunene | $C_{15}H_{24}$ | STP | 41.40 | 3.00 | 761 | 76 |
| Cadinene | $C_{15}H_{28}$ | STP | 42.10 | 2.38 | 804 | 80 |
| α-Farnesene | $C_{15}H_{24}$ | STP | 42.90 | 3.11 | 811 | 81 |
| β-Guaiene | $C_{15}H_{24}$ | STP | 43.10 | 3.04 | 813 | 81 |
| α-coastal | $C_{15}H_{22}O$ | FSTP | 43.10 | 3.00 | 895 | 89 |

**Supplementary Table 11.** (Cont.)

| Compound Name | Molecular formula | Classification | RT I (min) | RT II (min) | MF | P (%) |
|---|---|---|---|---|---|---|
| Aristolochene | $C_{15}H_{24}$ | STP | 43.10 | 3.00 | 815 | 82 |
| Eremophilene | $C_{15}H_{24}$ | STP | 43.10 | 2.98 | 768 | 77 |
| α-himachalene | $C_{15}H_{20}$ | FSTP | 43.30 | 3.59 | 783 | 78 |
| β-Bisabolenol | $C_{15}H_{24}O$ | FSTP | 43.30 | 3.21 | 763 | 76 |
| Eudesmatriene | $C_{15}H_{22}$ | STP | 43.40 | 3.17 | 821 | 82 |
| α-agorofuran | $C_{15}H_{24}O$ | FSTP | 46.90 | 3.99 | 838 | 84 |
| Longifolene | $C_{15}H_{20}$ | FSTP | 47.20 | 3.69 | 841 | 84 |
| Aromadendran | $C_{15}H_{26}$ | STP | 47.40 | 4.25 | 842 | 84 |
| Neoisolongifolene | $C_{15}H_{22}$ | STP | 49.30 | 2.72 | 848 | 85 |
| α-Santalol | $C_{15}H_{24}O$ | FSTP | 49.50 | 4.72 | 849 | 85 |
| isoledene | $C_{15}H_{24}$ | STP | 50.10 | 3.47 | 852 | 85 |
| Humulene epoxide I | $C_{15}H_{24}O$ | FSTP | 50.50 | 3.87 | 856 | 86 |
| γ-himachalene | $C_{15}H_{24}$ | STP | 51.10 | 4.17 | 859 | 86 |
| δ-Guaiene | $C_{15}H_{24}$ | STP | 52.00 | 3.75 | 864 | 86 |
| β-Caryophyllene | $C_{15}H_{24}$ | STP | 52.10 | 4.09 | 866 | 87 |
| τ-Cadinol | $C_{15}H_{26}O$ | FSTP | 52.10 | 3.71 | 867 | 87 |
| α-Guaiene | $C_{15}H_{24}$ | STP | 52.30 | 3.85 | 872 | 87 |
| Aristolene | $C_{15}H_{24}$ | STP | 52.50 | 4.19 | 875 | 88 |
| Agarospirol | $C_{15}H_{26}O$ | FSTP | 52.80 | 4.07 | 877 | 88 |
| Alloaromadendrenol | $C_{15}H_{26}O$ | FSTP | 52.80 | 4.07 | 986 | 99 |
| Globulol | $C_{15}H_{26}O$ | FSTP | 52.80 | 4.31 | 880 | 88 |
| Cubenol | $C_{15}H_{26}O$ | FSTP | 52.90 | 4.13 | 881 | 88 |
| Ledol | $C_{15}H_{26}O$ | FSTP | 52.90 | 5.81 | 884 | 88 |
| Guaiol | $C_{15}H_{26}O$ | FSTP | 53.00 | 4.09 | 886 | 89 |
| β-Longipinene | $C_{15}H_{24}$ | STP | 53.20 | 4.33 | 892 | 89 |
| Cedrol | $C_{15}H_{26}O$ | FSTP | 53.30 | 4.33 | 894 | 89 |
| α-Santalol | $C_{15}H_{24}O$ | FSTP | 53.70 | 3.85 | 898 | 90 |

**Supplementary Table 11.** (Cont.)

| Compound Name | Molecular formula | Classification | RT I (min) | RT II (min) | MF | P (%) |
|---|---|---|---|---|---|---|
| α.Eudesmol | $C_{15}H_{24}O$ | FSTP | 54.60 | 4.21 | 904 | 90 |
| Valerenol | $C_{15}H_{24}O$ | FSTP | 54.70 | 4.36 | 851 | 85 |
| Valerianol | $C_{15}H_{24}O$ | FSTP | 54.70 | 4.36 | 868 | 87 |
| α-Curcumene | $C_{15}H_{22}$ | STP | 54.90 | 4.21 | 750 | 75 |
| Valencene | $C_{15}H_{24}$ | STP | 54.90 | 4.21 | 911 | 91 |
| Agidol | $C_{15}H_{22}O$ | FSTP | 57.00 | 5.00 | 990 | 99 |
| Elemol | $C_{15}H_{26}O$ | FSTP | 57.30 | 5.73 | 928 | 93 |
| Eudesmenol | $C_{15}H_{26}O$ | FSTP | 57.60 | 5.46 | 933 | 93 |
| Aromadendrenol | $C_{15}H_{26}O$ | FSTP | 58.00 | 3.59 | 977 | 98 |
| Valerenic acid | $C_{15}H_{22}O_2$ | FSTP | 58.60 | 5.73 | 937 | 94 |
| γ-Gurjunenepoxide | $C_{15}H_{24}O$ | FSTP | 59.50 | 4.64 | 943 | 94 |
| α-Hexylcinnamaldehyde | $C_{15}H_{20}O$ | FSTP | 59.67 | 3.75 | 930 | 93 |
| Ylangenal | $C_{15}H_{22}O$ | FSTP | 60.40 | 5.77 | 949 | 95 |
| Nootkatone | $C_{15}H_{22}O$ | FSTP | 61.00 | 5.28 | 951 | 95 |
| β-acoradienol | $C_{15}H_{24}O$ | FSTP | 61.00 | 4.23 | 953 | 95 |
| Caryophylladienol | $C_{15}H_{26}O$ | FSTP | 61.10 | 6.01 | 957 | 96 |
| β-Eudesmol | $C_{15}H_{26}O$ | FSTP | 61.60 | 5.12 | 959 | 96 |
| α-isocomene | $C_{15}H_{24}$ | STP | 62.10 | 3.71 | 909 | 91 |
| Alloaromadendrene oxide | $C_{15}H_{24}O$ | FSTP | 63.00 | 5.08 | 968 | 97 |
| Isoaromadendrene epoxide | $C_{15}H_{24}O$ | FSTP | 63.50 | 5.24 | 864 | 86 |
| γ-gurjunene | $C_{15}H_{24}$ | STP | 64.30 | 5.20 | 982 | 98 |
| α-cyperone | $C_{15}H_{20}O$ | FSTP | 64.70 | 4.36 | 923 | 92 |
| Curcumenol | $C_{15}H_{28}O$ | FSTP | 64.70 | 4.07 | 988 | 99 |
| Bourbonenol | $C_{15}H_{26}O$ | FSTP | 65.00 | 3.21 | 989 | 99 |
| Cumanin | $C_{15}H_{22}O_4$ | FSTP | 66.50 | 1.88 | 988 | 99 |
| α-Panasinsen | $C_{15}H_{24}$ | STP | 66.80 | 0.62 | 991 | 99 |
| Valerenolic acid | $C_{15}H_{22}O_3$ | FSTP | 67.00 | 5.57 | 935 | 94 |

**Supplementary Table 11.** (Cont.)

| Compound Name | Molecular formula | Classification | RT I (min) | RT II (min) | MF | P (%) |
|---|---|---|---|---|---|---|
| Aromadendrene oxide | $C_{15}H_{24}O$ | FSTP | 67.30 | 5.85 | 940 | 94 |
| γ-Elemene | $C_{15}H_{24}$ | STP | 69.10 | 5.83 | 954 | 95 |
| Humulenol-II | $C_{15}H_{24}O$ | FSTP | 69.40 | 6.17 | 975 | 97 |
| α-selinene | $C_{15}H_{24}O$ | FSTP | 69.60 | 6.19 | 951 | 95 |
| Farnesenol | $C_{15}H_{26}O$ | FSTP | 69.60 | 6.19 | 970 | 97 |
| Nootkaton epoxide | $C_{15}H_{22}O_2$ | FSTP | 71.70 | 1.09 | 922 | 92 |
| α-Eudesmol | $C_{15}H_{22}O_2$ | FSTP | 72.40 | 1.03 | 961 | 96 |
| α-copaene | $C_{15}H_{24}O$ | FSTP | 73.20 | 4.52 | 898 | 90 |
| Gurjunenol | $C_{15}H_{28}O$ | FSTP | 73.20 | 4.52 | 981 | 98 |
| Patchouli alcohol | $C_{15}H_{24}O$ | FSTP | 73.20 | 4.52 | 865 | 86 |
| Hydroxymuurolene | $C_{15}H_{24}O$ | FSTP | 73.60 | 6.33 | 958 | 96 |
| Cedrandiol | $C_{15}H_{26}O_2$ | FSTP | 73.90 | 1.41 | 958 | 96 |
| Copaenol | $C_{15}H_{26}O$ | FSTP | 78.00 | 4.07 | 980 | 98 |
| α-bisabol | $C_{15}H_{24}$ | STP | 78.30 | 2.92 | 866 | 87 |
| Santalenol | $C_{15}H_{26}O$ | FSTP | 78.30 | 2.92 | 985 | 99 |
| Cadinol | $C_{15}H_{26}O$ | FSTP | 81.00 | 3.17 | 987 | 99 |
| Santalol | $C_{15}H_{26}O$ | FSTP | 81.00 | 3.55 | 976 | 98 |
| Caryophyllene oxide | $C_{15}H_{24}O$ | FSTP | 82.40 | 3.65 | 950 | 95 |
| Seychellene alcohol | $C_{15}H_{22}O$ | FSTP | 82.80 | 4.01 | 984 | 98 |
| δ-Elemene | $C_{15}H_{24}$ | STP | 82.80 | 4.01 | 951 | 95 |
| Ylangenol | $C_{15}H_{24}O$ | FSTP | 83.10 | 2.98 | 966 | 97 |
| Zizaenol | $C_{15}H_{26}O$ | FSTP | 85.20 | 4.58 | 986 | 99 |
| Caryophyllene alcohol | $C_{15}H_{26}O$ | FSTP | 86.00 | 4.19 | 982 | 98 |

**Supplementary Table 12.** Mass spectra of sesquiterpenoids identified by GC – MS.

| Compound Name | Molecular formula | Structure | Mass spectra |
|---|---|---|---|
| Santalene | $C_{15}H_{24}$ | 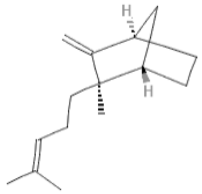 | 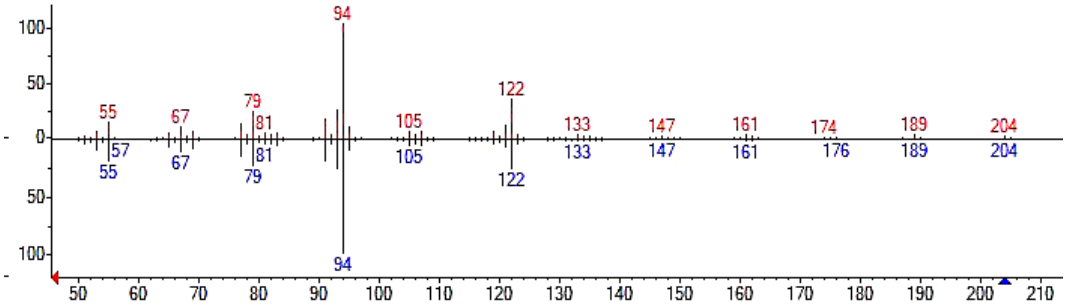 |
| Bergamotene | $C_{15}H_{24}$ | 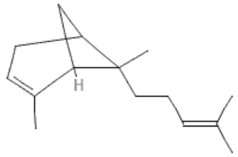 | 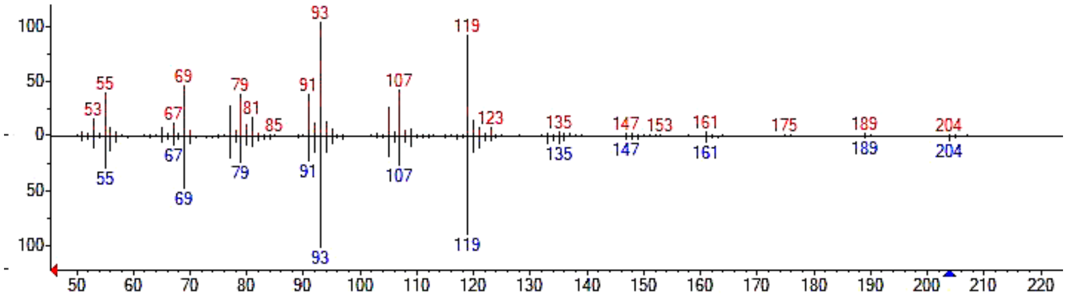 |
| Zizaene | $C_{15}H_{24}$ | 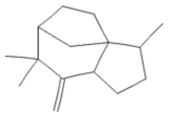 | 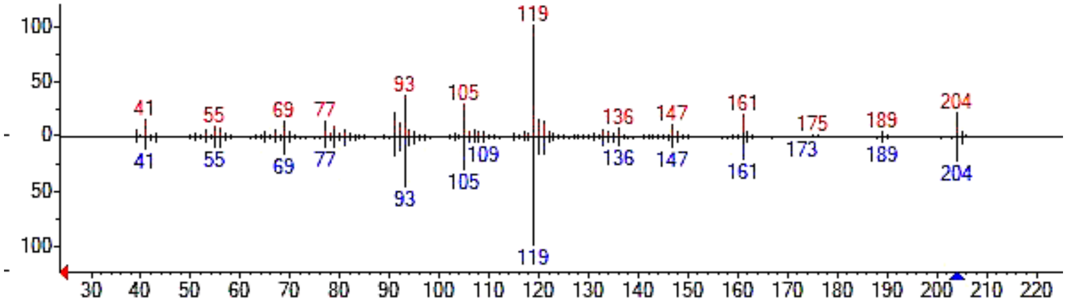 |

**Supplementary Table 11.** (Cont.)

| Compound Name | Molecular formula | Structure | Mass spectra |
|---|---|---|---|
| Cedrene | $C_{15}H_{24}$ | 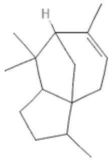 | 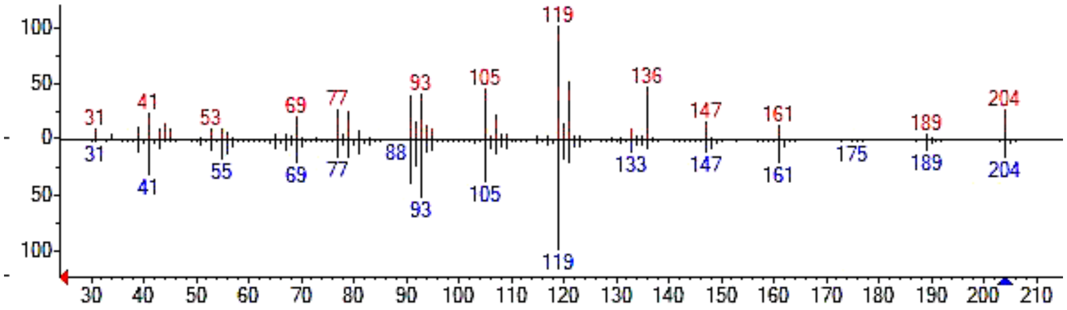 |
| Curcumene | $C_{15}H_{22}$ | 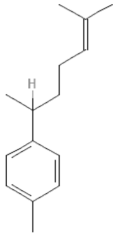 | 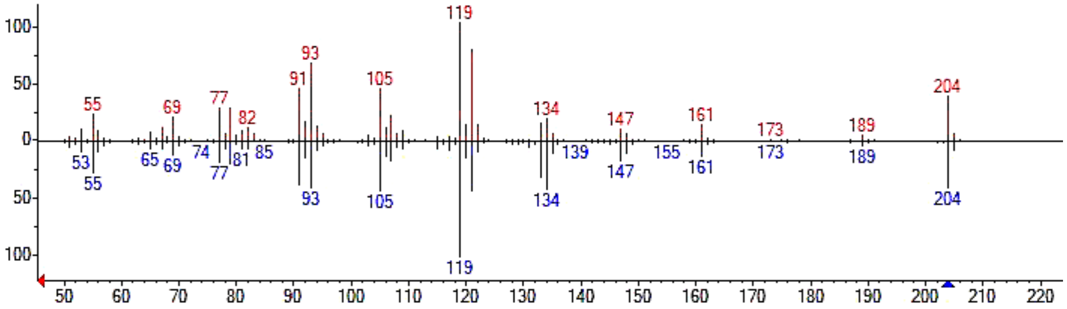 |
| Himalachene | $C_{15}H_{24}$ | 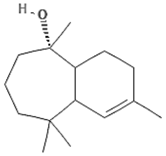 | 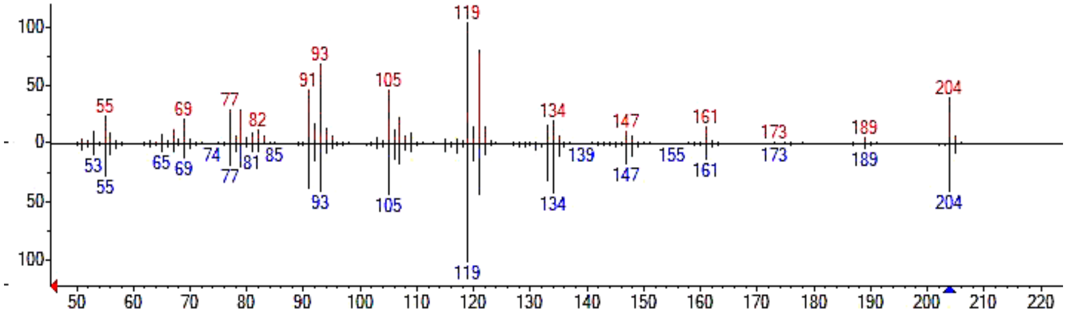 |

**Supplementary Table 11.** (Cont.)

| Compound Name | Molecular formula | Structure | Mass spectra |
|---|---|---|---|
| Isoledene | $C_{15}H_{24}$ | 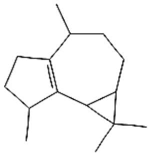 | 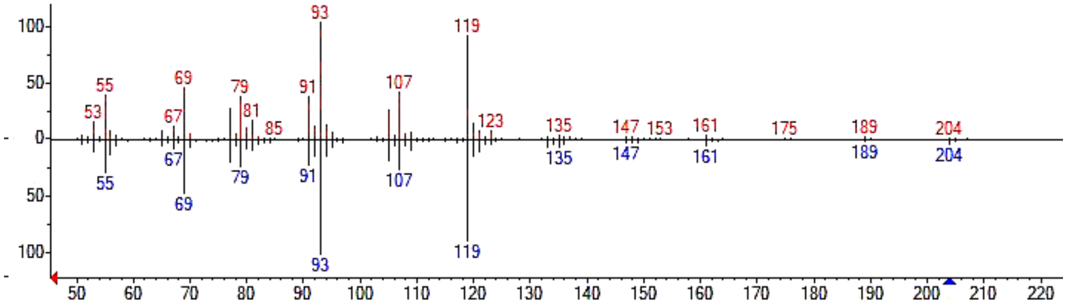 |
| Longipinene | $C_{15}H_{24}$ | 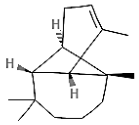 | 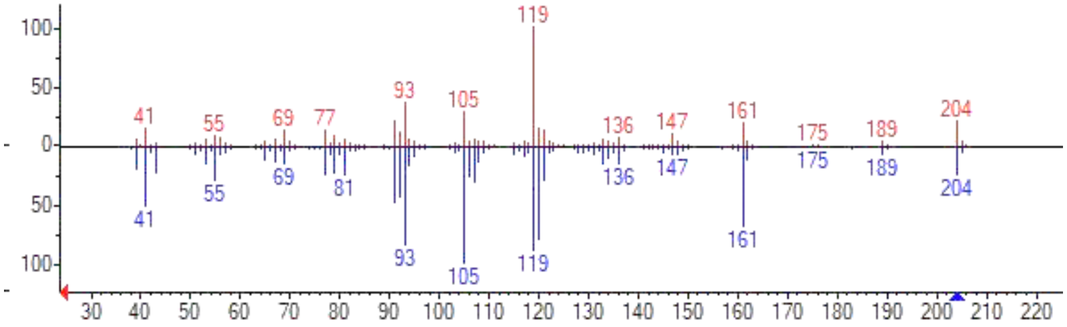 |
| Germacrene D | $C_{15}H_{24}$ | 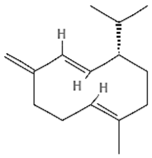 | 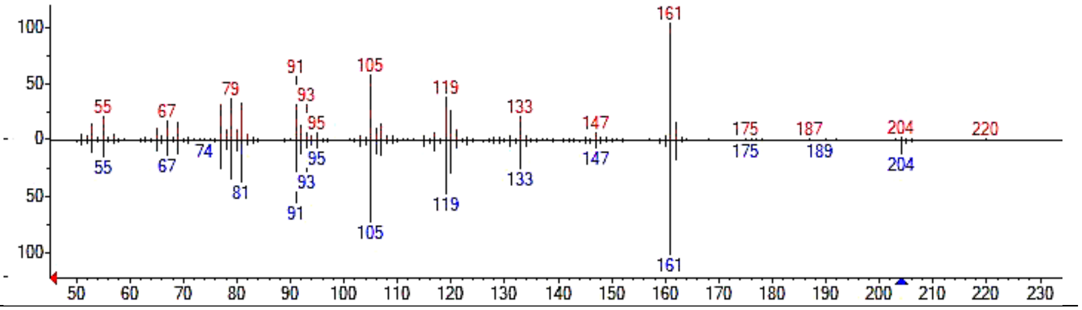 |

**Supplementary Table 11.** (Cont.)

| Compound Name | Molecular formula | Structure | Mass spectra |
|---|---|---|---|
| γ-Muurolene | $C_{15}H_{24}$ | 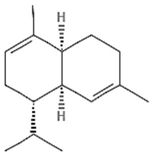 | 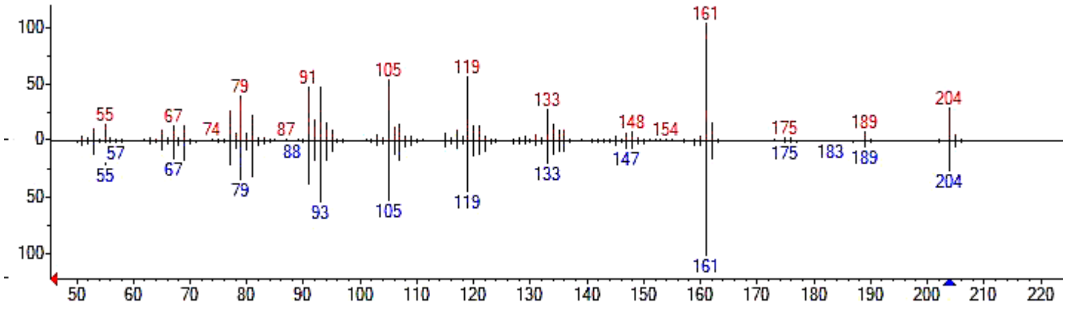 |
| Caryophyllene | $C_{15}H_{24}$ | 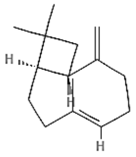 | 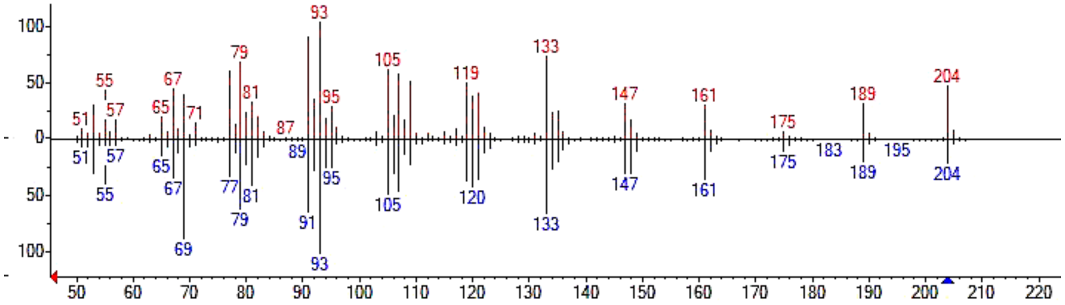 |
| β-Agarofuran | $C_{15}H_{26}O$ | 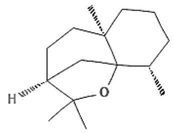 | 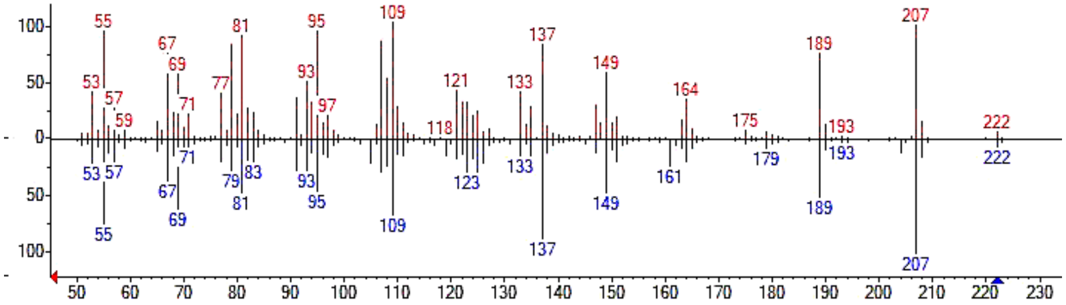 |
| α-Humulene | $C_{15}H_{24}$ | 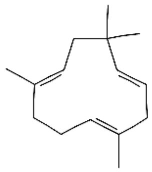 | 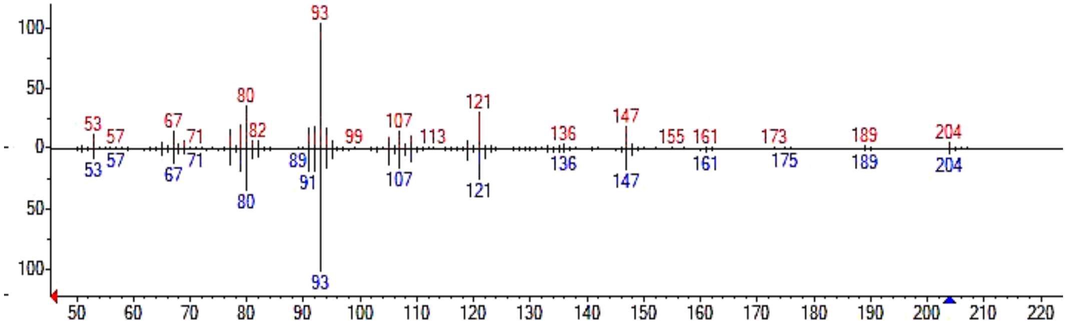 |

**Supplementary Table 11.** (Cont.)

| Compound Name | Molecular formula | Structure | Mass spectra |
|---|---|---|---|
| α-Guaiene | $C_{15}H_{24}$ | 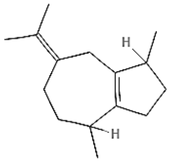 | 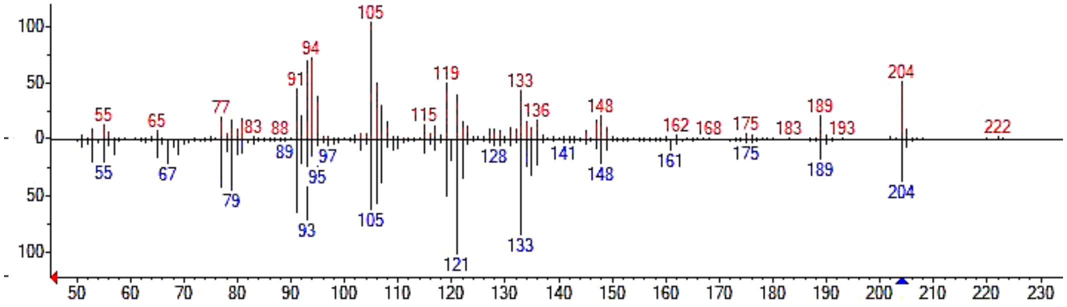  |
| Copaene | $C_{15}H_{24}$ | 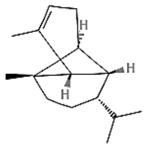 | 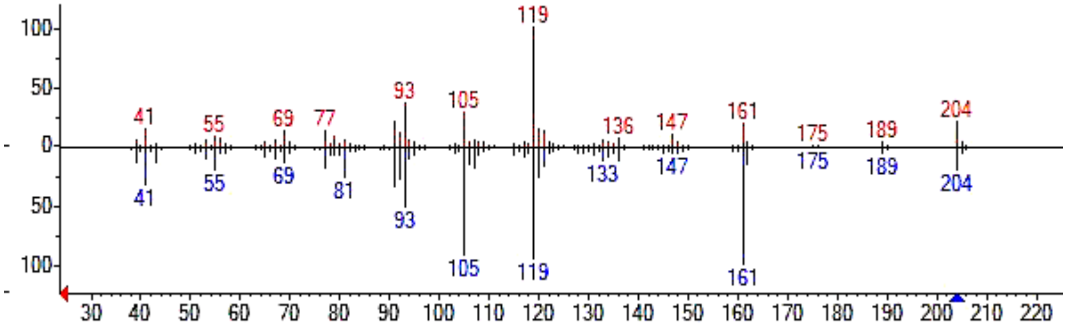  |
| Aromadendrene | $C_{15}H_{24}$ | 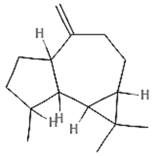 | 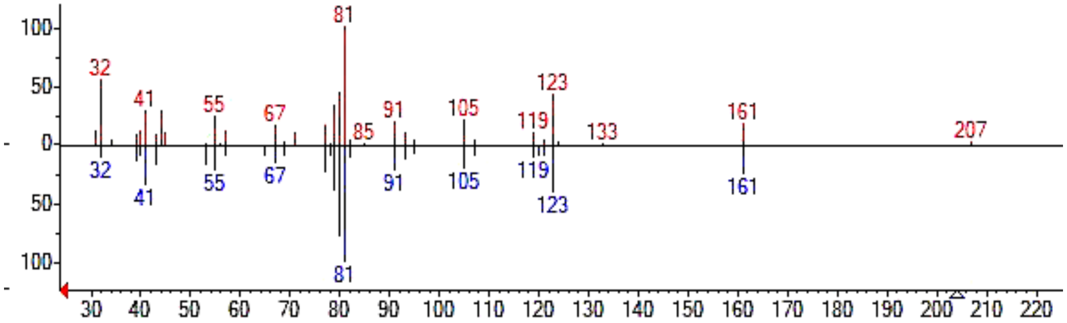  |

**Supplementary Table 11.** (Cont.)

| Compound Name | Molecular formula | Structure | Mass spectra |
|---|---|---|---|
| β-Bourbonene | $C_{15}H_{24}$ | 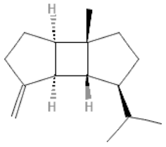 | 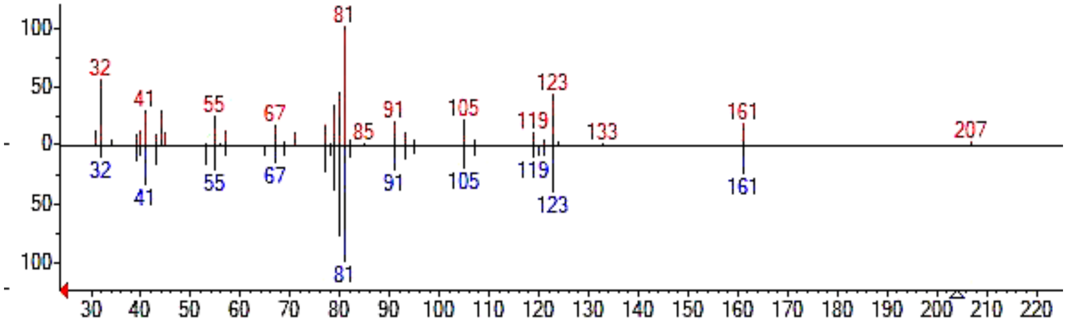  |
| α-Bourbonene | $C_{15}H_{24}$ | 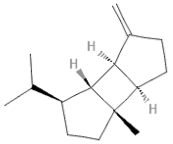 | 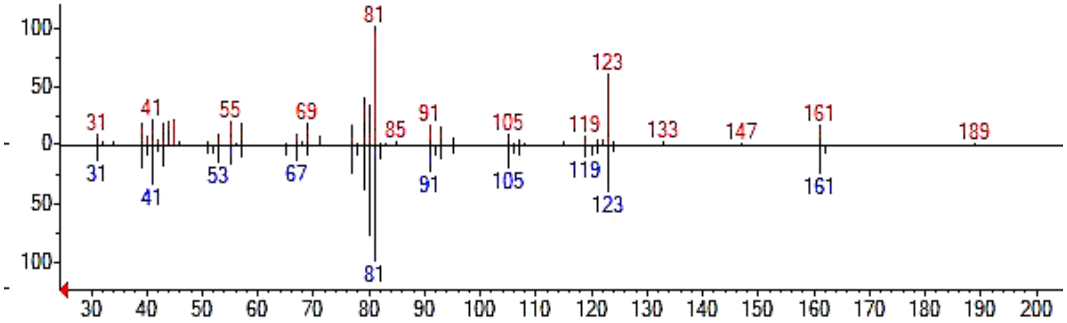  |
| Alloaromadendrene | $C_{15}H_{24}$ | 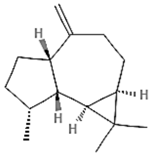 | 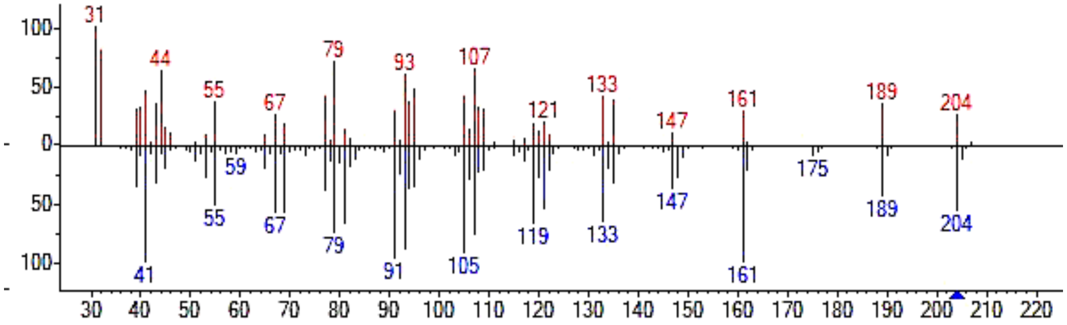  |

**Supplementary Table 11.** (Cont.)

| Compound Name | Molecular formula | Structure | Mass spectra |
|---|---|---|---|
| Seychellene | $C_{15}H_{24}$ | 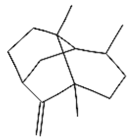 | 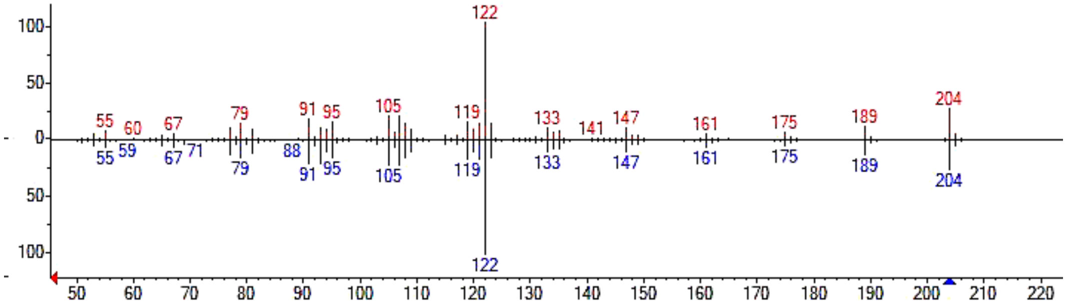 |
| Patchoulene | $C_{15}H_{24}$ | 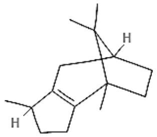 | 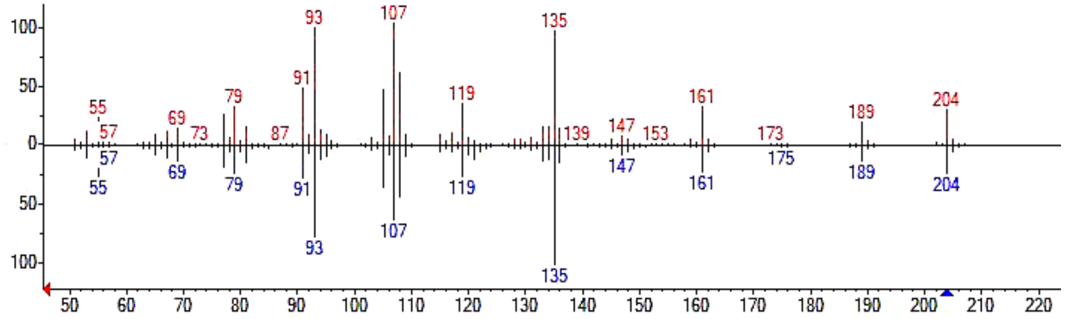 |
| Cadinol | $C_{15}H_{26}O$ | 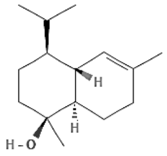 | 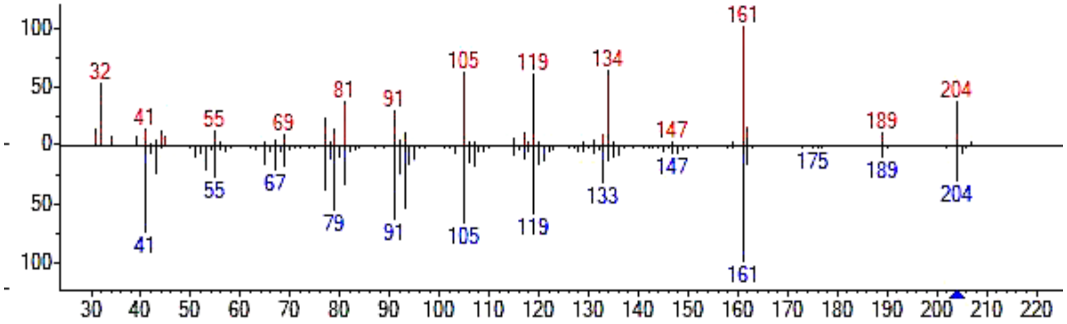 |

**Supplementary Table 11.** (Cont.)

| Compound Name | Molecular formula | Structure | Mass spectra |
|---|---|---|---|
| Selinene | $C_{15}H_{24}$ | 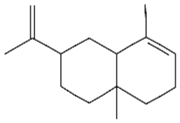 | 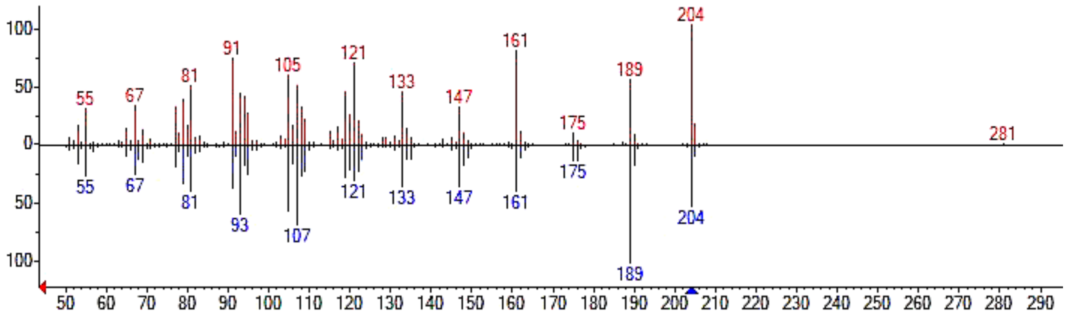 |
| Eremophilane | $C_{15}H_{24}$ | 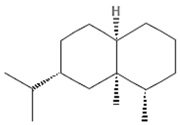 | 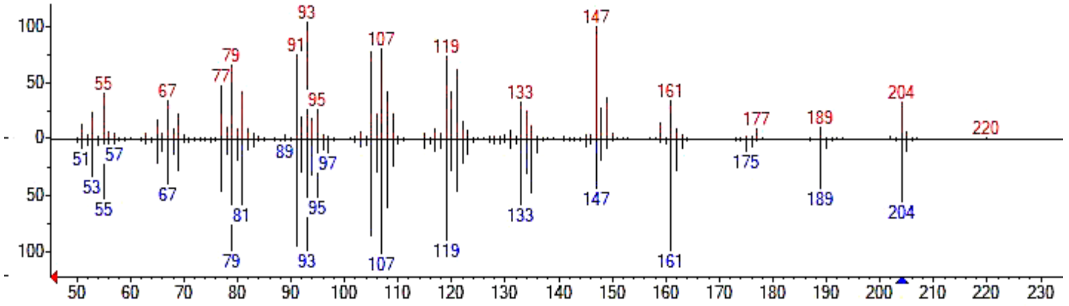 |
| δ-Guaiene | $C_{15}H_{24}$ | 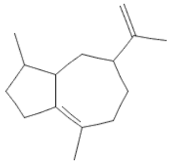 | 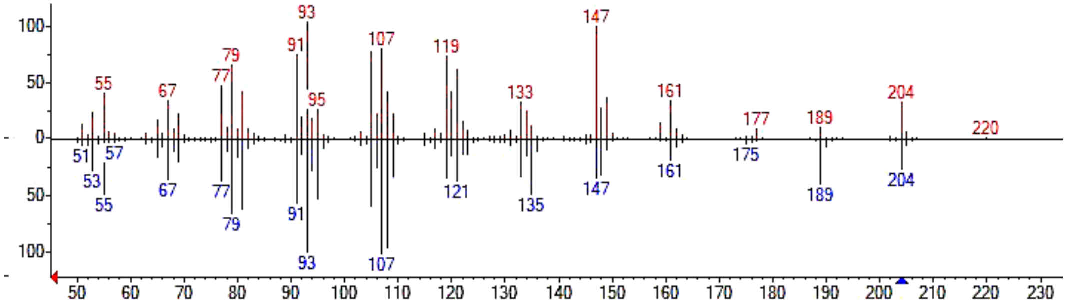 |

**Supplementary Table 11.** (Cont.)

| Compound Name | Molecular formula | Structure | Mass spectra |
|---|---|---|---|
| Aristolochene | $C_{15}H_{24}$ | 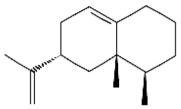 | 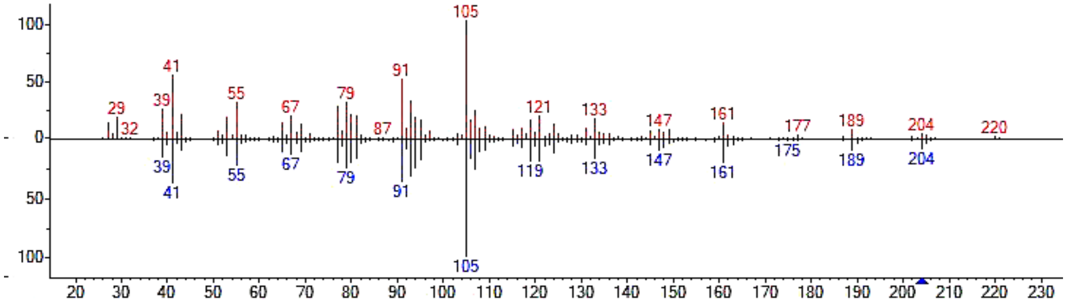 |
| Caryophyllene oxide | $C_{15}H_{24}O$ | 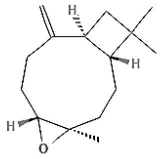 | 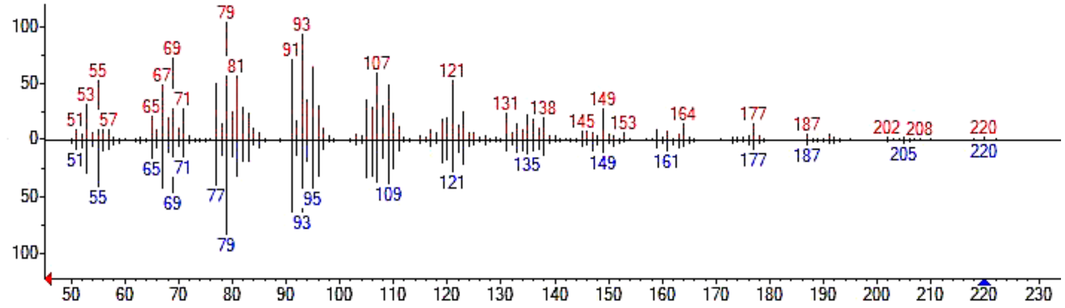 |
| Valerianol | $C_{15}H_{26}O$ | 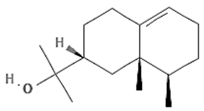 | 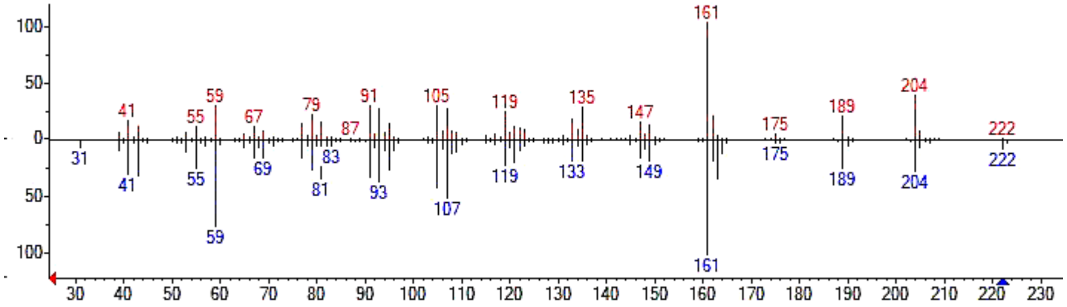 |
| Patchoulol | $C_{15}H_{26}O$ | 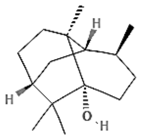 | 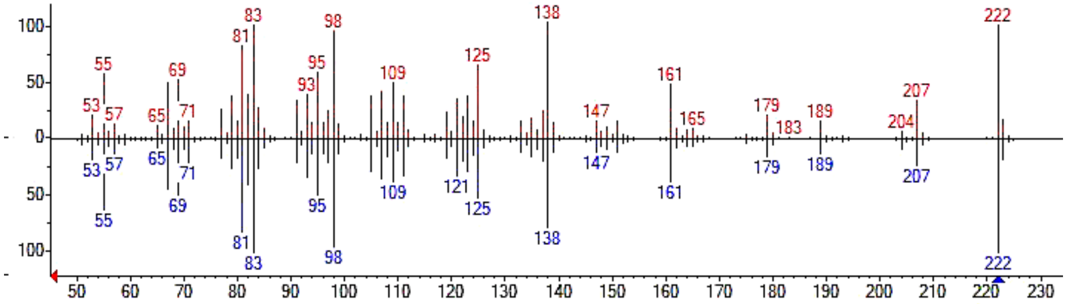 |

**Supplementary Table 11.** (Cont.)

| Compound Name | Molecular formula | Structure | Mass spectra |
|---|---|---|---|
| Patchoulene | $C_{15}H_{24}$ | 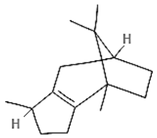 | 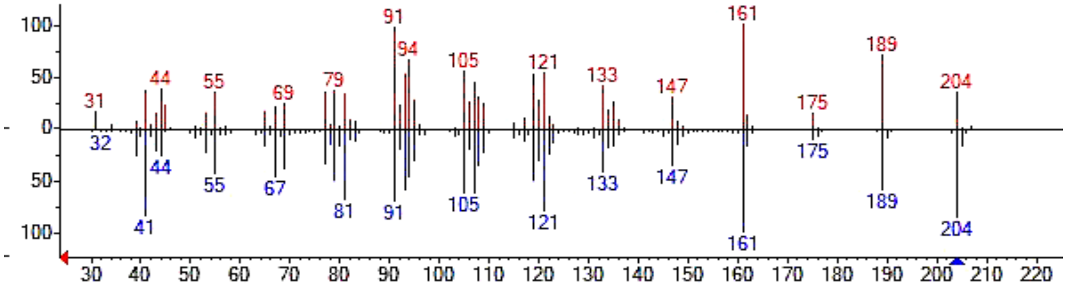 |
| Bisabolol | $C_{15}H_{26}O$ | 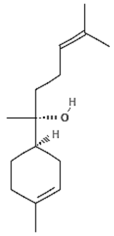 | 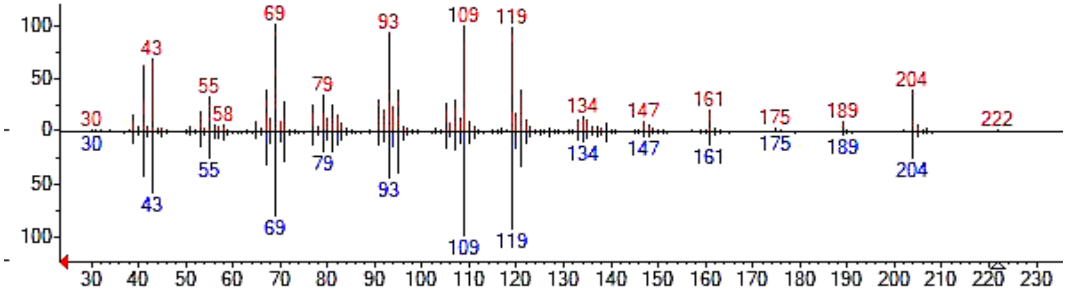 |
| Valencene | $C_{15}H_{24}$ | 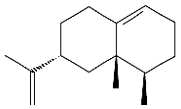 | 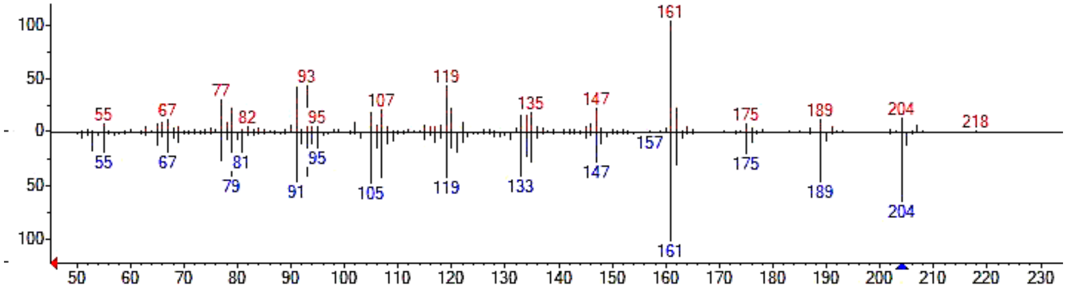 |
| Gurjunene | $C_{15}H_{24}$ | 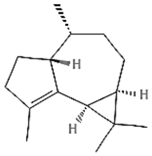 | 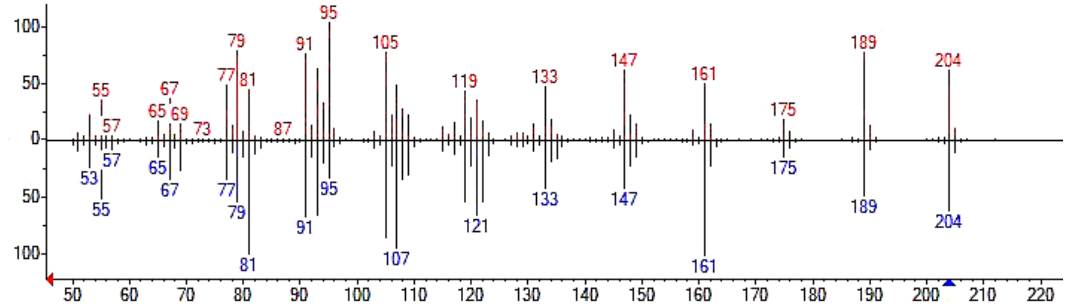 |

**Supplementary Table 11.** (Cont.)

| Compound Name | Molecular formula | Structure | Mass spectra |
|---|---|---|---|
| α-Santalol | $C_{15}H_{24}O$ | 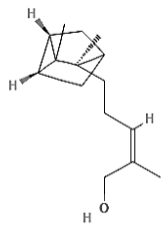 | 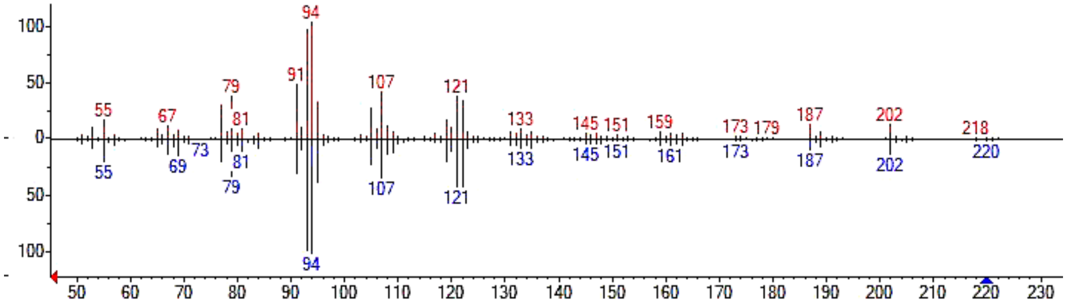  |
| β-Santalol | $C_{15}H_{24}O$ | 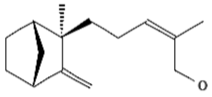 | 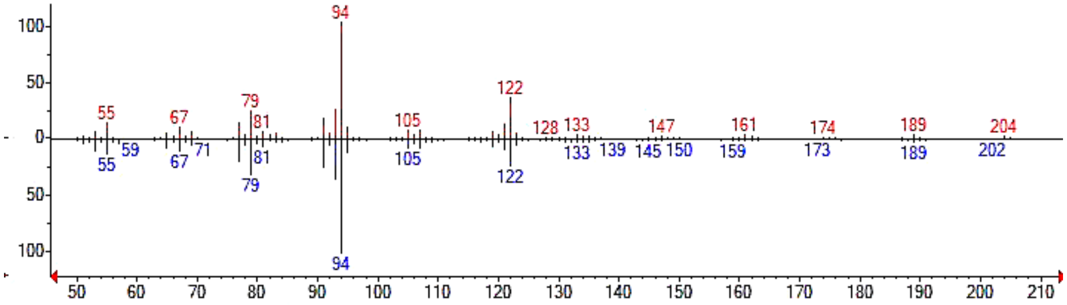  |
| Isolongifolol | $C_{16}H_{28}O$ | 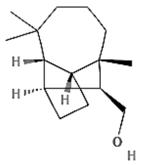 | 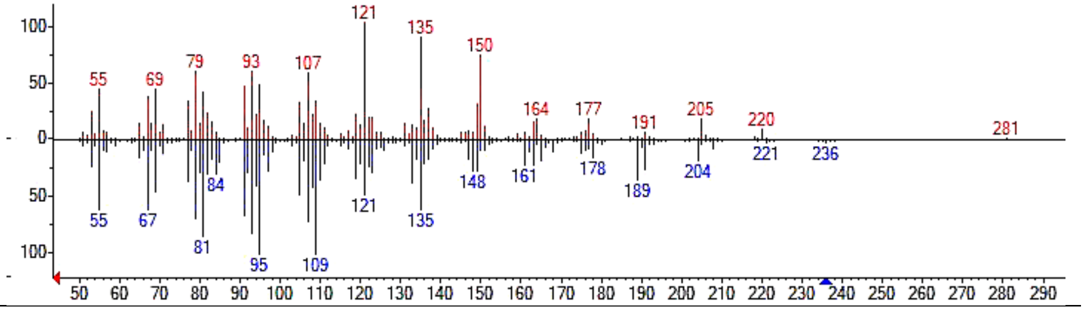  |

**Supplementary Table 11.** (Cont.)

| Compound Name | Molecular formula | Structure | Mass spectra |
|---|---|---|---|
| Cubenol | $C_{15}H_{26}O$ | 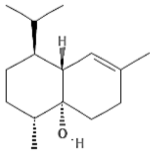  | 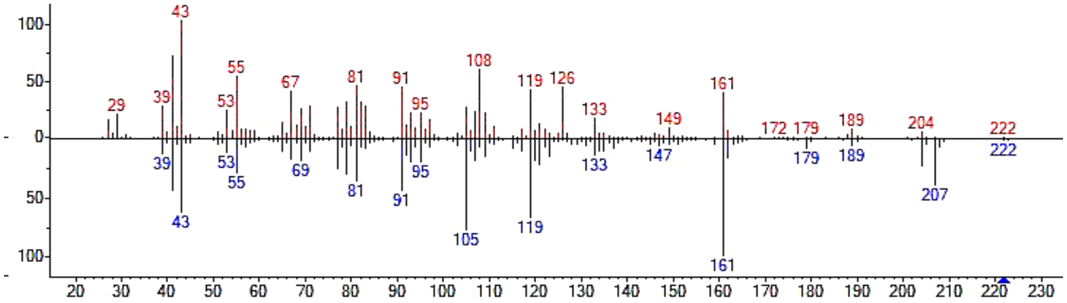  |
| Khusimol | $C_{15}H_{24}O$ | 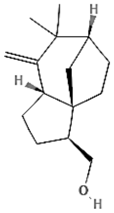  | 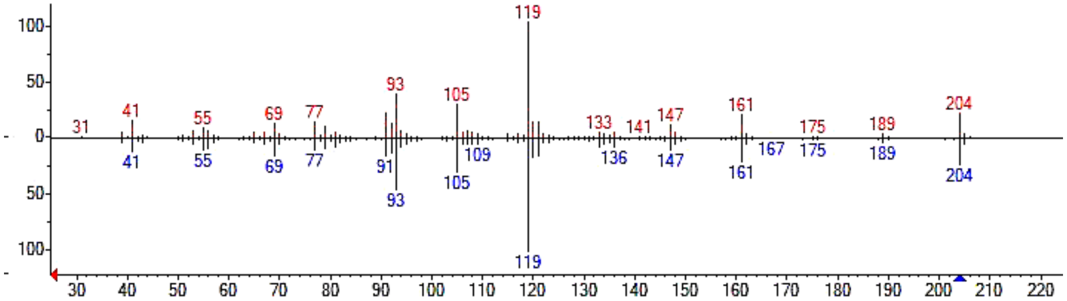  |
| Dihydrokaranone | $C_{15}H_{22}O$ | 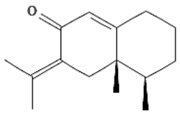  | 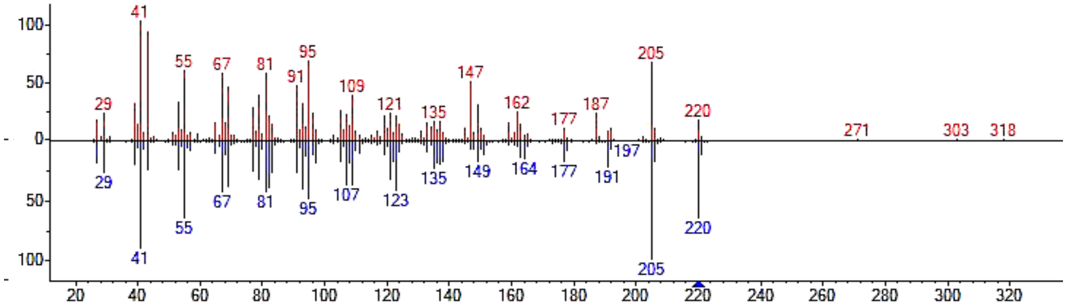  |
| Cadinene | $C_{15}H_{24}$ | 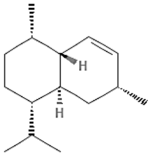  | 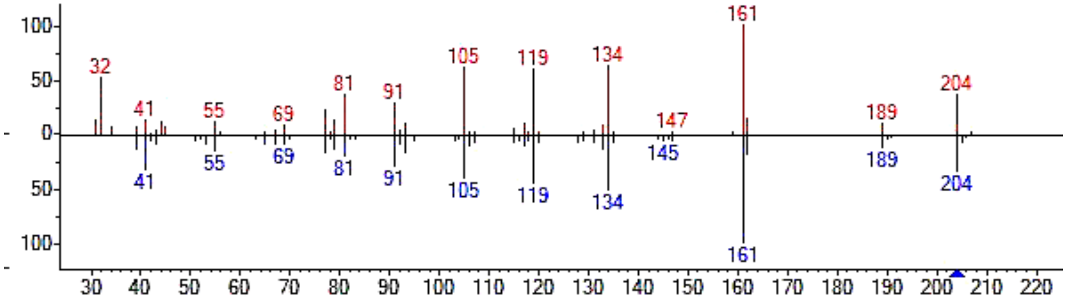  |

**Supplementary Table 11.** (Cont.)

| Compound Name | Molecular formula | Structure | Mass spectra |
|---|---|---|---|
| Guaiol | $C_{15}H_{26}O$ | 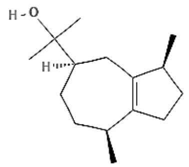 | 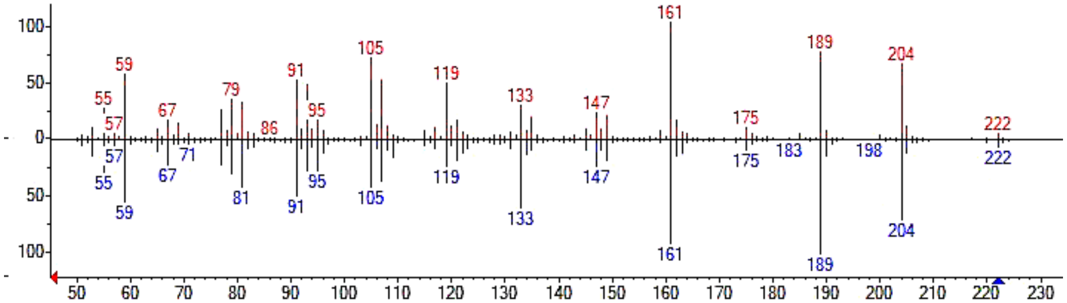 |
| Globulol | $C_{15}H_{26}O$ | 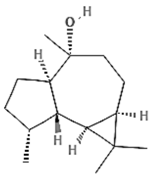 | 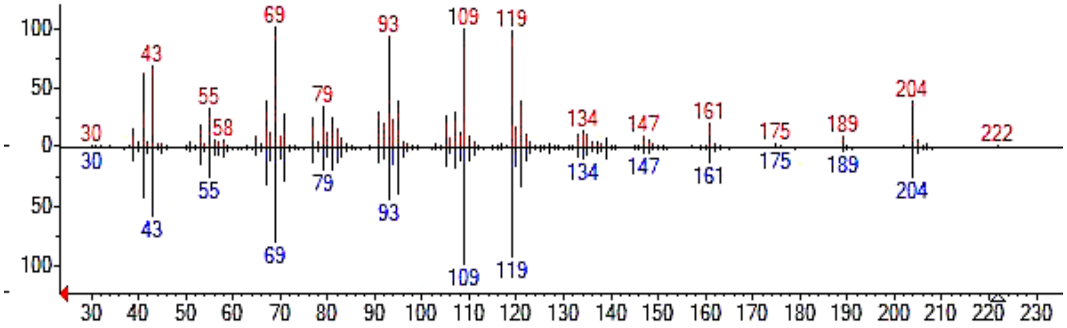 |
| Guaia-1(10),11-dien-15,2-olide | $C_{15}H_{20}O_2$ | 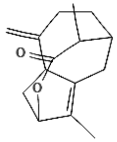 | 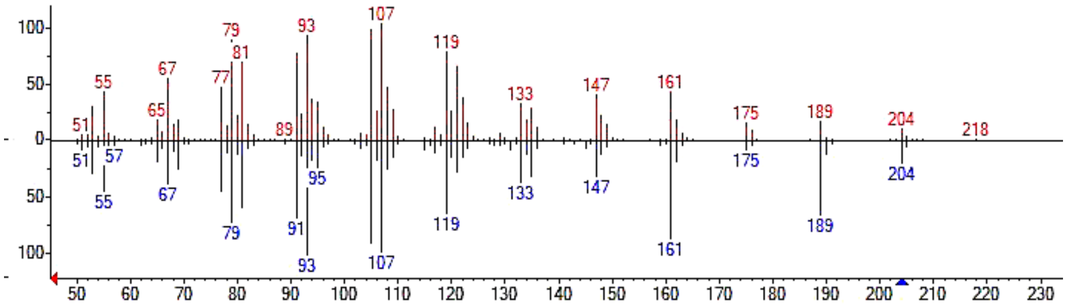 |
| Eudesmol | $C_{15}H_{26}0$ | 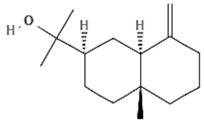 | 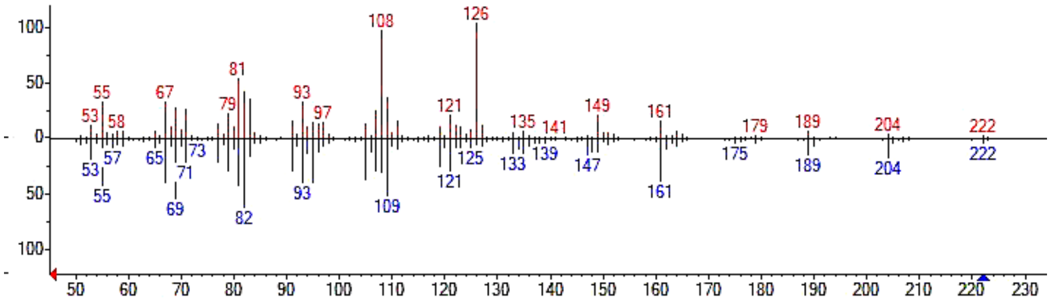 |

**Supplementary Table 11.** (Cont.)

| Compound Name | Molecular formula | Structure | Mass spectra |
|---|---|---|---|
| Jinkoheremol | $C_{15}H_{26}O$ | 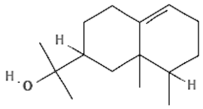 | 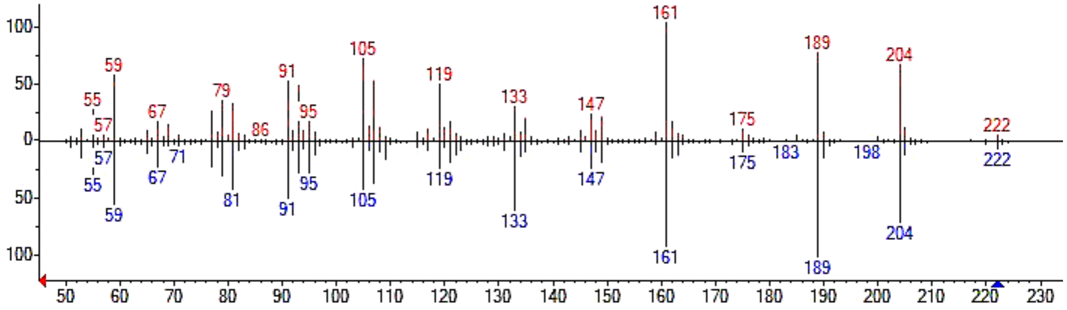  |
| Humulane-1,6-dien-3-ol | $C_{15}H_{26}O$ | 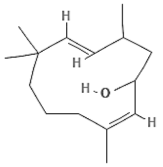 | 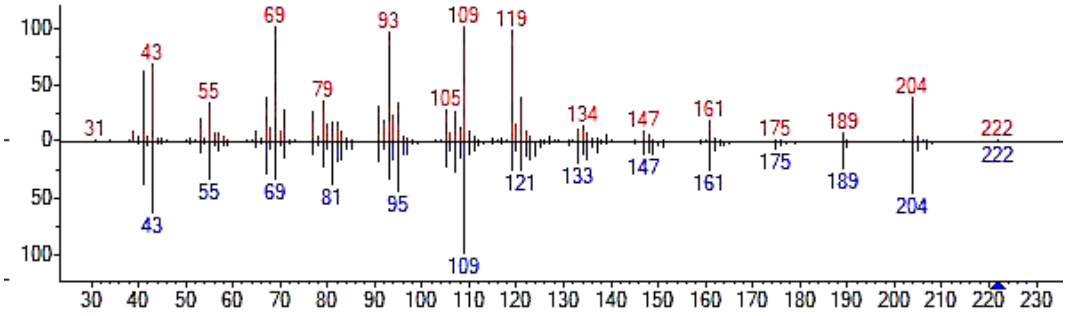  |
| Bisabolol oxide B | $C_{15}H_{26}O_2$ | 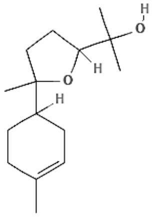 | 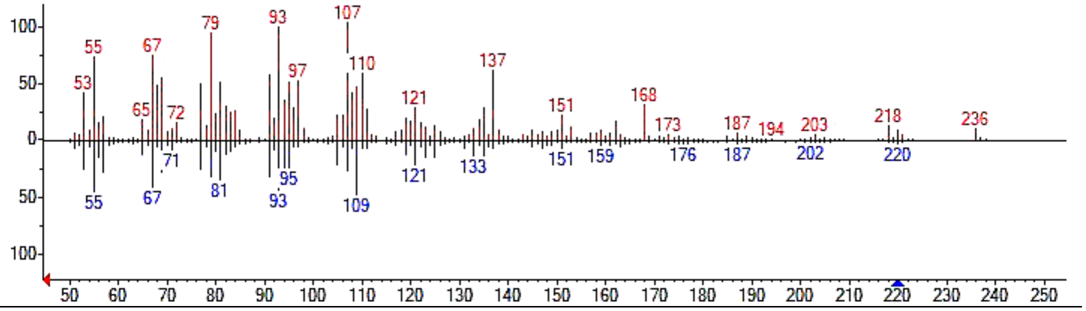  |

**Supplementary Table 11.** (Cont.)

| Compound Name | Molecular formula | Structure | Mass spectra |
|---|---|---|---|
| Bisabolol oxide A | $C_{15}H_{26}O_2$ | 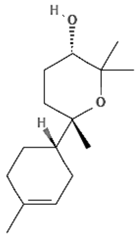 | 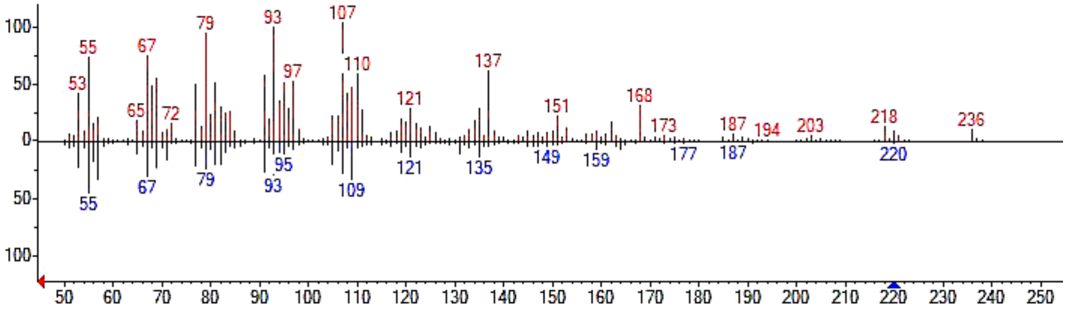 |
| Ylangene | $C_{15}H_{24}$ | 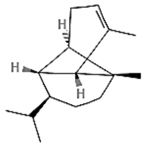 | 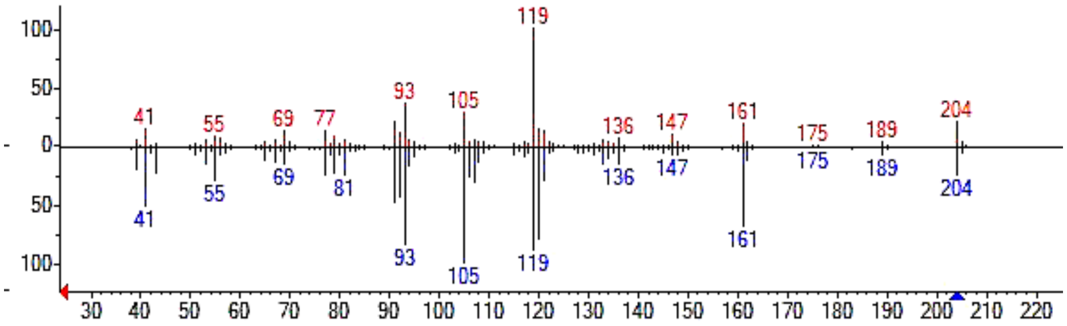 |
| Agidol | $C_{15}H_{24}O$ | 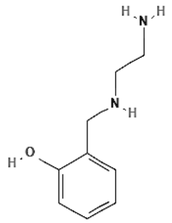 | 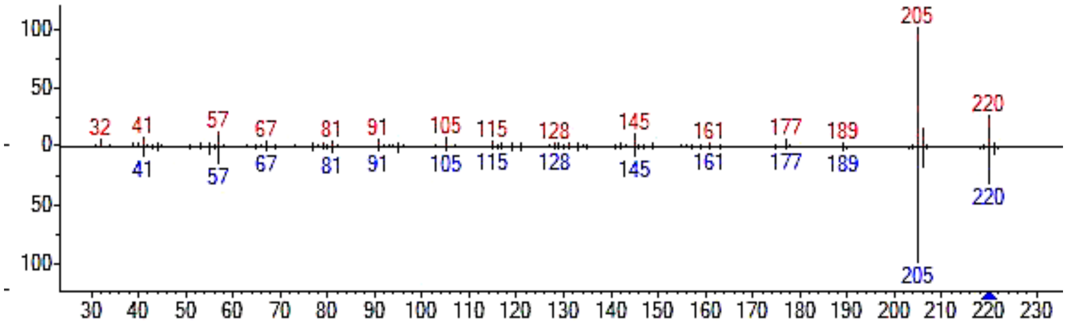 |

**Supplementary Table 11.** (Cont.)

| Compound Name | Molecular formula | Structure | Mass spectra |
|---|---|---|---|
| Ledol | $C_{15}H_{26}O$ | 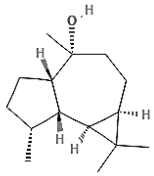 | 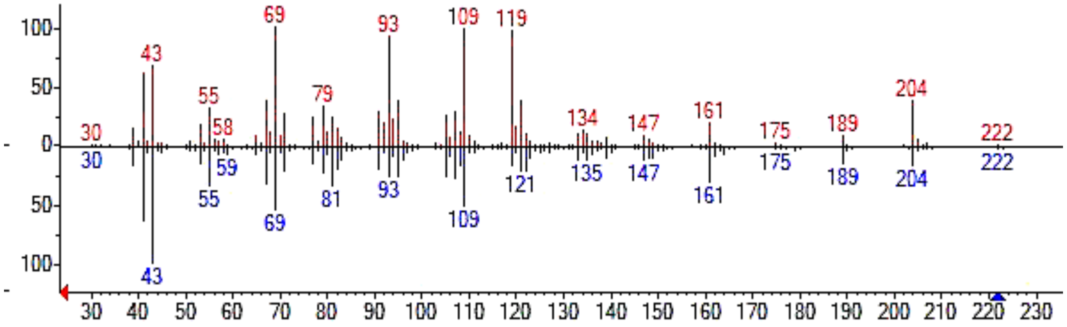  |
| Aromadendrene oxide | $C_{15}H_{24}O$ | 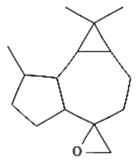 | 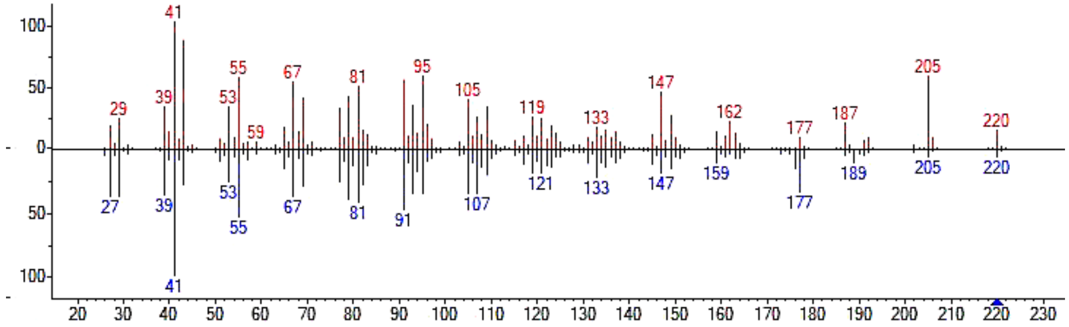  |
